# SESCA: Predicting Circular Dichroism Spectra from Protein Molecular Structures

**DOI:** 10.1101/279752

**Authors:** Gabor Nagy, Maxim Igaev, Søren V. Hoffmann, Nykola C. Jones, Helmut Grubmüller

**Affiliations:** Department of Theoretical and Computational Biophysics, Max Planck Institute for Biophysical Chemistry, Am Fassberg 11, D-37077 Göttingen, Germany; ISA, Department of Physics & Astronomy, Aarhus University, Ny Munkegade 120, DK 8000 Aarhus C, Denmark

**Keywords:** protein structure, CD spectrum prediction, structure validation, secondary structure

## Abstract

Circular dichroism spectroscopy is a highly sensitive, but low-resolution technique to study the structure of proteins. Combined with molecular modelling or other complementary techniques, CD spectroscopy can provide essential information at higher resolution. To this end, we introduce a new computational method to calculate the electronic circular dichroism spectra of proteins from a structural model or ensemble using the average secondary structure composition and a pre-calculated set of basis spectra. We compared the predictive power of our method to existing algorithms – namely DichroCalc and PDB2CD – and found that it predicts CD spectra more accurately, with a 50% smaller average deviation from the measured CD spectra. Our results indicate that the derived basis sets are robust to experimental errors in the reference spectra and to the choice of the secondary structure classification algorithm. For over 80% of the globular reference proteins, our basis sets accurately predict the experimental spectrum solely from their secondary structure composition. For the remaining 20%, correcting for intensity normalization considerably improves the prediction power. Additionally, we show that the predictions for short peptides and intrinsically disordered proteins strongly benefit from accounting for side-chain contributions and structural flexibility.

**Table Of Content Graphics:** 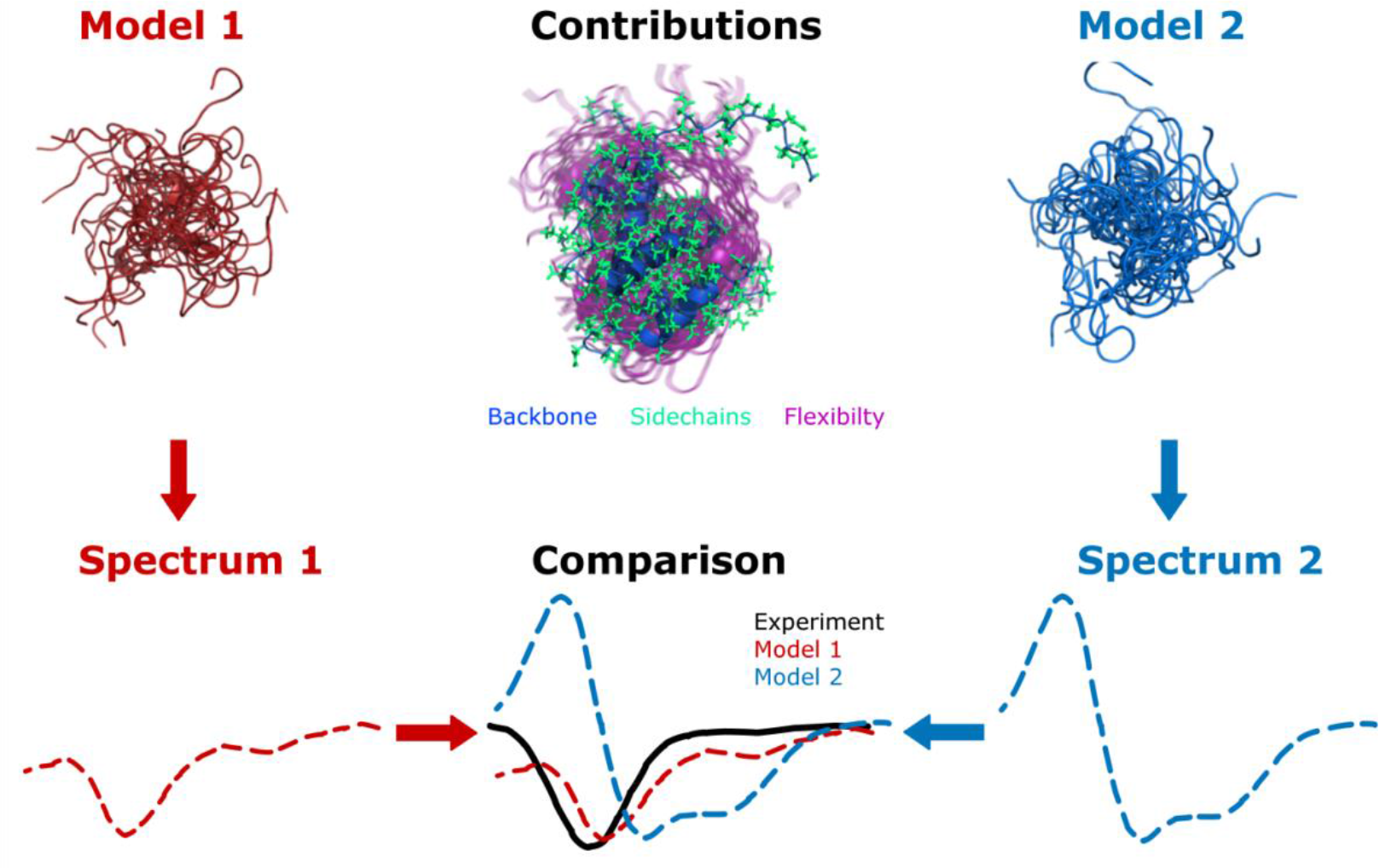

## Introduction

Electronic circular dichroism (CD) spectroscopy is a widely applied optical method to study the structure and structural changes of biomolecules such as proteins, nucleic acids, and carbohydrates ^1^. As a very sensitive tool, CD is often used as a quality control for recombinant proteins or to monitor changes of the protein structure during folding, aggregation, and binding events. Because of its sensitivity, CD spectroscopy does not require large amounts of protein or special labelling and can be readily used in aqueous solutions.

At a more quantitative level, the CD spectra of proteins in the far ultraviolet (UV) range (180-250 nm) provide structural information. The main contributor to a protein CD spectrum in this range is the absorption of partially delocalized peptide bonds of the backbone, such that the spectrum is mainly determined by the secondary structure (SS) ^2–5^. However, isolated amino acids, except glycine, also show a CD signal in this wavelength range ^6–8^. Thus, amino acid (AA) side chains also contribute to the protein CD spectrum, albeit to a smaller extent.

To extract information from CD spectra, it is essential to establish a quantitative link between structural models and the observed spectra. Since the 1980‘s, two major categories of methods have been established: spectrum deconvolution methods (Fig. 1A) predict the SS composition of a protein from its CD spectrum, whereas spectrum prediction methods (Fig. 1B-1D), *vice versa*, determine the CD spectrum from a structure.

**Figure 1:**
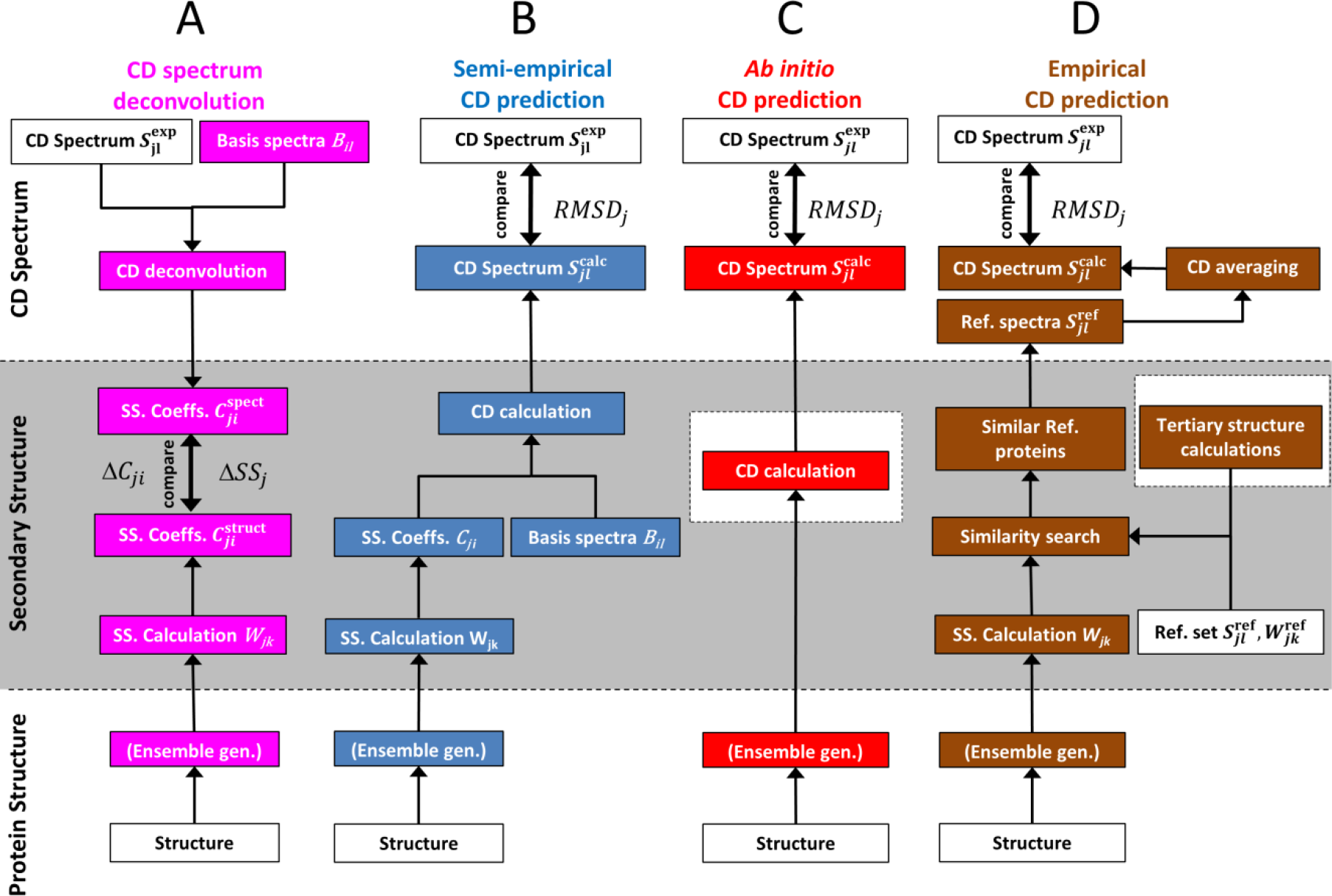
Relating a protein structure to its circular dichroism spectrum. White rectangles represent experimental data, magenta, blue, red, and brown fields are related to spectrum deconvolution, semi-empirical, *ab initio*, and empirical spectrum calculations, respectively. For spectrum deconvolution (column A), the secondary structure is determined independently both from the CD spectrum and from the structural model, and subsequently compared. In contrast, semi-empirical (column B), *ab initio* (column C), and empirical (column D) spectrum prediction methods compute a CD spectrum from the structure, that is compared to the measured spectrum. Fields in the shaded area denote calculation steps at the secondary structure level.

Spectrum deconvolution methods (e.g., CCA, K2D3, BestSel)^9–11^ estimate the SS composition of proteins with unknown structure, based on the observation that proteins with different SS have different CD spectra. Accordingly, the measured CD spectrum is approximated by a linear combination of pure SS spectra or, more generally, properly chosen basis spectra. The coefficients obtained by this approximation provide the fractions of SS elements in the target protein (top-down arrows in Fig. 1A). Basis spectra have been previously derived either from the CD spectra of model peptides or from a larger set of reference proteins with known CD spectra and SS composition.^10^

The obtained SS information can also be used to validate structural models. To this end, SS estimates from CD deconvolution are compared to the SS composition of a proposed model (Fig. 1A bottom). However, as the fitting procedure is unstable, small experimental errors of the measured CD spectrum may already induce considerable uncertainties in the SS estimates, which renders the comparison to model structures problematic. Therefore, for model validation, spectrum prediction methods are preferred.

The most advanced CD spectrum prediction methods are *ab initio* (Fig. 1C) and typically require computationally demanding exited-state quantum mechanics (QM) or density functional calculations ^12–14^. These methods determine the light absorption of molecules directly from their structure, but the large computational effort usually limits such calculations to rather small peptides. To speed up calculations, a simplified *ab initio* spectrum prediction algorithm, called the matrix method ^15^, has been implemented in the program DichroCalc ^16^. DichroCalc determines the most important features of the CD spectrum of a protein based on its average crystallographic structure and parameters derived from *ab initio* QM calculations, albeit with limited accuracy ^17^.

In contrast, and as an alternative to *ab initio* spectrum predictions, the recently proposed PDB2CD ^17^ method (Fig. 1D) estimates the CD spectrum of a target protein from known CD spectra of structurally similar proteins selected from a reference set. By substituting the computationally demanding QM calculations with a secondary and tertiary structure based similarity search among the reference proteins, PDB2CD achieves markedly higher average prediction accuracy than DichroCalc. However, this accuracy is limited by the number of reference proteins structurally similar to the target. Thus, PDB2CD may be less accurate for proteins with an unusual fold or high levels of disorder. Additionally, CD spectra yielded by PDB2CD are calculated from only a few reference spectra, and therefore, its predictions are prone to the experimental error of those spectra, decreasing the robustness of the method.

Here, we develop and test a spectrum prediction method for the efficient validation and refinement of protein structures against measured CD spectra. Our Semi-Empirical Spectrum Calculation Approach (SESCA) avoids the disadvantages of the above methods by combining elements of both deconvolution and spectrum prediction. Specifically, like deconvolution methods, it uses structure-related basis spectra, but avoids a potentially unstable fit by obtaining the required coefficients directly from a model structure (Fig 1B), similarly to other bottom-up CD prediction methods (Fig. 1C-1D).

Using basis spectra for protein CD predictions provides several advantages over the previously available methods to allow fast, yet accurate spectrum predictions. Firstly, SESCA basis sets combine all the experimentally determined structural and spectral information to describe the average CD signals for the local conformations of peptide bonds (‘SS classes’). Because our method is based on statistics of peptide bond conformations rather than on identifying similar tertiary structures, we expect SESCA to have larger predictive power than PDB2CD, particularly for proteins with structures dissimilar from those in the reference set, or even with no structure at all. Additionally, SESCA is more robust to experimental noise of the individual CD spectra than PDB2CD, because our basis spectra are averaged over the complete reference set, whereas PDB2CD uses only a small fraction for each prediction.

Secondly, extracting pre-calculated basis spectra from the available reference proteins reduces the computational complexity of CD predictions to the calculation of a single linear combination. Further, to determine basis spectrum coefficients SESCA only requires the SS composition extracted from the model structure. Consequently, it avoids both the costly chromophore calculations of DichroCalc, as well as the computational overhead from the fold-recognition and the similarity search used in PDB2CD. Finally, by taking advantage of the existing high-throughput SS classification methods, SESCA allows CD predictions from large structural ensembles that currently neither DichroCalc nor PDB2CD can perform.

In this study, our approach is evaluated and optimized using various available SS classification algorithms. Additionally, we address (a) the effects of conformational flexibility by using structural ensembles during CD predictions, and (b) the contribution of natural amino acid side chains. We show that including these contributions increases the prediction accuracy particularly for peptides and an intrinsically disordered protein (IDP) complex, and thus should also enable the refinement of IDP ensemble models against measured CD spectra.

## Theoretical background

### 2.1 Basis spectrum calculations

We will initially assume that the CD spectra are mainly determined by the local conformation of the peptide bonds, and subsequently also consider the effects of AA side chain groups within the same framework. The workflow of our semi-empirical CD prediction method is described in Fig. 2. Firstly, the local backbone conformation is grouped into secondary structure elements with established methods (Fig. 2A), to obtain the SS information from the protein structure required for the SESCA calculations. Secondly, these SS elements are combined into broader classes (Fig. 2B) for which basis spectra are determined (Fig. 2C). This grouping is necessary to control the number of basis spectra and optimize the prediction accuracy. The predicted CD spectrum of the protein is calculated from weighted averages of basis spectra (Fig. 2D), which is compared to the measured CD spectrum to determine the quality of the model structure. Note that besides the SS composition obtained from the model, the parameters SESCA predictions require are stored in a collection of basis spectra, and the assignment matrix used for grouping the SS elements. Henceforth, we refer to these parameters as a basis set.

**Figure 2:**
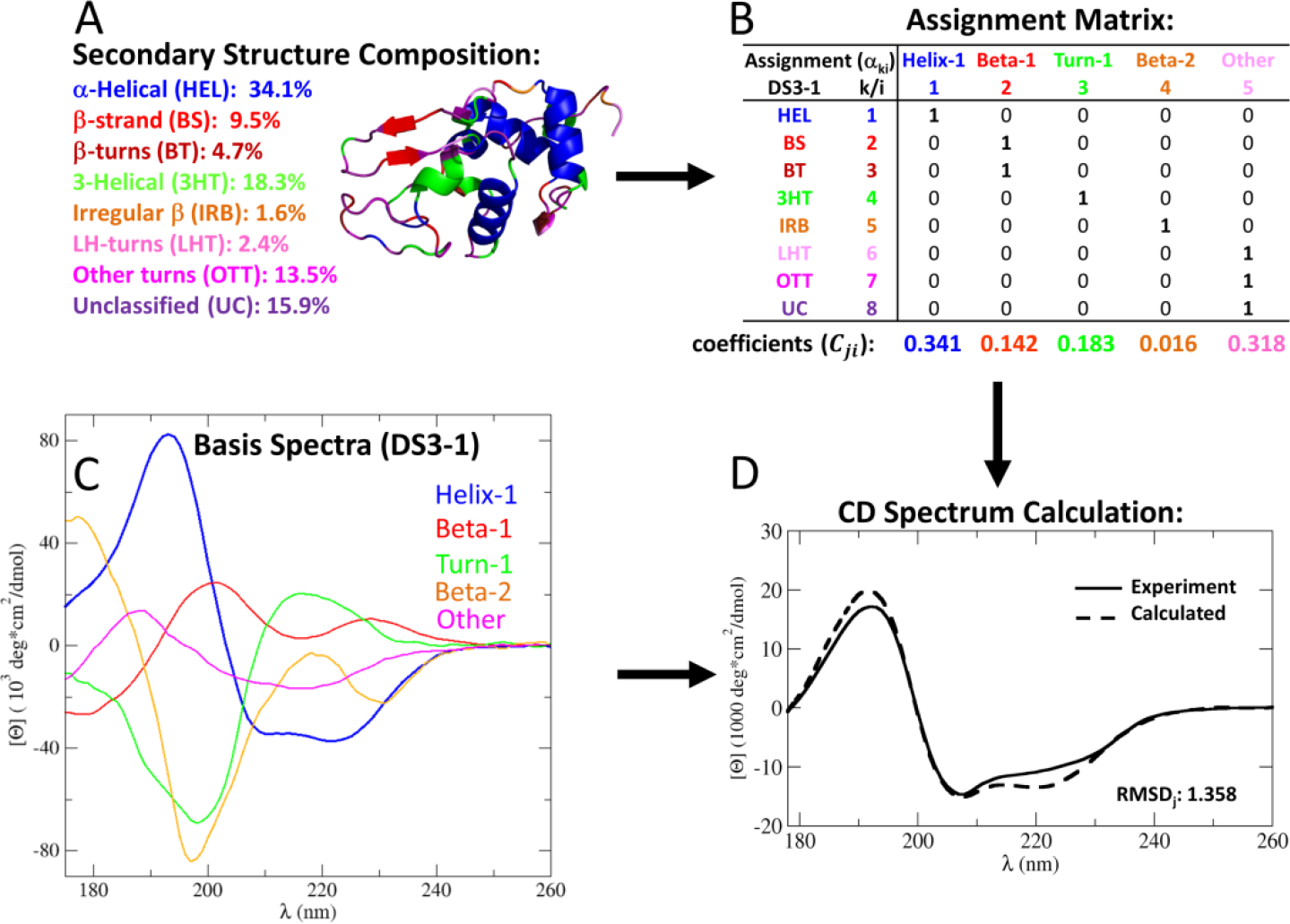
Semi-empirical CD spectrum calculation scheme. A) Secondary structure composition (in colors) and a cartoon representation of a protein *j* (here, lysozyme). The secondary structure information is translated into a CD spectrum via a basis set (here, DS3-1) comprised of an assignment matrix and a set of basis spectra. B) The assignment matrix *α*_*ki*_ groups the secondary structure elements *k* (on the left) into secondary structure classes *i* (on the top). The secondary structure class composition (C_*ji*_ bottom line) determines the coefficients in the weighted average of the basis spectra in C), that are used to calculate the CD spectrum for the protein. D) The calculated CD spectrum (dashed line) is compared to the measured spectrum (solid line) via a root mean squared deviation RMSD_j_. Throughout, mean residue ellipticity units are used for the spectra.

We derive SESCA basis sets from a set of *N* globular reference proteins with known structures and CD spectra 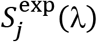. The latter are approximated by a weighted sum of *F* basis spectra *B*_*i*_(*λ*)

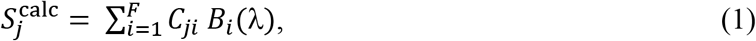

where the weights (coefficients) *C*_*ji*_ are the fractions of AAs in protein *j* that were assigned to the SS class *i*. We derive the basis spectra by minimizing the average root-mean-squared deviation (RMSD) between the measured spectra of the reference proteins 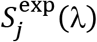 and those calculated from the secondary structure 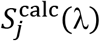 for all reference proteins,

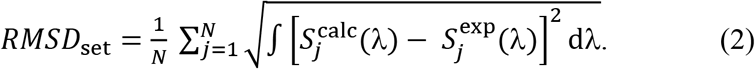

as described in the supplementary materials (SM) Section S1.

We note that in spectrum deconvolution methods ^9,10,18^ basis spectra are derived via the same notion, but are applied differently. In deconvolution, the basis spectrum coefficients (*C_ji_*) are treated as fit parameters which yield the SS content from the known CD spectrum of a target protein (Fig.1A). In our approach, the SS content is extracted from the known structure and combined into the basis spectrum coefficients to predict the CD spectrum. *C*_*ji*_ are calculated from the fraction of residues (***W***_*jk*_,) classified as SS element *k* in the structural model of protein *j* via an assignment matrix **A**={*α*_*ki*_} (Fig. 2B) such that

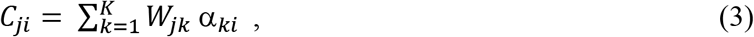

and the SS information described by the *K* secondary structure elements is now contained in *F* structural classes. Combining equations 1 and 3 relates the CD spectrum of a protein to its secondary structure composition

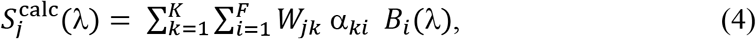

in a way, that for a given **A** matrix, the optimal basis spectra *B*_*i*_(*λ*) are readily calculated from the reference proteins by minimizing *RMSD*_set_. The assignment factors *α*_*ki*_ determine the average backbone structure of SS classes, the number and shape of the optimal basis spectra, and consequently strongly influence the predictive power of the basis set.

### 2.2 Assignment optimization

To find the most predictive basis sets, the number of basis spectra (basis set size) and the assignment of the SS elements are optimized. To this end, a Monte-Carlo (MC) search is performed among the possible **A** matrices, where both *α*_*ki*_ and 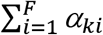 are limited to {0,1}. When basis sets are optimized under these constraints – henceforth referred to as ‘hard basis sets’ and ‘hard optimization’ – the procedure results in orthogonal SS classes and normalized basis spectra. For every resulting assignment a subset of the reference proteins (training set) is used to calculate the ideal basis spectra as described in SM Section S1, which are then used to predict the CD spectra of remaining reference proteins (evaluation set). The average deviation of the predicted CD spectra (RMSD_set_) is used to score the predictive power of the assignment during the optimization. Finally, the basis spectra of the best assignments are recalculated using all the reference proteins to acquire the final optimized basis set. Further details on the hard optimization procedure are provided in SM Section S2.

### 2.3 Including side-chain contributions

To assess the contribution of AA side chains, we assume that there are two main contributors to the CD spectra of proteins: the SS-dependent signal of the peptide bonds and the chromophores of the amino acid side chains, with no coupling between the side chains and the rest of the protein. This assumption allows us to represent the average contribution of side chain groups by additional basis spectra and the calculation of a backbone-independent side chain baseline. The baseline for each protein is determined by the weighted average of the individual side-chain basis spectra, where the weighing factor is the corresponding AA content extracted from the protein sequence (similarly to eq. 1). Details of the calculation of side-chain contributions are provided in SM Section S3. Note that the results we present in Section 4 are based purely on the SS information of the protein, and the effects of side-chain corrections are discussed in Section 5.3.

### 2.4 Basis set quality assessment

The quality of the optimized basis sets is assessed as follows. First, we determine the predictive power of each basis set by cross-validation. The prediction accuracy is quantified by computing RMSD_set_ for a set of proteins that is not used during the basis set determination (henceforth: prediction accuracy or RMSD_cross_). Secondly, we calculate RMSD_set_ for CD spectra of the reference protein set from which the basis sets are derived (henceforth fitting accuracy or RMSD_fit_). The difference between RMSD_fit_ and RMSD_cross_ for a basis set quantifies overfitting during the optimization. For basis sets with no overfitting, RMSD_fit_ and RMSD_cross_ should be similar, whilst for a basis set or spectrum prediction method with significant overfitting, RMSD_fit_ should be significantly lower than RMSD_cross_.

We also perform additional analyses to determine the limits of basis set accuracy using the applied reference protein set. To this end, we derive series of specialized basis sets with 1 to *F* basis spectra (where *F* is the number of the SS elements in the classification protocol used). These basis sets are derived by a different, unconstrained optimization scheme aimed solely to minimize RMSD_fit_, with no regard to overfitting or predictive power. Details about this optimization approach (‘soft optimization’) are provided in SM Section S4. The ‘soft basis sets’ serve as reference points on how accurately the CD spectra of the reference protein set can be described using the limited information encoded in the SS composition of the protein.

Finally, we perform a principal component analysis (PCA) on the CD spectra of the reference protein set we derived our basis sets from. Constructing basis sets from the principle component (PC) vectors of the CD spectra allows us to estimate the upper limit of prediction accuracy for globular proteins using a given basis set size. Details of this analysis are described in SM Section S5. This upper accuracy limit is used to estimate how much could we improve our prediction algorithm by including extra structural information in addition to the SS composition.

### 2.5 Structure validation

Once derived, SESCA basis sets can be used to validate protein structural models based on the measured CD spectrum of the protein. Specifically, one can estimate the total error in the SS composition of a structural model (*ΔSS*_*j*_) for protein *j*, based on the average deviation (*RMSD*_*j*_) between its predicted and experimentally measured CD spectrum. Note that even for the correct SS composition 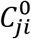 (i.e., *ΔSS*_*j*_ = 0), the CD spectrum predicted at wavelength *l* from the model would still deviate from the measured spectrum by 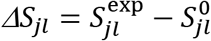, due to the approximations in our prediction method (Section 2.1), as well as due to the experimental error of the measured CD spectrum. To accurately determine *ΔSS*_*j*_ for our model, one thus has to separate the SS-dependent error (*M*_*j*_) in the predicted spectrum – introduced by the structural model – from the SS-independent error 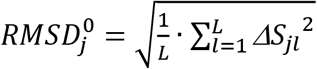, which describes the deviation between the experimental spectrum and the spectrum predicted from 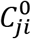.

To this end, we consider predicting the CD spectrum from an imperfect structural model with SS composition *C*_*ji*_, which deviates from the correct SS composition by 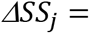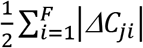, where 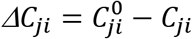is the error in the coefficient of SS class *i* for protein *j*. In Section S6, we show that the SS-dependent error for a given *ΔC*_*ji*_/*ΔSS*_*j*_ ratio is proportional to *ΔSS*_*j*_(*M*_*j*_ = *m*_*j*_ · *ΔSS*_*j*_) with a slope *m*_*j*_ that depends on the basis spectra as well as on the *ΔC*_*ji*_ ratios. Assuming that the SS-dependent and SS-independent errors at different wavelengths are statistically independent, the RMSD of the predicted spectrum is

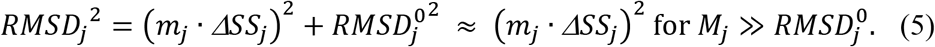

Even if the two error terms are correlated, the average error of the predicted spectra may change between 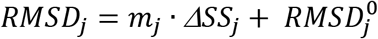 and 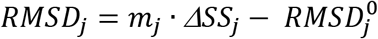. We note that if the CD spectra are free of systematic errors, 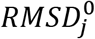 describes the average noise of the CD measurements, and should be statistically independent of ***M***_*j*_.

## Materials and Methods

### 3.1 Protein reference sets for calibration

To derive and assess the basis sets required for SESCA, 102 reference proteins were collected for which both the CD spectrum and the structure have been determined. These were grouped into non-overlapping subsets in order to separate basis set optimization (set ‘SP175’) from independent subsequent quality assessment (set ‘TS8’) by several levels of cross validation (for a full list see Tables S1-S3). All crystallographic structures were obtained from the protein databank (PDB) ^19^.

The SP175 set (Table S1) is comprised of 71 globular protein structures and their corresponding CD spectra, assembled by Lees *et al.* ^20^ such that (1) its SS distribution reflects that of the full PDB, (2) it features high resolution PDB structures (X-ray diffraction and nuclear magnetic resonance structures, average resolution 1.9 Å) and (3) high quality synchrotron radiation CD spectra (wavelength range 175-269 nm), and (4) the set represents the major protein folds as defined by the CATH ^21^ database. The CATH database classifies proteins based on structural similarity as well as evolutionary relations. We mainly use set SP175 to derive and optimize SESCA basis sets and to determine their fitting accuracy, but this set also represents globular proteins during our analyses, e.g. during the PCA of protein CD spectra

The cross-validation set TS8 (Table S2) comprises eight additional globular proteins, selected from a set of 22 proteins, previously used for CD spectrum deconvolution ^22^. The CD spectra of the TS8 set were obtained from Hollósi *et. al*. ^10^ and cover a slightly shorter spectral range (178-260 nm) than those of the SP175 set. The TS8 crystal structures have an average resolution of 1.7 Å, and contain no missing residues. Set TS8 was used to assess the prediction accuracy of both hard and soft basis sets after optimization (Fig. S1), as well as the PDB2CD method.

The SP175 data set is divided into two subsets for the hard optimization approach, a larger training set of 64 proteins (TR64) for optimizing the basis spectra, and a smaller evaluation set EV9 of nine proteins (Table S3) for assessing the predictive power of basis sets during their optimization. Two additional proteins with a β-sheet architecture were added to set EV9 from Hollósi *et al.* to obtain a balanced distribution of main folds and sufficient sampling to avoid overfitting. Also, the protein structures of set EV9 (average resolution 1.6 Å) contain no missing residues.

A third subset of SP175 (GP59 for ‘globular protein’ set) comprising 59 globular proteins is used to estimate the average contribution of side chains to the CD spectra. Accordingly, GP59 was formed such that (1) its proteins maintain a wide variation of SS compositions and (2) their spectra are predicted with sufficient accuracy (see section 5.1). Twenty short peptides (the GXG20 set) with the consensus sequence of Ac-GXG-NH_2_ (X stands for any AA) were added to GP59 to form a reference set of ‘mixed polypeptides’ termed MP79. This final reference set is used to optimize all ‘mixed basis sets’ with both backbone and side-chain contributions.

### 3.2 Circular dichroism measurements

The CD spectra of set SP175 were provided by Kardos *et al.* and are deposited in the protein circular dichroism databank (PCDDB)^23^. The CD spectra of set TS8 were obtained from the literature (see Section 3.1). The CD spectra of set GXG20 peptides and the complex of the two disordered protein domains P53-AD2 and CBP-NCBD were recorded on the AU-CD beam line at the ASTRID2 synchrotron radiation source (Aarhus, Denmark) under similar conditions (298 K, in 50 mM NaF solution with Na_2_HPO_4_ buffer, pH = 7.1) over the wavelength range 178-300 nm. Protein concentrations (0.5-2.0 mg/ml) were determined from light absorption at 214 nm ^24^ and, when possible, at 280 nm (for both domains as well as GYG and GWG). Both peptides and protein domains were produced by the company Karebay using solid state peptide synthesis. The CD spectrum P53/CBP complex was recorded after 30 minutes of incubation time at room temperature with a 1:1 molar ratio. The CD spectra in all data sets were converted to Mean Residue Ellipticity ([Θ] or MRE) shown in the units of 10^3^ degree*cm^2^/dmol, abbreviated as kMRE.

### 3.3 Structural models and molecular dynamics simulations

The crystal structures of set SP175 and TS8 were obtained from the PDB ^19^ (entry codes and subset information provided in Tables S1-S3), as was the NMR solution structure of the P53/CBP complex (used in Section 5.4, PDB code 2L14). Structural ensembles for each GXG20 peptide were generated using a 10 μs long molecular dynamics (MD) simulation (recorded every 2 ns) using the GROMACS simulation package ^25^ (version 5.06) and the Charmm36m parameter set with the explicit TIP3P water model modified for the force field. ^26^

The simulations were performed under periodic boundary conditions at 298 K, with Na^+^ and Cl^−^ ions at 50 mM ionic strength and protonation states corresponding to pH = 7. The size of the simulation box was chosen so as to keep ~2 nm distance between any solute atom and the box boundaries, resulting in a simulation box with ~5500 atoms. All GXG20 simulations were started from an extended conformation.

The P53/CBP simulation was carried out similarly, except that the Charmm22* parameter set ^27^ was used and the simulation box contained ~82 000 atoms. The simulation was started using the first conformation of the NMR bundle, and protein conformations were recorded every 10 ns throughout a 10 μs long simulation trajectory, resulting in an ensemble of 1000 conformations. Here, the obtained structural ensemble was compared with the starting structure as well as with the 20 conformations of the full NMR bundle for predicting the measured CD spectrum and the C_α_ secondary chemical shifts.

### 3.4 Secondary structure determination

The SS of all proteins was determined from the protein structure using three different algorithms: DSSP (Dictionary of Secondary Structure for Proteins) ^28^, DISICL (DIhedral based Segment Identification and CLassification) ^29^, and the in-house algorithm HbSS (Hydrogen-bond based Secondary Structure) described below.

DSSP identifies eight SS elements (in Table S4) based on their distinctive backbone hydrogen-bonding patterns. DISICL uses two (ϕ,Ψ) backbone dihedral angle pairs to classify tetra-peptide segments into 19 SS elements (DS_det, Table S5), which are grouped into eight broader SS classes in the simplified DISICL protocol (DS_sim).

The HbSS algorithm was used to distinguish between parallel and antiparallel β-strands (Fig. S7), determined based on backbone hydrogen-bonding patterns. In addition, HbSS determined helical and turn-based SS elements (listed in Table S6) similarly to DSSP. The HbSS classification was also extended (HBSS_ext) based on the β-strand twist to determine the amount of left-handed, relaxed (non-twisted) and right-handed β-strands described by Ho *et al.* ^30^, with boundaries of 0° and 23°, respectively, for both parallel and anti-parallel strand arrangements. This extended structural classification protocol is directly comparable with the estimates of the deconvolution algorithm BestSel ^9^ (Table S7).

### 3.5 CD spectrum and SS predictions

Prior to analysis, crystallographic water, non-standard residues, and cofactors were removed from the crystal structures of the data sets. Residue numbers and chain codes were relabeled to ensure compatibility with the analysis software. For all entries of the reference protein sets, the AA composition and the SS content was determined (Section 3.4). The CD spectra of reference proteins were predicted using several SESCA optimized basis sets (see SM Tables S8-S9) and the SS compositions obtained, as well as using the DichroCalc and PDB2CD software and the processed reference structures. For PDB2CD, the SP175 reference set was used to predict the spectra, whilst for DichroCalc, the peptide parameter set from Hirst *et al.* was selected for CD predictions, including backbone charge-transfer transitions, as well as aromatic- and acidic side-chain chromophores. A PCA was applied to the SP175 CD spectra to determine the number of necessary spectral components and to probe correlations between the principal components, SS elements, and AA composition (see SM Section S5). For comparison, the SS content of each protein was estimated from their CD-spectrum using the deconvolution algorithms SELCON ^18^ and BestSel. These estimates were also included in the spectral component analysis (SM Section S5).

### 3.6 Averaging over prediction results

In this work, CD spectrum predictions were often used to determine properties averaged over a set of reference proteins such as the mean accuracy for the calculated spectra (*RMSD*_set_), or the error of structural models (*ΔSS*_set_). These obtained properties are influenced by the errors of the prediction method (for SESCA, the errors of the basis spectra). To reduce the effect of non-systematic errors from the prediction method, we calculated the mean property for each protein *j* from multiple predictions, denoted as

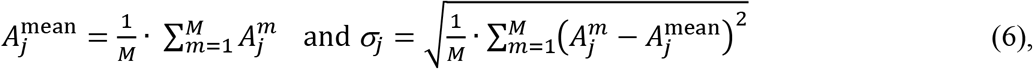

where 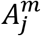 is the property *A* obtained for protein *j* using the prediction *m*, and *σ*_*j*_ is the scatter of the obtained properties. The average property over the protein set 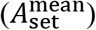 and its standard deviation (*σ*_set_) was then calculated as

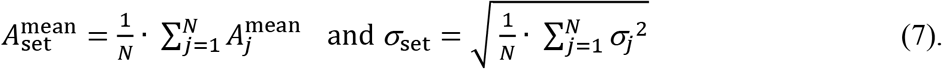

When calculating mean properties for SESCA, we averaged over results obtained using four optimized basis sets: DS-dT, DSSP-1, HBSS-3, and DS5-4, which differ in size and in the underlying SS classification protocol (see Table S8). In cases where the properties were not SESCA specific – such as the scaling factors 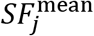 used in Section 5.2 – two other methods, namely PDB2CD and BestSel_der, were also included in the calculations. PDB2CD predicts CD spectra from the reference crystal structures, but (unlike SESCA) does not rely basis spectra to do so. BestSel_der calculates the CD spectra only from the SS composition predicted by the established deconvolution algorithm BestSel ^9^ and therefore does not contain systematic errors from the reference structures. Note that the BestSel_der basis set was derived using the methodology described in SM Section S1.

## Results and Discussion

We present our results in two sections. In Section 4, we assess the accuracy of our semi-empirical spectrum calculation approach, SESCA, using SS information only. In Section 5, we include the effects of side groups and backbone dynamics, and explore the potential to further improve SESCA predictions for folded proteins, peptides, and IDPs.

## 4. Secondary structure based CD calculations

As the main determinant of prediction accuracy for SESCA is the choice of the basis set (defined in the Section 2.1), we first assess the effects of this choice in Section 4.1. To this end, using the 71 proteins of the SP175 reference set, we derived different basis sets with varying number and shape of the basis spectra, using different underlying SS classification protocols. All of these basis sets were determined by the hard optimization approach (see Section S2) and were subsequently cross-validated against the proteins of the TS8 reference set. The prediction and fitting accuracies for all basis sets were determined as described in Section 2.4, and the most predictive basis sets were analyzed further. In Section 4.2, we compare the prediction accuracies achieved with the two established CD prediction methods DichroCalc and PDB2CD. Possible limitations of SESCA are explored in Section 4.3 using PCA and soft basis sets (see Section 2.4). Finally, in Section 4.4 we determine the sensitivity of spectrum prediction accuracy with respect to errors in the SS composition of the protein model.

### 4.1 SESCA basis set assessment

We derived basis sets with three to a maximum of 19 basis spectra, using five SS classification protocols in total (See Section 3.4). Each basis set was optimized using the TR64 and EV9 subsets of SP175 for training and evaluation, respectively. Subsequently, the basis spectra were recalculated from the full SP175 set, and thereafter, the basis sets were cross-validated against the TS8 set. The 10 top-ranking optimized basis sets achieved an average prediction accuracy of RMSD_TS8_ = 3.2 ± 0.6 kMRE units (10^3^ deg*cm^2^/dmol). Notably, the average fitting accuracy RMSD_SP175_ = 3.6 ± 0.2 kMRE is very similar, indicating little to no overfitting

Table S8 (upper half) lists the average accuracies of the top-ranking basis sets for the predicted CD spectra of the SP175 set, its training (TR64) and evaluation (EV9) subsets, as well as the TS8 cross-validation set. For comparison, an additional seven non-optimized basis sets are shown as well (Table S8, lower half). To assess the effects of the underlying SS classification, we compared the most accurate predictions for each classification algorithm from the basis sets DS6-1 (DISICL), DSSP-T (DSSP), and HBSS-3 (HbSS), respectively (marked by asterisks in Table S8). These three basis sets achieved an RMSD_TS8_ of 3.04 ± 0.57, 3.01 ± 0.54, and 3.29 ± 0.63 kMRE, respectively. As these prediction accuracies are similar within statistical error, the choice of the SS classification protocol does not seem to markedly affect the predictive power of SESCA.

Unexpectedly, for the three most accurate basis sets mentioned above – regardless of the underlying classification algorithm – the prediction accuracy is also somewhat higher than the fitting accuracy (RMSD_SP175_) of the SP175 reference set (3.27 ± 0.23, 3.82 ± 0.25, and 3.75 ± 0.23 kMRE, respectively). We tentatively attribute this to the slightly higher average data quality of the TS8 set.

Notably, all ten top-ranking basis sets listed in Table S8 contain between three and six basis spectra. In fact, in all basis sets considered, the number of basis spectra drops to eight or less but never below three during optimization. This finding indicates that at least three distinct basis spectra are required to explain the diversity of the observed CD spectra, and that the limited accuracy of both structures and measured CD spectra in the reference data sets does not allow for more than eight basis spectra without overfitting. This finding is also in line with the prediction accuracies of the non-optimized basis sets shown in Table S8 (bottom) as well as with the PCA analysis of the best achievable fitting accuracy in Section 4.3.

Figures 3A-3C show the basis spectra of the three most accurate basis sets. As the most distinctive common feature, all three sets contain a very similar basis spectrum largely representing α-helical structure elements (blue lines), which, as one should expect, closely resembles the basis spectra attributed to α-helices in previous studies. ^4,31,32^ The β-strand basis spectra are also similar to each other, although smaller differences exist depending on the underlying β-strand classifications. These two basis spectra appear consistently throughout the top-ranking optimized basis sets as well (examples shown in Figs. S8-S15).

**Figure 3:**
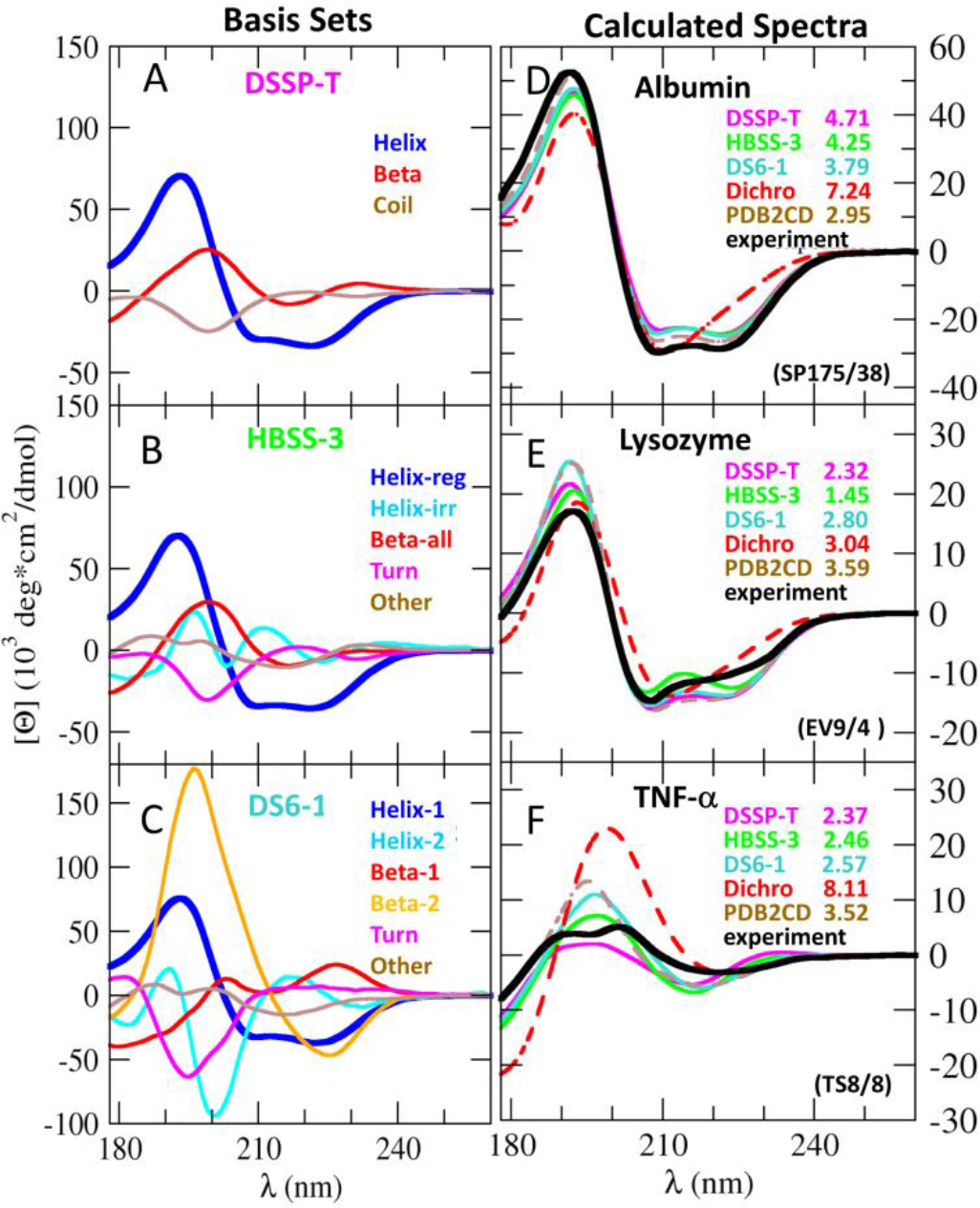
Comparison of basis spectrum sets and predicted CD spectra. Left column: basis spectra of three high-accuracy basis sets (DSSP-T, HBSS-3, and DS6-1). Basis spectra for similar secondary structure classes are indicated by similar colors. Right column: measured CD spectra (black solid lines) for three example proteins (human serum albumin, lysozyme, and tumor necrosis factor α) compared to predicted CD spectra from three SESCA basis sets shown in the left column, DichroCalc (Dichro), and PDB2CD. The respective RMSD of the measured spectrum is given for each method in the plots in 10^3^ deg·cm^2^/dmol (kMRE) units.

The third basis spectrum in Fig. 3A (brown line) shows the typical negative peak at 198 nm usually attributed to random coil structures. Notably, the basis sets with more detailed SS classifications in Figs. 3B-3C allow us to more precisely attribute this feature to certain turn types (Turn-cap, β-bulge, Turn type I) and extended structures (poly-proline-helical and β-cap SS elements) as represented by the Turn and Helix2 classes (Fig. 3C, cyan and magenta). The fact that no such peak is seen in the basis spectrum representing the remaining unassigned structures (basis sets: Other) in Figs. 3B and 3C shows that the negative peak previously assigned to ‘random coil’ can indeed be mainly attributed to these three secondary structure elements. The fact that different SS sub-classifications yield similar average fitting and prediction accuracies can be exploited to enhance the sensitivity for particular SS changes, e.g., from left-handed helices to parallel β-strands.

In summary, using different SS classifications we derived several basis sets which best predict the experimental protein spectra. All of these are comprised of three to six basis spectra, which seem to provide the best trade-off between available information content and overfitting due to experimental inaccuracies. Consistently, the measured CD spectra are predicted with an accuracy of ~3.1 kMRE (RMSD_cross_) while avoiding overfitting.

### 4.2 Performance comparison

To evaluate the performance of our method, we used the SP175 and the TS8 data sets to compare SESCA with the other two available CD calculation methods outlined in the introduction, DichroCalc and PDB2CD. We emphasize that these algorithms represent different approaches of quantitative predictions based on CD spectroscopy, and DichroCalc – being an *ab initio* spectrum calculation method – was not parametrized to reproduce any particular protein reference set. In contrast, PDB2CD was developed based on the SP175 reference protein set, which enables a fair comparison to SESCA using the two data sets above.

Table S9 lists the average accuracies (RMSD) of the CD spectra calculated from the crystallographic structures of set SP175 and TS8 for both DichroCalc and PDB2CD. As can be seen, the average RMSD of CD spectra predicted by DichroCalc are 6.06 ± 0.33 and 6.12 ± 1.12 kMRE units, respectively. As expected for an *ab initio* approach, DichroCalc yields similar average accuracies for both data sets. However, these accuracies are markedly lower than those achieved by SESCA or PDB2CD. To investigate the reason for this finding, Figs. 3D-3F show CD spectra for three representative sample proteins: one α-helical (D), one α/β (E), and one β-sheet (F) protein. The comparison of the calculated CD spectra (red dashed lines) with the respective measured spectra (black lines) shows that DichroCalc only determines the most prominent spectral features, without reproducing the sub-structure of the peaks.

Because the purely empirical method PDB2CD calculates a weighted average of CD spectra measured for structurally similar reference proteins of the SP175 set, it is – unsurprisingly – markedly more accurate for this set (RMSD_fit_= 2.40 ± 0.26 kMRE, brown dashed lines in Figs. 3D-3F) than any of the SESCA basis sets, or DichroCalc. However, in contrast to DichroCalc and the optimized SESCA basis sets, its prediction accuracy for the TS8 set drops markedly (RMSD_cross_= 4.73 ± 0.69 kMRE), suggesting less predictive power compared to our SESCA basis sets (RMSD_cross_ ranging from 3.0 to 3.9 kMRE).

Note that this result differs from a previous PDB2CD cross-validation study by Mavridis *et al.* ^17^ which used a set of 14 protein structures (‘TS14’, details in Table S10) and reported prediction accuracies that are very similar to their corresponding SP175 fitting accuracy. To investigate this, we performed a cross-validation on the TS14 set too, and obtained an RMSD_set_ of ~3.8 kMRE units both for SESCA basis sets and PDB2CD, whilst DichroCalc performed somewhat worse (~5.6 kMRE). We attribute this discrepancy between the cross-validation results to the fact that Mavridis *et al.* ^17^ report relative deviations from the measured CD spectra, whereas we report absolute deviations. Notably, from the best eight cases of the TS14 set for which the previous study ^17^ reported very accurate predictions from PDB2CD, four β-crystallin proteins were also part of the SP175 reference set.

Because SESCA uses relatively few distances and angles for determining the SS composition, as well as pre-calculated basis sets, it should be computationally much more efficient than PDB2CD or DichroCalc. To test this expectation, we benchmarked all three methods using the structure of an average-sized protein (14-3-3ζ, 490 AAs, PDB code 2WH0) The spectrum of this protein was calculated by SESCA in 0.3 seconds for a single structure, and the average CD spectrum for an ensemble of 1000 conformations was determined in just under five minutes. The PDB2CD and DichroCalc benchmarks on the same protein, using the publicly available servers took nineteen and eight minutes, respectively (excluding queuing times), which as expected was longer than SESCA. High efficiency CD spectrum predictions are particularly important for, e.g., the iterative refinement of structural ensembles, where the calculation of CD spectra for 10^4^ -10^5^ structures are often required.

### 4.3 Basis set accuracy limits

Next, we investigated to what extent the accuracy of our basis sets could possibly be improved (a) when including all available structural information and (b) when using secondary structure information only. As the best available representation of globular proteins^20^, we used the SP175 reference set for this purpose. The best achievable accuracy was determined by PCA of the reference spectra as defined in Section 2.4 (see SM Section S5 for details). By definition, the fitting accuracy RMSD_fit_ achieved using the first *n* principal components provides the upper accuracy limit for a basis set of size *n* (see the chapter on dimensionality reduction in Ref ^33^).

Figure 4A shows this absolute limit (black dotted-dashed line) as a function of basis set size. For example, with two basis spectra – even when using the full structural information – an accuracy not better than 1.6 kMRE can be achieved. This optimal accuracy rapidly decreases from an initial 6.40 kMRE (using only the average spectrum and no structural information) to 1.34 kMRE using up to three basis spectra, followed by a more gradual decrease to 0.24 kMRE for eight basis spectra, and a slow decline to 0.18 kMRE at 10 basis spectra. Therefore, for accurate predictions at least three basis spectra are required, whereas using more than eight basis spectra does not increase the achievable accuracy further, in line with the size range of optimized basis sets obtained in Section 4.1.

**Figure 4:**
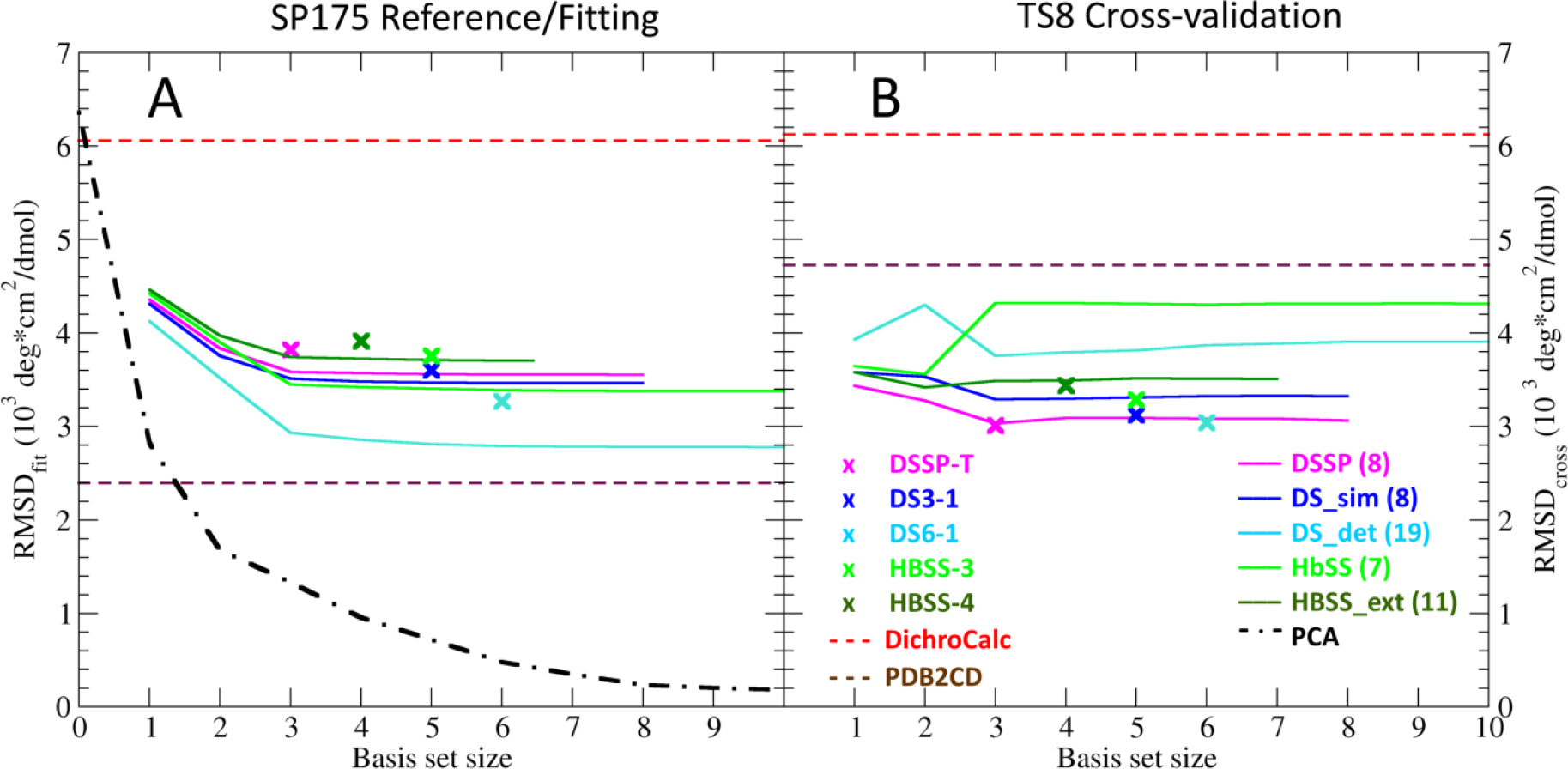
Fitting (RMSD_fit_) and prediction (RMSD_cross_) accuracies for globular proteins. Average accuracies are shown for A) globular proteins of the SP175 reference set, used for basis set determination and B) for the globular proteins of a small independent set TS8, used for cross-validation. The average RMSD values between measured and calculated CD spectra are shown as a function of basis set size, colored according to the SS classification protocol used (see legend on the right, the number of SS elements is shown in parentheses). Crosses mark the most predictive hard SESCA basis sets. For comparison, the maximum achievable accuracy (black dash-dotted line, determined via PCA in Section S11) is shown in panel A, together with the accuracy limit of determining the CD spectra only from the SS composition of the proteins (outlined by soft SESCA basis sets derived in Section S4, shown as colored solid lines). In panel B, the accuracy of most soft basis sets (but not of hard basis sets) is reduced significantly due to overfitting. The average accuracy of DichroCalc and PDB2CD (red and brown dashed lines) are also shown on both plots to compare their predictive power with SESCA basis sets.

Because the actual achievable accuracy of our basis sets is reduced by the limited structural information contained in the SS composition, as well as by the error in determining this composition from the crystal structure of the protein, we also determined the best achievable accuracy when predicting the spectra from SS information. To account for these limitations, we determined basis sets using the soft optimization method (described in Section 2.4), which varies the SS assignment and the shape of basis spectra iteratively to acquire the best possible fitting accuracy (details in Section S4).

The colored lines in Fig. 4A show the fitting accuracy (RMSD_fit_) of the acquired soft basis sets for every possible basis set size using the SS information determined by the five structure classification protocols described in Section 3.4 (DSSP, DS_det, DS_sim, HbSS and HbSS_ext). Although these lines also decrease monotonically with the basis set size, the best accuracy of only 3.2 ± 0.5 kMRE is achieved, depending on the SS classification, with the lowest RMSD_fit_ of 2.8 kMRE obtained for the DS_det protocol with 19 SS elements (light blue line). The ~2.5 kMRE difference between PCA and DS_det basis sets shows that the limited information of the SS composition (derived from a crystal structure) markedly reduces the accuracy of the predicted CD spectra.

The fitting accuracies achieved by the most predictive optimized basis sets (marked by crosses in Fig. 4A, data in Table S8) are all within 0.5 kMRE of the determined accuracy limit for their respective size and SS classification. This also indicates that further significant improvements in basis set accuracies are unlikely based only on the crystal-structure derived SS information.

Figure 4B compares the predictive power of hard (crosses) and soft (lines) basis sets, as determined by cross-validation. As can be seen, the average prediction accuracy of hard basis sets (RMSD_cross_ = 3.2 ± 0.6 kMRE) is also within the accuracy limits established above (3.2 ± 0.5 kMRE), which indicates that overfitting is largely avoided. In comparison, overfitting seems to be more severe for the soft basis sets; unlike their fitting accuracies in Fig. 4A, prediction accuracies for soft basis sets are systematically lower than for hard basis sets, and in addition, do not improve monotonically with basis set size. The overfitting seems to be particularly severe for basis sets using SS classification protocols with more than eight structural elements (DS_det and HbSS_ext, light blue and green lines, respectively). Because of their lower predictive power, soft basis sets are not considered further for predicting protein spectra in this study.

It is instructive to also compare the average RMSDs obtained from DichroCalc and PDB2CD, respectively, to the accuracy limits determined above. The calculated average RMSDs of DichroCalc (Fig. 4, red dashed lines) are close to what can be achieved using only the averaged CD spectrum of the SP175 set (6.4 kMRE, PCA-0). The observed variation of the individual RMSDs is large, however, and some of the DichroCalc predictions are rather accurate (best RMSD 1.9 kMRE). On the other hand, the RMSD_fit_ obtained by PDB2CD (2.4 kMRE) is even below that of the soft SESCA basis sets, probably because PDB2CD also uses tertiary structure information. This high accuracy, however, is not reached during the cross-validation, where hard SESCA basis sets yield higher predictive power.

### 4.4 Model validation based on CD predictions

The CD spectrum predictions characterized above enable us to address our main question: can one assess the quality of a protein structural model from the deviation (RMSD) between its predicted and measured CD spectrum? Specifically, we focus on two questions: 1) Given a RMSD value, how large is the total error (ΔSS) in the model’s secondary structure composition, and 2) given two proposed models, what is the minimum difference in the SS composition (ΔSS_min_) to reliably discriminate between the two?

To answer these questions, one needs to determine how *RMSD*_*j*_ between the measured and the predicted spectra of protein *j* depends on *ΔSS*_*j*_, which is described in Section 2.5. In summary, to calculate *ΔSS*_*j*_, two parameters are required: *m*_*j*_ which defines the proportionality between *ΔSS*_*j*_ and SS-dependent error of the predicted CD spectrum (*M*_*j*_ = *m*_*j*_ · *ΔSS*_*j*_) and 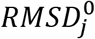 which is the average SS-independent error of the predicted spectrum, that provides an off-set for *RMSD*_*j*_ if the SS-dependent error is small, or the error terms are correlated.

Equation 5 in Section 2.5 allows one to calculate *ΔSS*_*j*_ for any structural model of protein *j* from *RMSD*_*j*_, provided that one knows *m*_*j*_, 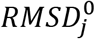, and the error terms for protein *j* are statistically independent. Unfortunately, both parameters depend on the correct SS composition 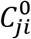, which is typically unknown. Furthermore, it is not known if the two error terms for a given protein are actually statistically independent. To address these problems, in Section S8 we developed an error model for each basis set that enables one to estimate *ΔSS*_*j*_ without prior knowledge of 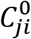, and allowing for possibly correlated error terms. The model is based on fitting the average parameters *m*_*f*_ and *R*_*f*_ to the *ΔSS*_*j*_ vs. *RMSD*_*j*_ values of SP175 reference proteins, which were used to derive the basis sets. Note that as shown in Fig. S16, *ΔSS*_*j*_ for each reference protein of the set was obtained by predicting its SS composition from the reference spectrum (through deconvolution, see Section S7) and comparing it to the SS composition of its reference structure. Similarly, *RMSD*_*j*_ values were obtained by predicting CD spectra from the reference structure and by comparing them to the reference spectra.

In Figure 5A the accuracy of our error model is assessed for the DS-dT basis set. For this basis set, we obtained *m*_*f*_ = 31.3 ± 1.3 kMRE and *m*_*f*_ = 1.79 ± 0.52 kMRE, which agree well with the mean of the individual *m*_*j*_, and 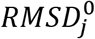 parameters calculated from the obtained 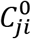, whose distributions are shown in Figs. 5B-5C, respectively. Using these fit parameters, the best estimate 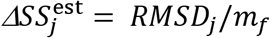 for a given *RMSD*_*j*_ (red line in Fig. 5A) reproduced the *ΔSS*_*j*_ values obtained with an average deviation (*ΔΔSS*_SP175_) of 2.6%. If we allow for strong correlations between SS-dependent and SS-independent error terms, the parameter *R*_*f*_ and its uncertainty *σ*_*R*_ determine the uncertainty of the best estimate (*σ*_*S*_ ≈ (*R*_*f*_ + *σ*_*R*_)/*m*_*f*_). However, because the correlation between error terms is very weak for most reference proteins, a more narrow estimate for the uncertainty was also defined as *σ*_*S*_ ≈ *ΔΔSS*_SP175_. The two estimated uncertainties for the DS-dT basis set are shown as green and purple dashed lines, which contained 67 (94%) and 53 (75%) of the 71 obtained reference data points within their boundaries, thereby confirming that our error estimates are indeed appropriate.

**Figure 5:**
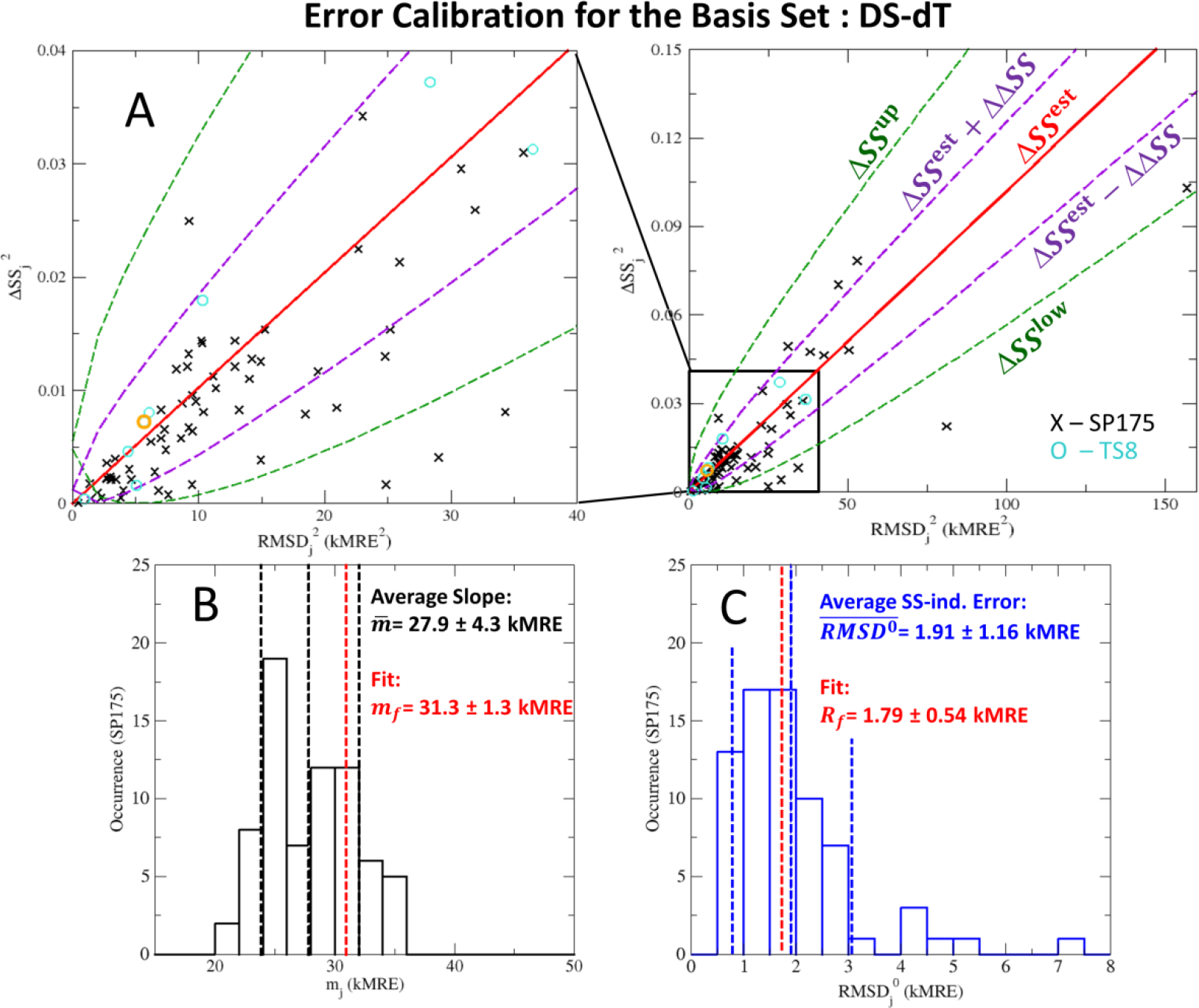
Error calibration for the basis set DS-dT. A) The error in secondary structure composition (ΔSS_*j*_) for a proposed structural model of protein *j* is estimated based on the RMSD_*j*_ between its predicted and the measured CD spectrum. The best estimate of ΔSS_*j*_ is shown as a red solid line. The estimated upper and lower bounds of ΔSS_*j*_ are shown as dark green dashed lines. The prediction RMSD vs. the real value of ΔSS_*j*_ for the 71 proteins in the SP175 dataset are shown as black crosses. The reference proteins of the TS8 cross-validation data set are shown as light blue circles, with a representative example, Prealbumin indicated in orange. B) and C) show the distribution of the SS-independent error (RMSD_*j*_^0^) and the slope of the SS-dependent error (*m*_*j*_) for all proteins in the SP175 reference set, determined by deconvolution of their measured CD spectrum. The fit parameters for our error model representing the average of these properties are shown as vertical, red dashed lines. The average error of the estimated ΔSS_*j*_ (±ΔΔSS, in purple) is also indicated.

This error model now allows one to answer the first of the above two questions, to assess the quality of any given structural model by estimating *ΔSS*_*j*_ from the RMSD between its predicted and measured CD spectrum. As an example, we estimated *ΔSS*_*j*_ for the crystal structure of Prealbumin (TS8/3), a reference protein that was not used in our error model determination. The measured CD spectrum of Prealbumin deviates by an RMSD of 2.38 kMRE from the spectrum predicted from its crystal structure using the DS-dT basis set. The error model for this basis set estimates *ΔSS*_*j*_ between 0.2-15.5%, with a 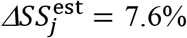, which is very close to the *ΔSS*_*j*_= 8.5% obtained from CD deconvolution (Fig. 5A, orange circle).

Performing the same test on the *ΔSS*_*j*_ values obtained for the TS8 cross-validation set (light blue circles in Fig. 5A) showed that for the DS-dT basis set all are within the tighter estimated boundaries with an average deviation of *ΔΔSS*_TS8_ = 1.7% for the best estimates. Finally, we performed the same error calibration for four optimized basis sets listed in Section 3.6 and obtained similar results with 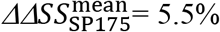, 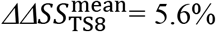, with an average 95% and 75% of data points contained between the two estimated boundaries, respectively, further corroborating our approach.

To address the second question concerning the sensitivity of our model validation, we consider two proposed models with 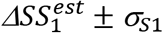 and 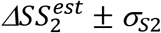, with typical experimental errors in the reference structures and measured CD spectrum. To reliably discriminate between the two structural models, the difference 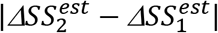 has to be larger than 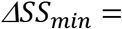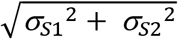. If we assume the most likely scenario, with weak correlations between the SS-dependent and SS-independent error terms for both structural models, the uncertainties σ_*S*1_ = σ_*S*2_ = *ΔΔ*_SP175_ and thus 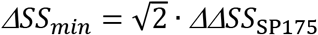. These assumptions yield *ΔSS*_min_ values between 3.7% (DS-dT) and 11.2% (DS5-4) for the optimized basis sets tested. We note that both the average *ΔSS*_*j*_ and *ΔSS*_min_ increases with increasing basis set size. The trend in the *ΔSS*_*j*_ values obtained can be explained by the larger basis sets extracting more structural information from the models, and thus registering small deviations between the reference and solution structures that smaller basis sets do not detect. The increasing *ΔSS*_min_ values are likely due to the larger variation of *m*_*j*_, and 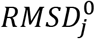 parameters observed for larger basis sets.

We also note that the obtained distributions of both 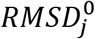 (shown in Fig. 5C) and *ΔSS*_*j*_ values are asymmetric in the SP175 set. For over 60% of the proteins, both error terms are below the average, whereas relatively few proteins show exceptionally large errors. In particular, all four proteins outside the estimated error range depicted in Fig. 5A also exhibit large SS-independent errors 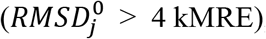, suggesting that the measured CD spectra of these proteins cannot be accurately predicted by the DS-dT basis set. In addition, seven proteins in the SP175 reference set have a large SS-dependent error (*ΔSS*_*j*_ > 20%), which may have reduced the accuracy of our basis sets. To further improve the accuracy of SESCA predictions, in the next sections, we investigate several potential sources of these errors.

## 5. Improving the CD prediction accuracy

In Section 4, we derived several SESCA basis sets to predict the CD spectra of globular proteins and determined their achieved prediction accuracy. In this section, we focus on whether the prediction accuracy of our basis sets can be further improved by changing the applied methodology or the reference set. First, we study how much the use of crystal structures as structural models affects the spectrum prediction accuracy. Second, we analyze reference proteins with large CD prediction errors in our training set, and their effects on the robustness of SESCA predictions. Third, we determine and include contributions from the AA side chain groups. Finally, we present resulting improvements of the SESCA methodology through the example of a highly flexible protein complex.

### 5.1 Effect of the structural models: solution vs. crystal structure

To calculate the average deviation between the SS composition of crystal and solution structures for globular proteins, we averaged over the total model error (*ΔSS*_*j*_) of reference proteins determined in Section 4.4. Because the individual *ΔSS*_*j*_ values for the same protein scatter significantly (6.3% on average) depending on which basis set was used to deconvolve the measured CD spectrum, we calculated mean deviations 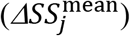 using four optimized basis sets (See Section 3.6). The 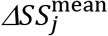 values show similar average differences between solution and crystal structures for the SP175 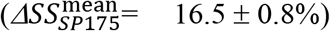 and the TS8 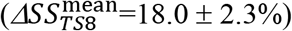 reference sets.

To assess how much these structural differences limit the accuracy of SESCA predictions, we also determined the difference between the mean prediction error 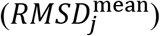 and the mean SS-independent prediction error 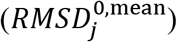 for each protein *j*, averaged over the four basis sets above. Based on these calculations, the structural differences between crystal and solution structures contribute a sizeable (up to 2.0 kMRE) portion of the mean fitting accuracy 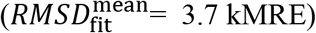 for the SP175 reference set, and introduce a similar error to the prediction accuracy 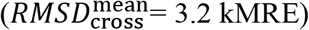 for the TS8 cross-validation set.

From the above results, we conclude that using the crystal structure is a major limitation to the accuracy of CD spectrum predictions for our reference proteins. We speculate that structural models derived from nuclear magnetic resonance (NMR) spectroscopy or MD simulations may allow more accurate CD spectrum predictions, as they reflect the average solution structure of the protein better. Furthermore, for several proteins of the SP175 reference set, the CD spectra were predicted with relatively poor accuracy even from the ideal SS composition, regardless of the basis set used. This points to either systematic errors in the measured CD spectra of these proteins, or to strong contributions to the spectrum that cannot be predicted from the SS composition. We investigate these possibilities in the following sections.

### 5.2 Potential measurement errors of the reference set

Next we asked if a particularly large mean error 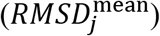 of the calculated spectra points to systematic measurement errors in our reference sets. To obtain a more precise 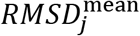 for each protein in the SP175 and TS8 reference sets, we averaged over the errors (*RMSD*_*j*_) of the CD spectra calculated by the four SESCA basis sets and two other methods, PDB2CD and BestSel_der (see Section 3.6). Then, we selected twelve proteins from the SP175 reference set (highlighted in Fig. S3A and listed in S3B) and one from TS8 as particularly hard to predict, because their 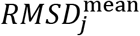 exceeded the average prediction error 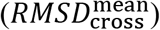 of the TS8 set (solid line) by more than one standard deviation (top dashed line) as described in Section S9. Recalculating the basis spectra of SESCA basis sets from the SP175 set without the 12 outliers (GP59 set) indeed improved the mean RMSD from 3.3 to 2.7 kMRE units (Fig. 6A, black and blue lines), whereas the mean prediction accuracy of the basis sets shown in Fig. 6B was essentially unchanged. Therefore, we conclude that the prediction accuracy of our basis sets is robust with respect to the presence of the hard-to-predict proteins.

**Figure 6:**
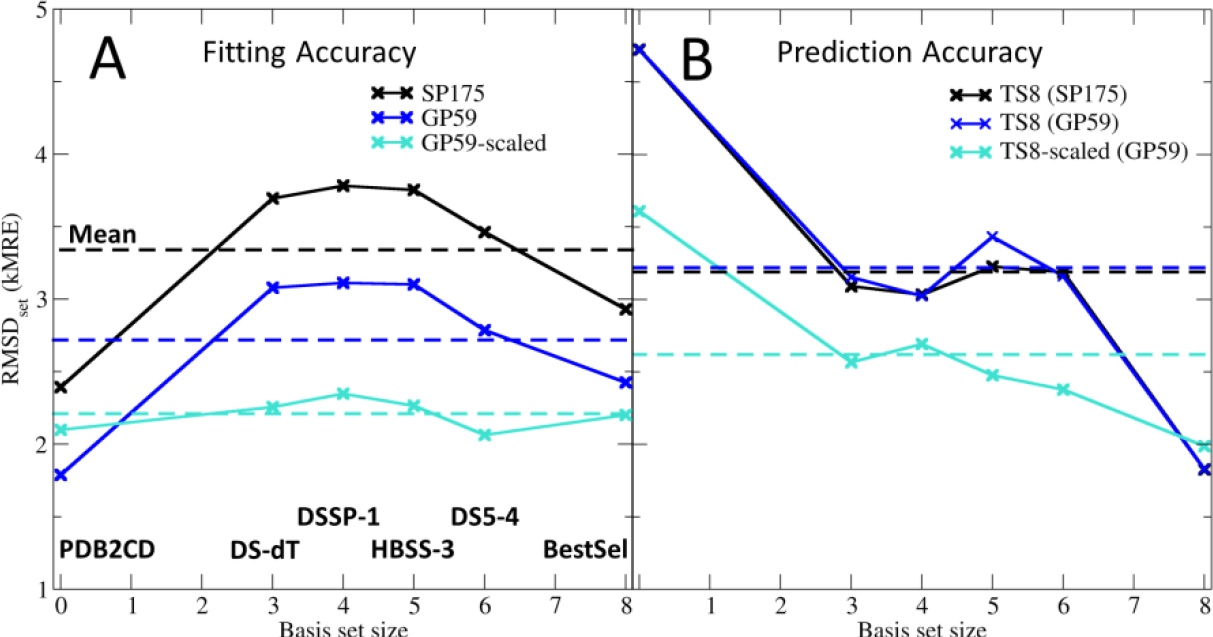
Changes in the mean fitting (A) and prediction accuracy (B) upon the removal of outliers and intensity scaling. Method independent mean RMSDs (shown as dashed lines) for the fitting (SP175) and the cross-validation (TS8) data sets were calculated as the average RMSD_set_ of six spectrum prediction methods (crosses) including PDB2CD, four optimized SESCA basis sets of different sizes and underlying classification schemes (DS-dT, DSSP-1, HBSS-3, and DS5-4), and the BestSel_der reconstruction basis set. Accuracies calculated for the original unmodified data sets are shown in black, those computed after the removal of hard-to-predict proteins from the SP175 set (GP59) and subsequent recalculation of the SESCA basis spectra are shown in dark blue, and those determined after additional rescaling of measured CD intensities are shown in cyan.

Interestingly, for five of the 13 outliers the calculated CD spectra agree well with the measured CD spectra after a simple rescaling (blue in Fig. S3B, example in Fig. S3C). This finding suggests that at least in these cases, inaccurate intensity normalization is the major source of RMSD between calculated and measured spectra, most likely due to uncertainties in the independently measured protein concentrations. For two outliers (magenta in Fig. S3B), intensity scaling improved the agreement with the predicted spectra, but peak positions or relative peak intensities still differed, most likely due to a large deviation between solution structure of the protein during CD measurements and the reference structure used. For the remaining six proteins (marked red, including Jacalin shown in Fig. S3D) scaling factors reduced the RMSD between the measured and predicted spectra, but did not yield a good agreement between the two even after fitting the SS composition of the protein model, which suggests additional contributions to measured CD spectra (discussed in SM Section S9).

To test whether inaccuracies in the measured concentrations may generally limit the accuracy of our CD spectrum calculations, we applied scaling factors 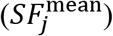 to the measured CD spectra of all proteins in the SP175 and the TS8 data sets. These scaling factors were determined based on the six predicted spectra per protein from which 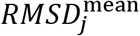 values were calculated (see Section 3.6). First, we computed the six individual scaling factors (*SF*_*j*_) that minimize the RMSD between the measured CD spectrum and one of the calculated spectra, then we averaged them to obtain a mean scaling factor 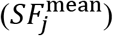.

Indeed, as shown in Fig. 6 (cyan lines), rescaling the measured spectrum intensities by 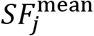 improved both 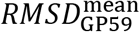 from 2.7 to 2.2 kMRE and 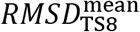 from 3.4 to 2.7 kMRE units. The scaling factors that achieved this improvement averaged around 0.89 and 1.02 for the SP175 and TS8 reference sets, respectively, with scattering by standard deviations of 0.29 and 0.25 around these averages. The latter variations are in line with typical uncertainties of 30-15% ^34–36^ for the two mainly employed concentration measurement methods, quantitative amino acid analysis and UV absorption. In addition, for seven proteins in the SP175 set independent CD measurements are reported by Hollósi et. al ^10^. For these proteins, the intensity of the measured CD spectra differs by factors between 1.1 and 0.8, further supporting the use of scaling factors to match our predictions.

### 5.3 Including side-chain contributions to the CD spectrum

Because the best achievable prediction accuracy (Section 4.3) is still markedly higher than the ones we obtained based solely on the SS composition (even after rescaling the measured CD spectra), including additional information should further improve the CD prediction accuracy. As the second most common type of chromophores in proteins, we therefore quantified the contribution of AA side chain groups to protein CD spectra in the far-UV range, and included these contributions within the SESCA scheme. We note that side-chain contributions are also considered as optional corrections in DichroCalc, and some deconvolution basis sets also include side-chain basis spectra ^5^.

To this end, we assembled a new reference set (MP79) comprised of 59 globular proteins (GP59) and 20 short peptides (GXG20) containing only a single side chain (see Section 3.1). Then, we calculated and subtracted the backbone contributions (see Section S3) and derived the average CD signal of all 20 AA side chains (shown and discussed in Section S9). Overall, we found that almost all side chains contribute markedly to the CD signal in the far-UV range (Fig. S4B), and to a similar extent as the protein backbone. Notably, the obtained side-chain basis spectra also differ significantly from CD spectra of individual AAs (Fig. S4A).

The analysis of the PCA basis sets (Section S11) suggests that only a few basis spectra are actually required to represent the contribution of the side chains. Therefore, to obtain basis sets that can optimally represent side-chain contributions, we combined the 20 AA side chains into classes (e.g. in Fig. 7C), thus allowing each basis spectrum to represent multiple side chains. The assignment of the AA side chains to these classes was optimized by the hard optimization scheme, similarly to the assignment of SS elements for backbone contributions. Incorporating the side chains into our basis set determination protocol (described in Section S3) yielded ‘mixed’ basis sets where some of the basis spectra represented the SS-dependent backbone contribution of the peptide bonds, whilst others provided the side-chain contributions, that depend only on the AA composition.

**Figure 7:**
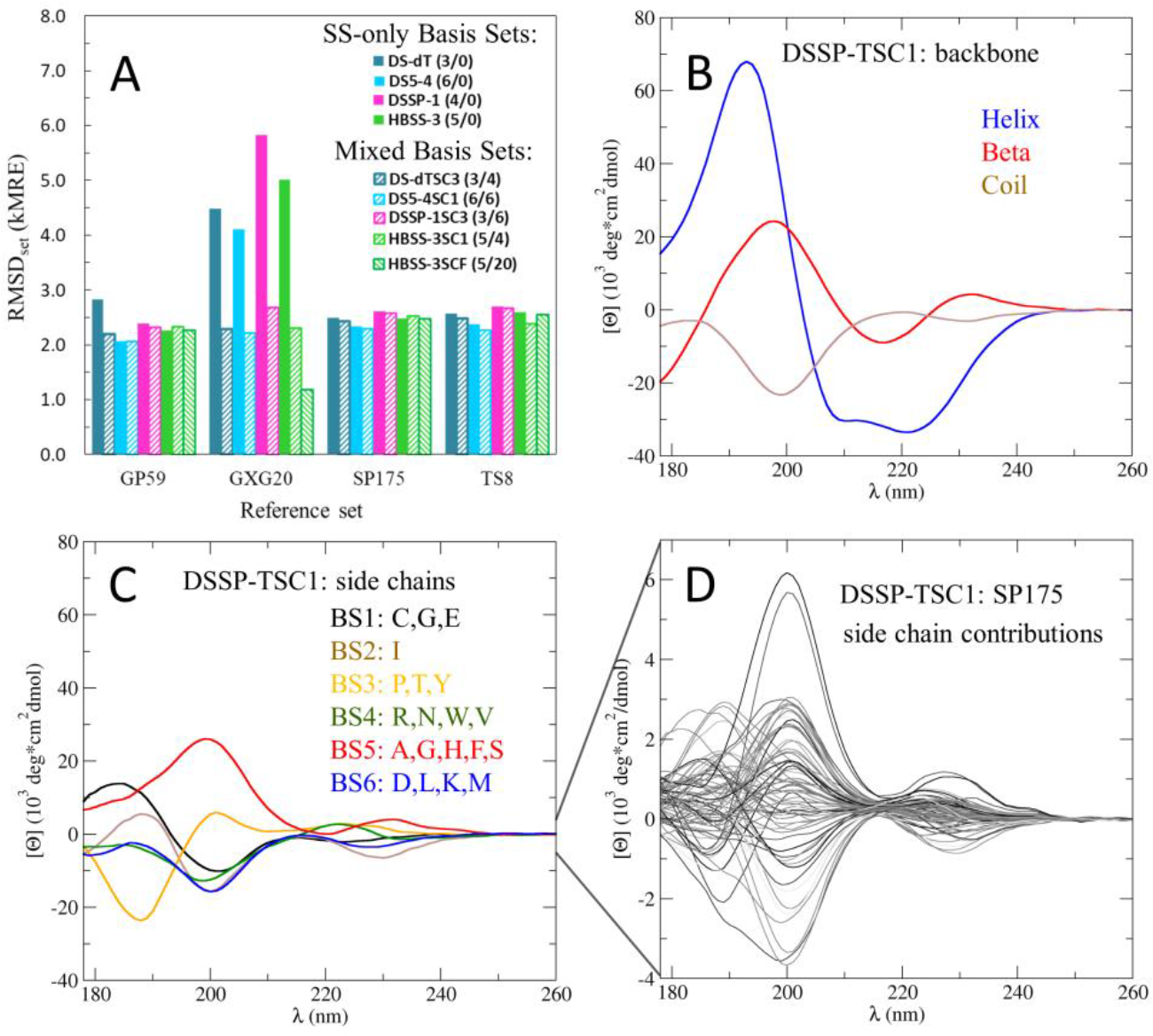
Comparison of backbone and side-chain contributions to the calculated CD spectra. A) Comparison of achieved average accuracies (RMSD_set_) for the globular protein (GP59) and short peptide (GXG20) subsets of the used training set, as well as for the SP175 reference set, and the TS8 cross-validation set. Different colors indicate different basis sets with (shaded) and without (filled) side-chain corrections; legends show the name of the basis set followed by the number of used backbone and side-chain basis spectra in parentheses. B) Backbone and C) side-chain basis spectra for the DSSP-TSC1 basis set; the grouping of side chains is indicated by one-letter codes of their respective AAs. D) Combined side-chain contributions for all SP175 proteins from the DSSP-dT1SC basis set; note the magnified y-axis.

The resulting optimized mixed basis sets (examples shown in Figs. S17-S22) typically include 3-6 backbone basis spectra and 4-7 side-chain basis spectra, with one or two basis spectra representing the positive CD signals of the aromatic residues. Figure 7A compares the average RMSD_set_-s achieved by optimized basis sets with (shaded bars) and without (solid bars) side-chain contributions. For a fair comparison, predictions for all globular proteins were compared to their rescaled reference spectra, and the basis spectra of SS-only basis sets were recalculated using the rescaled GP59 set (as discussed in Section 5.2). As the figure shows, including side-chain information improves both 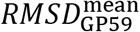 (training) by 0.16 kMRE and 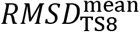 (cross-validation) by 0.11 kMRE. Notably, the gain in prediction accuracy is larger (0.29 kMRE) when the mixed basis sets are compared to the original optimized basis sets while predicting the unmodified TS8 spectra. The relatively small influence of the side groups for globular proteins is in line with the PCA analysis of the SP175 spectra, which suggested an upper limit of ~0.4 kMRE, and underscores the robustness of the secondary-structure based SESCA predictions.

Given that the intensities of the backbone and the side-chain basis spectra are rather similar (Figs. 7B-7C show examples for the DSSP-T1SC basis set), the improvement due to inclusion of the side-chain spectra is small. We attribute this unexpected result mainly to two factors. The first is the partial cancellation of contributions from side-chain basis spectra with opposite signs and similar coefficients. This is observed for most reference proteins, because most globular proteins have similar AA compositions. For example, Fig. 7D shows the total contribution of the side chains for each protein in the SP175 reference set, that, overall, are an order of magnitude smaller than the individual side-chain basis spectra. The second factor is the correlation between AA and SS compositions (Pearson coefficients between 0.2 and 0.6 for SP175), which implies that a substantial fraction of the side-chain contributions is described already by the backbone basis spectra.

A potential third factor is that the side-chain contributions strongly depend on their environment, which therefore cannot be accurately described by just one basis spectrum. Examples are contributions from buried versus solvent accessible side chains or side chains in different protonation states. However, we will not explore this possibility further here.

Whereas including the side-chain corrections improves the predicted CD spectra only slightly for most globular proteins, the RMSD_set_ calculated for the GXG20 peptides decreases markedly by 1.9 to 3.2 kMRE, because their CD spectra are largely defined by the side chain signals. This result suggests that the side-chain contributions might be more important for the prediction of CD spectra of small peptides or proteins with unusual AA compositions such as the low complexity regions and sequence repeats often found in intrinsically disordered proteins. Therefore, in the next section, we analyze a system comprised of two disordered proteins to test if side chains indeed play a larger role in predicting their CD spectrum.

### 5.4 Case study: CD predictions for a flexible protein complex

To quantify the outlined improvements in prediction accuracy due to model quality, spectrum rescaling, and side-chain contributions, we selected the flexible protein complex formed by the two IDP domains P53-AD2 and CBP-NCBD^37^ as a test system. These domains form an ordered complex, the structure of which (PDB code 2L14) was determined by NMR spectroscopy. We chose this complex because without the above improvements, a poor mean prediction accuracy (an 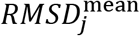 and *σ*_*j*_= 1.58 kMRE) was achieved by the six methods described in Section 3.6, and only the HBSS-3 basis set predicted the CD spectrum with above average accuracy (2.0 kMRE). In this sense, it is a particularly challenging test system.

To put the prediction accuracy improvements in perspective, we first added a ‘Null’ prediction accuracy as the RMSD between the measured spectrum (Fig. 8B, black solid line) and a ‘flat line prediction’ 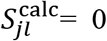 (RMSD= 8.23 kMRE, black dashed line). Then, we predicted the CD spectrum of the P53/CBP complex from the unmodified NMR bundle shown (Fig 8A left, blue), which contained 20 conformations. For the CD predictions, we used the mixed basis set DS-dT3SC, as it has both high accuracy (*RMSD*_cross_ = 3.72 kMRE) and sensitivity (*ΔSS*_*min*_ = 6.9%). The first predicted spectrum (Fig. 8B, blue dashed line) was calculated with only the backbone basis spectra without scaling or side-chain corrections, and differed from the measured spectrum by an RMSD of 7.23 kMRE (Fig. 8C, ‘NMR/Cryst’).

**Figure 8:**
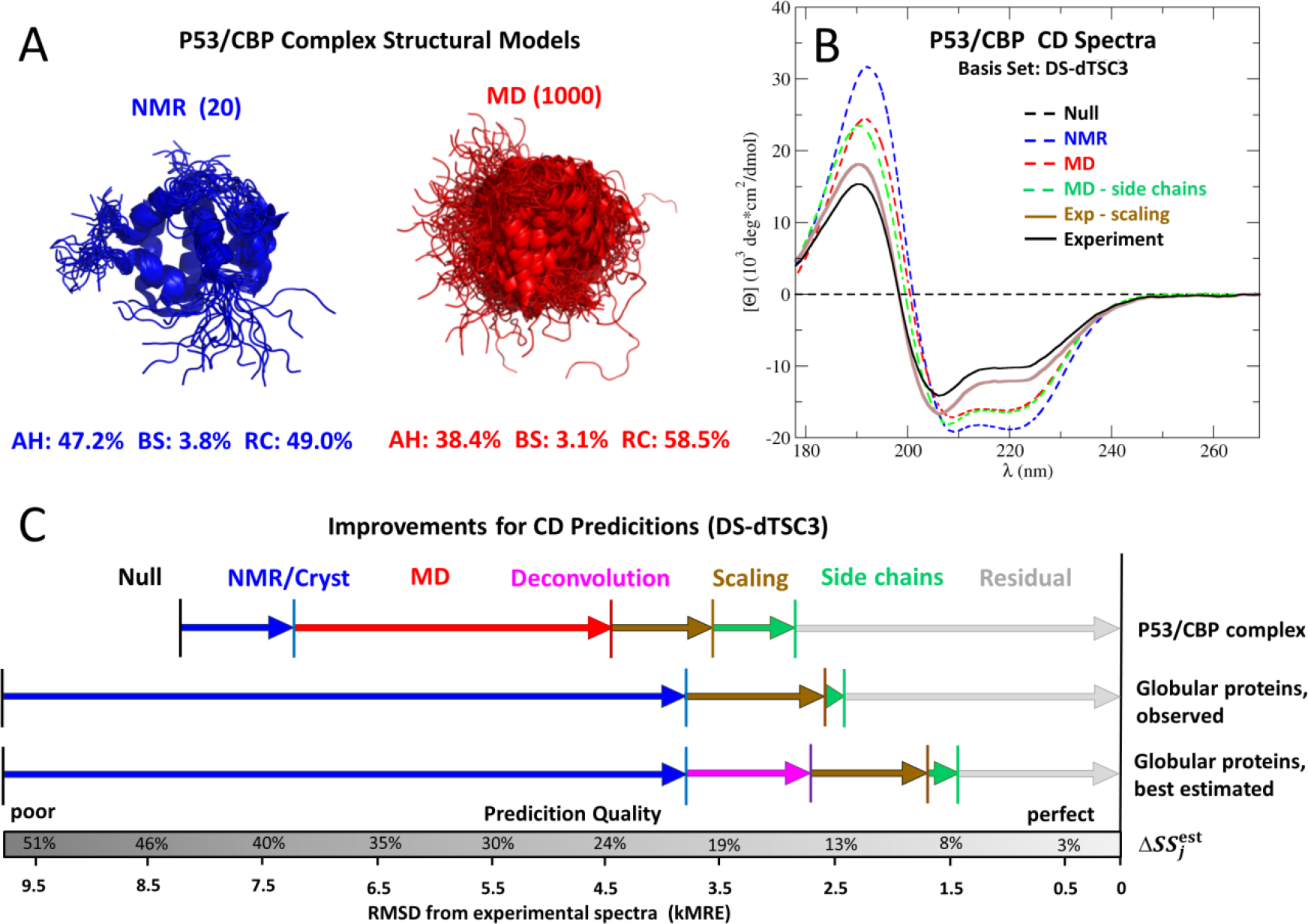
CD spectrum predictions and accuracy improvements for the P53/CBP protein complex. A) Comparison of two structural ensemble models, 20 structures from a nuclear magnetic resonance bundle (‘NMR’) and an ensemble of 1000 snapshots (‘MD’) extracted from a 10 μs MD simulation of the same complex. The percentages below indicate average secondary structure contents, divided into α-helix (AH), β-strand (BS), and random coil (RC). B) Improvement of predicted CD spectra (dashed lines) with respect to measured spectra (solid lines). Predicted spectra are shown as ‘Null’ (black, S^calc^(λ) = 0), ‘NMR’ (blue, using the NMR bundle without any further corrections), ‘MD’ (red, using the MD ensemble instead), and ‘MD -side chains’ (green, MD with side-chain corrections), all calculated with the SESCA basis set DS-dTSC3. The experimental CD spectrum is shown with (brown) and without (black) intensity scaling. C) The sequence of arrows shows, to scale, the increase in prediction accuracy due to the above four improvement steps for the P53/CBP complex (top) and for the SP175 reference set (averaged, mid). The bottom arrows indicate the best achievable accuracies for the SP175 set using ‘perfect’ secondary structure compositions obtained by CD spectrum deconvolution (magenta). The bottom grey bar translates the achieved RMSDs from the measured spectra into estimated secondary structure errors for the used models.

To probe the effects of the underlying structural model on prediction accuracy, we generated a second model for the P53/CBP complex (Fig. 8A right, red). This model was a structural ensemble of 1000 conformations obtained from an MD simulation, which was started from the NMR model as described in Section 3.3. The MD ensemble described conformational heterogeneity and flexibility of the system, and its SS composition contained 9.5% more random coil than the NMR bundle. The MD model was validated by predicting the average backbone NMR chemical shifts from both models using the Sparta+ program ^38^, and by comparing the resulting chemical shift profiles (discussed in Section S12) with the original NMR measurements. The lower average deviation of the calculated C_α_ secondary chemical shifts (1.06 ppm vs. 1.39 ppm) – which are strongly correlated with the SS – suggests that the MD model represents the SS composition of the P53/CBP complex in solution better than the original NMR model.

As expected, the RMSD of the spectrum predicted from the MD model (Fig. 8B, red) is markedly smaller (4.45 kMRE), improving the prediction accuracy by 2.48 kMRE (Fig. 8C, red arrow). Additionally, the better model also improved the 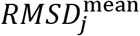 of the complex by 2.0 kMRE when the CD spectra were predicted using the four SESCA basis sets listed in Section 3.6.

Next, we assessed the effect of intensity scaling. To this end, we determined a mean scaling factor 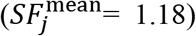 by averaging over the optimal scaling factors required to match the spectra predicted from the MD model using the four basis sets mentioned above. For the DS-dTSC3 basis set, rescaling the measured CD spectrum by 1.18 (Fig. 8B, brown) further reduced the prediction RMSD for the MD model from 4.45 kMRE to 3.59 kMRE (Fig. 8C, brown arrow).

Finally, including side-chain contributions shifts the peaks of the predicted spectrum (Fig. 8B, green) between 190 nm and 210 nm, and reduces the deviation from the rescaled spectrum to *RMSD*_*j*_ = 2.85 kMRE (Fig 8C, green arrow). Combined, the applied corrections improve the prediction accuracy for the P53/CBP complex by 4.1 kMRE (7.23 kMRE vs. 2.85 kMRE), indicating that such corrections may allow accurate SESCA predictions for other IDPs as well.

For comparison, the CD spectrum of P53/CBP complex was also predicted from the NMR model by DichroCalc (6.16 kMRE, without side chains) and PDB2CD (7.79 kMRE). Unfortunately, for technical reasons, predicting the CD spectrum from the MD model was not possible using these methods. Rescaling the measured spectrum improves the RMSD for both methods to 5.03 kMRE and 6.50 kMRE, respectively. Additional accounting for side-chain contributions improves the RMSD further for DichroCalc to a remarkable 3.46 kMRE. Thus, when both scaling and side-chain corrections are applied, the prediction RMSD of DichroCalc is between 2.85 kMRE and 5.62 kMRE, the values obtained using the tested SESCA basis set with and without accounting for the conformational flexibility by MD.

The second series of arrows in Fig. 8C (‘Globular proteins, observed’) shows the corresponding average RMSD improvements achieved for the SP175 reference set using the same DS-dTSC3 basis set. We followed the same steps as outlined for the P53/CBP complex, except that improvements due to MD simulations were not calculated, as validated MD simulations were mostly not available. As indicated by the gray bar at the bottom of Fig. 8C, the average quality of the (mostly) crystal structures in the reference set (*RMSD*_set_ = 3.73 kMRE, 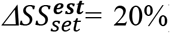) is similar to that obtained for the P53/CBP complex by NMR with subsequent MD simulation 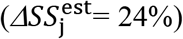. Using these structural models, *RMSD*_set_ improves mostly due to intensity scaling (1.21 kMRE, brown arrow), whilst side-chain corrections – as expected – yield much smaller average improvements (0.16 kMRE, green arrow) compared to the IDP complex (0.74 kMRE). Applying the above corrections also reduces the estimated model errors to 13% for the SP175 set and to 15% for the P53/CBP complex, respectively.

Finally, we also calculated an upper limit for the prediction accuracy that can be achieved with ‘perfect’ structure models, obtained by deconvolving (similar to Sec 5.1) the rescaled and side-chain corrected CD spectra with the backbone basis spectra. Applying all further improvements, as outlined above, results in the third sequence of arrows (Fig. 8C, Globular proteins, best estimated), and an optimal RMSD_set_=1.43 kMRE is achieved.

## Conclusions

In this study we presented a new semi-empirical spectrum calculation approach (SESCA) to predict the electronic circular dichroism (CD) spectra of proteins from their secondary structure (SS) composition, which in turn was derived from their model structures. For several structure classification algorithms (DSSP, DISICL, and HbSS) basis spectrum sets were derived and their prediction accuracies were assessed by comparison to measured CD spectra.

All basis spectra were derived and optimized using a reference set consisting of 71 globular proteins; subsequently, the prediction accuracies of the basis sets were determined by cross-validation on a second, non-overlapping set of eight selected proteins, covering a broad range of SS contents. SESCA predicts the experimental CD spectra of these proteins with an average root-mean-squared deviation (RMSD) as small as 3.2 ± 0.6 × 10^3^ degree·cm^2^/dmol (3.2 ± 0.6 kMRE) in mean residue ellipticity units or 0.9 ± 0.2 M^−1^cm^−1^ in Δε units. This prediction accuracy is markedly better than that of the best currently available algorithms PDB2CD (4.7 kMRE) and DichroCalc (6.1 kMRE).

Closer analysis of the optimized basis sets shows that the best possible prediction accuracy given the current reference set and SS information was reached using 3-8 basis spectra, and that this prediction accuracy is similar for all the tested SS classification protocols.

Further, we demonstrate that SESCA enables one to validate structural models based on the RMSD between its predictions and measured protein CD spectra. Depending on the basis set used, our results show that SESCA is able to discriminate between structural models differing as little as 4-12% in their SS composition. Using this methodology and the measured CD spectra, we estimate that, on average, the SS composition of model structures in our reference sets differs by 15–20% from the solution structure of their respective proteins.

Investigating 13 reference proteins of which the CD spectra were particularly hard to predict, we determined that these SS differences, together with inaccurate normalization of the measured CD spectra are the major factors limiting our prediction accuracy. Nevertheless, even though inaccurate protein models and CD spectra lead to poor RMSDs for some of the reference proteins, our CD spectrum prediction method proved to be relatively robust with respect to these outliers in the training set.

Rescaling the reference spectra, discarding the worst outliers from the training set, and accounting for AA side-chain contributions via an additional 4-7 basis spectra, we derived ‘mixed’ basis sets that improved the prediction accuracy of SESCA to 2.5 ± 0.2 kMRE.

For globular proteins, including side-chain contributions yielded only small (on average, less than 0.2 kMRE) improvements for CD predictions. In contrast, for small peptides as well as for a sample complex of two intrinsically disordered protein (IDP) domains, namely P53-AD2 and CBP-NCBD (P53/CBP), side-chain contributions improved the prediction accuracies substantially.

Due to the simple SS calculations involved, as well as the use of pre-calculated basis sets, SESCA is computationally highly efficient and can be applied to rather large structural ensembles. This feature allows one to go beyond average structures and to account for effects of a protein’s conformational flexibility on its CD spectrum. Indeed, for the P53/CBP complex, an extended molecular dynamics trajectory improves the accuracy of the calculated CD spectrum considerably (by more than 2.0 kMRE).

Specifically, the example of the P53/CBP complex suggests that SESCA may be particularly helpful in modelling IDPs. These biologically highly relevant molecules are notoriously hard to characterize, and structural ensembles are usually required to understand their conformational flexibility, which is often closely related to the IDPs function. A more systematic assessment of SESCA performance regarding IDPs will be addressed in a separate study.

Here, we have exploited the high sensitivity of CD spectroscopy to the average SS of α-L-amino acid polypeptides to evaluate and improve protein structural models. Our SESCA approach is not restricted to proteins and peptides, however. Because CD spectroscopy is also sensitive to the structures of poly-nucleic acids (DNA and RNA), modified polypeptides (e.g., by post-translational modifications or ones comprised of unnatural amino acids), as well as certain carbohydrates (such as glycan structures), SESCA can also be applied to these biomolecules as soon as appropriate reference data sets become available.

A python implementation of our semi-empirical CD calculation method SESCA, as well as basis sets and tools compatible with the SS classification algorithms DISICL and DSSP are publicly available online: **http://www.mpibpc.mpg.de/sesca**.

## Supporting information

All supplementary materials

## Acknowledgements

The authors would like to thank J. Kardos and Cs. Micsonai for prodiving CD spectra for the SP175 protein data set and the BestSel algorithm, to J. Hritz, S. Becker, and C. Griesinger for providing protein samples for the CD measurements, B. Gyurcsik for the aid with the experimental design, and P. Kellers for editing the manuscript. We would like to thank the storage ring facilities of Aarhus University (ISA) for awarding beam time on the AU-CD beam line, ASTRID2, for the measurements of the CD spectra of the short peptides. This research project was funded and supported by the Alexander von Humboldt Foundation and the Max Planck Society.

