## Supplementary material for "SESCA: Predicting Circular Dichroism Spectra from Protein Molecular Structures": All supplementary materials

*Gabor Nagy[1], Maxim Igaev[1], Søren V. Hoffmann[2], Nykola C. Jones[2], Helmut Grubmüller[1*]*

[1]: Department of Theoretical and Computational Biophysics, Max Planck Institute for Biophysical Chemistry, Am Fassberg 11, D-37077 Göttingen, Germany

[2]: ISA, Department of Physics & Astronomy, Aarhus University, Ny Munkegade 120, DK 8000 Aarhus C, Denmark

[*]Corresponding author

Keywords: protein structure, CD spectrum prediction, structure validation, secondary structure

### Supplementary Information:

#### S1 Calculation of basis spectra

The basis spectra $B_i(\lambda)$ are derived using eq. 1 independently for each available wavelength λ from a sufficiently large training set of protein structures and their CD-spectra $S_j(\lambda)$. Because typically the number of basis spectra $F$ is smaller than the number of available training spectra $N$ (here, $F$=1…20 and $N$=64), eq. 1 represents an over-determined linear equation system. The basis spectra that minimize the average RMSD between calculated and measured CD spectra according to eq. 2, where $S_j^{\mathrm{calc}}(\lambda) = \sum_{i=1}^{F} C_{ji} B_i(\lambda)$, are obtained via

$$\boldsymbol{b}(\lambda) = (\boldsymbol{C}^{\boldsymbol{T}} \boldsymbol{C})^{-1} \boldsymbol{C}^{\boldsymbol{T}} \boldsymbol{s}(\lambda). \qquad \text{(S1)}$$

We have used matrix notation for the coefficients $\mathbf{C} = \{C_{ij}\}$ and the vector notation for the basis spectra $\mathbf{b}(\lambda)=\{B_i(\lambda)\}$, and CD spectra $\mathbf{s}(\lambda)=\{S_j(\lambda)\}$, respectively. Figures S24-S26 show basis spectrum sets that were derived by determining the basis spectra for each secondary structure (SS) element in structure classification protocol. This was done by taking the fraction of amino acids in the protein classified to be in structural element $i$ as $C_{ij}$ and applying eq. S1 on the far UV (175-269 nm) wavelength range sampled in 1 nm steps, for all 64 proteins in the TR64 set (see section 3.1).

#### S2 Basis set determination: the 'Hard approach'

For the hard basis set optimization approach (Fig. S1A), our aim was to find basis spectrum sets that provide the most accurate prediction of protein CD spectra. To trade-off some of the fitting accuracy for reduced overfitting, we applied a Monte Carlo (MC) approach with a cross-validation, during the search for the optimal assignments and number of basis spectra. To this aim, the protein reference set was divided into two subsets. The larger subset (training set) was used to derive the basis spectra, and the basis set accuracy was evaluated by the average RMSD of the calculated CD spectra of the smaller subset (evaluation set) according to eq. 2. During each optimization cycle, random changes were applied to the assignment matrix, the corresponding basis spectra for the given assignment were calculated (described in Section S1), and the new assignment was accepted or rejected based on its effect on the obtained basis set accuracy of the evaluation set ($RMSD_{eval}$). At the end of the optimization, the five assignments with the lowest $RMSD_{eval}$ were recorded and the complete reference set was used to fit the basis spectra and obtain the final optimized basis sets.

We imposed two constraints on the assignment factors of the hard basis sets: 1) $\sum_{i=1}^{F} \alpha_{ki} = 1$, and 2) $\alpha_{ki} \in \{1,0\}$. These constraints ensured that the resulting basis spectra are normalized, that there are no overlaps between the structural classes the basis spectra represent, and significantly reduced the search space of the MC algorithm. Initially, the hard optimization procedures were started from a naïve assignment (F=K) for each classification protocol, in which case **A** is the identity matrix ($\alpha_{ki}$ is 1 if $i=j$ and 0 otherwise). However, the basis sets resulting from the first optimization were used as initial guesses for subsequent optimization rounds until convergence was reached both for the number of basis spectra and $RMSD_{eval}$.

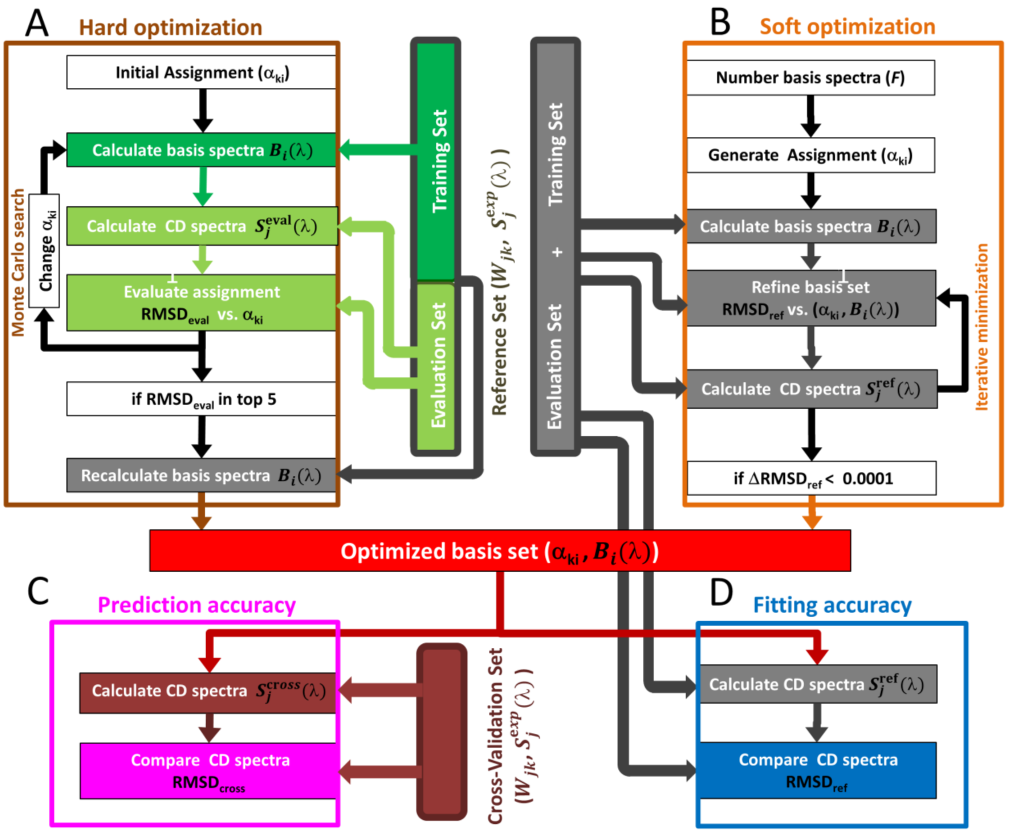


**Figure S1:** Basis set optimization and assessment schemes. The basis sets (shown in red) are derived and optimized either though the hard or the soft optimization approach, using the same reference set of proteins, including the SS information ($W_{jk}$) and CD spectra ($S_j^{\text{exp}}(\lambda)$) of each protein. During the hard optimization (panel A) the reference set was divided into a training set (dark green) and an evaluation set (light green) to perform an 'internal' cross validation during the search for optimal assignments. The undivided reference set (shown as grey boxes and arrows) was used during the soft optimization (panel B) as well as at the end of the hard optimization to calculate basis spectra for the best assignments. The same undivided reference set was used to assess the fitting accuracy (panel D) of the optimized basis set (regardless of the optimization method). In contrast, during the assessment of the prediction accuracy (panel C), a different set of proteins (shown in dark red) were used for cross-validating the predictive power of the optimized basis sets.

During each hard optimization step, a random change was introduced to the assignment matrix **A**, by reassigning one of the SS elements to another structural class. Then, the basis spectra $B_i(\lambda)$ were recalculated and the average deviation ($RMSD_{eval}$) from the experimental CD spectra was computed for the evaluation set both before and after the change was applied. If $e^{-\beta*(\Delta RMSD_{eval})}$ was larger than a randomly generated number between 0 and 1, the new assignment was accepted, otherwise rejected. In the next optimization step, a new random change was applied to the last accepted assignment. The acceptance ratio in this notation was controlled by $\beta$, the strictness parameter determining how often changes with an unfavourable $\Delta RMSD_{eval}$ are accepted. By default, $\beta = 8.0$ was applied to optimizations, which was lowered (down to 1.0) if the acceptance rate in an optimization dropped below 20%. Accepted assignments with the lowest five $RMSD_{eval}$ during the MC search were saved and used to calculate the basis spectra of optimized basis sets.

We note that the level of coarse graining for the SS information is given by the assignment matrix $\alpha_{ik}$. Extreme cases are (a) combining all SS elements provided by the particular SS classification protocol in use into $F$=1 class, and (b) into $F$=$K$ classes. In case (a), only very little (likely too little) information is retained – typically the α-helical content – whereas in the 'naive' case (b), the full SS information is provided with the possible risk of overfitting. Therefore, subsequent cross validation is crucial for determining the optimal level of coarse graining.

The search space for the hard optimization contains $F^K$ possible **A** matrices, where $F$ is the number of structural classes/basis spectra and $K$ is the number of the SS elements. For example, assigning five structural elements to three classes defines a search space of $3^5 = 243$ assignments, whilst 19 structural elements assigned to 10 classes result in a search space of $10^{19}$. When optimizing small basis sets with 5-8 SS elements, a single optimization process with 500 accepted moves was usually sufficient to completely explore the search space, often visiting the global optimum of the assignment space multiple times. In the case of more than 10 structural elements, several 10000-step optimizations were started from multiple initial assignments described below. In these cases, assignments resulting from the initial optimization procedure were used to start new parallel processes to more effectively explore the search space. To further increase the efficiency of the hard optimization, important SS elements – such as the α-helix and at least one of β-strand elements – were assigned to different classes and then excluded from being reassigned (effectively decreasing K). In addition, if the move resulted in a more favourable $RMSD_{eval}$, both structural classes with no assigned SS elements and the SS elements themselves could be temporarily eliminated from the basis set. Eliminated classes and SS elements could be reintroduced to the basis set through random changes during the same optimization process, and missing SS elements were reintroduced between subsequent optimization processes to conserve the normalization of basis spectra. We have performed several optimization processes for each SS classification protocol, until the number of basis spectra in the best optimized basis sets stabilized, and $RMSD_{ref}$ values similar to the soft basis sets of the same basis set size were reached (described in Section S4).

**Addenum: Initial basis sets.** Three deconvolution basis sets (Figs. S23-S25) were used to assess the applicability of our method without extensive optimization. The first basis set was determined by Sreerama and Woody (Set_Sreer-1) [1], contains six basis spectra (regular helix, irregular helix, regular strand, irregular strand, poly-proline helix, and disordered). ). The second basis set, (Set_Perczel-1) was derived by Hollósi and Perczel [2] and contains five basis spectra (α-helix. β-strand, Turn type I/III, unordered, and other contributions). Finally, the third basis set

(Set_BestSel-2) was derived for the BestSel program by Micsonai and Kardos [3], with eight basis spectra (regular helix, irregular helix, left-handed anti-parallel, relaxed anti-parallel, and right-handed anti parallel β-strands, parallel β-strand, turn structures, and others). For each of these basis spectra, SS elements from the structure classification algorithms (DSSP and DISICL for the first two and DISICL and HbSS for the third) were assigned based on the description of the basis set in their original publications. Once the assignment was complete, the CD spectra for the proteins of the TS8, EV9, TR64 and SP175 sets were calculated using the SS content of their crystal structure and were compared to the experimental spectra.

Furthermore, we derived naive basis sets for the classification algorithms (Figs. S26-S30) DSSP (Set_DSSP-F), simplified and detailed DISICL (Set_DS-simF and Set_DS-detF, respectively), normal and extended HbSS (Set_HBSS-F and Set_HBSS-E) and the deconvolution algorithm BestSel (Set_Bestsel-der, Fig. S31). These basis sets contained one basis spectrum for each of the algorithm's SS elements, and the SP175 data set was used as a reference set to calculate their basis spectra. These basis sets were used as initial guesses for the hard and soft optimization procedures.

#### S3Calculating side-chain contributions

The individual side-chain contributions were estimated from the CD spectra of the MP79 reference set. First, the SS contributions were calculated using four optimized basis sets (DS5-4, DS-dT, DSSP-1 or DSSP-T, see the Table S8 for further details on these basis sets). Then, the resulting predicted contributions were averaged and subtracted from the rescaled (see Section 5.2) measured CD spectra. These 'secondary-structure-free' CD spectra were used in eq. S1 with the amino acid (AA) composition of the proteins and peptides as coefficients to derive one basis spectrum for each AA side chain. We also derived basis sets with more simplified representations of the side-chain contributions. These ‘mixed basis sets’ were derived from the MP79 reference set in three steps. First, the SS contributions were calculated and subtracted from the CD spectra. Second, basis spectra for the side chains were derived and optimized using the AA composition and the secondary-structure free CD spectra of the reference proteins. Third, the side-chain contributions were calculated and subtracted from the experimental CD spectra, and the resulting ‘side-chain free’ CD spectra were used to recalculate the basis spectra for the SS-dependent backbone contributions.

The optimization of the side-chain and backbone basis spectra was performed by the hard optimization scheme separately with the following modifications. Before the optimization, the MP79 reference set was separated into six subsets (each containing 13 or 14 proteins). In each optimization step, after the SS elements / AAs were grouped and assigned to basis spectra, one of the MP79 subsets was designated as the evaluation set, whilst the rest of the reference proteins were used to derive the basis spectra (as a training set). The derived basis spectra were used to calculate the CD spectra of the evaluation set. This process was repeated six times such that each of the subsets was predicted once from the rest of the MP79 reference set. After calculating each of the evaluation subsets, their RMSD was averaged and used as $RMSD_{eval}$ to determine if the assignment is accepted or rejected. The optimization process was continued until 250 - 5000 accepted moves were reached (depending on the basis set size), with the five best assignments recorded for further use. Basis spectra for the recorded assignments were recalculated from the full MP79 reference set. These finalized basis spectra were used to predict the ‘secondary-structure free’ or ‘side-chain free’ CD spectra of the TS8 protein set as cross validation. The combination of side-chain and backbone basis spectra that predicted the TS8 protein set with lowest $RMSD_{cross}$ were combined into mixed basis sets. Then, mixed basis sets were used to calculate the CD spectra of the SP175, GXG20, GP59, and TS8 data sets, so that they can be compared with the SS-only optimized basis sets, and predictions from PDB2CD and DichroCalc.

#### S4 Basis set determination: The 'soft approach'

The hard optimization scheme introduced above is limited to a restricted assignment factor space ($\alpha_{ki} \in \{0,1\}$) and, therefore, it should be possible to further improve the accuracy of reconstructing the CD spectra from the SS information by removing this limitation. Accordingly, in our more general soft optimization approach, the assignment factors can be any real number ($\alpha_{ki} \in R$). During the soft optimization, we simultaneously derived the basis spectra and assignment factors that most accurately reproduced the CD spectra of the reference protein data set (best fitting accuracy). Consequently, besides the spectral and structural information of the reference data set, only the desired number of basis spectra is specified for the soft optimization, and no 'internal' cross-validation is required to trade-off the accuracy of the fit for an improved general predictive power. To obtain the optimal basis sets, the non-linear equation system defined by eq. 4 has to be solved simultaneously for all wavelengths of each protein spectrum in the reference data set. In matrix notation, this optimization problem is

$$||\boldsymbol{W}\,\boldsymbol{A}\,\boldsymbol{B}-\boldsymbol{S}||^2 \stackrel{!}{=} \min, \qquad \text{(S2)}$$

where $\mathbf{S}=\{S_{jl}\}$ and $\mathbf{W}=\{W_{jk}\}$ are the matrices containing the spectral and structural information of the reference set, respectively, and the matrix $\mathbf{B}=\{B_{il}\}$ describes the basis spectra. The matrix elements $S_{jl}$ and $B_{il}$ are obtained by discretizing the experimental CD spectra $S_j(\lambda)$ and basis spectra $B_i(\lambda)$ at L wavelengths. This optimization problem is solved simultaneously for the matrices **A** and **B** by setting their element-wise matrix derivatives to zero:

$$\frac{\partial}{\partial \boldsymbol{A}} \mathrm{tr}[(\boldsymbol{W}\,\boldsymbol{A}\,\boldsymbol{B}-\boldsymbol{S})^T\ (\boldsymbol{W}\,\boldsymbol{A}\,\boldsymbol{B}-\boldsymbol{S})] =$$

$$2\,\boldsymbol{B}\,\boldsymbol{B}^{\mathrm{T}}\,\boldsymbol{A}^{\mathrm{T}}\,\boldsymbol{W}^{\mathrm{T}}\,\boldsymbol{W} - 2\,\boldsymbol{B}\,\boldsymbol{S}^{\mathrm{T}}\,\boldsymbol{W} \overset{!}{=} 0 \qquad \text{(S3)}$$

$$\frac{\partial}{\partial \boldsymbol{B}} \mathrm{tr}[(\boldsymbol{W}\,\boldsymbol{A}\,\boldsymbol{B}-\boldsymbol{S})^T\ (\boldsymbol{W}\,\boldsymbol{A}\,\boldsymbol{B}-\boldsymbol{S})] =$$

$$2\,\boldsymbol{B}^{\mathrm{T}}\,\boldsymbol{A}^{\mathrm{T}}\,\boldsymbol{W}^{\mathrm{T}}\,\boldsymbol{W}\,\boldsymbol{A} - 2\,\boldsymbol{S}^{\mathrm{T}}\,\boldsymbol{W}\,\boldsymbol{A} \overset{!}{=} 0 \qquad \text{(S4)}$$

which, yields two coupled non-linear matrix equations

$$\boldsymbol{A} = (\boldsymbol{W}^{\mathrm{T}}\,\boldsymbol{W})^{-1}\,\boldsymbol{W}^{\mathrm{T}}\,\boldsymbol{S}\,\boldsymbol{B}^{\mathrm{T}}\,(\boldsymbol{B}^{\mathrm{T}}\,\boldsymbol{B})^{-1} \qquad \text{(S5)}$$

and

$$\boldsymbol{B} = (\boldsymbol{A}^{\mathrm{T}}\,\boldsymbol{W}^{\mathrm{T}}\,\boldsymbol{W}\,\boldsymbol{A})^{-1}\,\boldsymbol{A}^{\mathrm{T}}\,\boldsymbol{W}^{\mathrm{T}}\,\boldsymbol{S} \qquad \text{(S6)}$$

Equations S5 and S6 are solved iteratively, starting from a random generated matrix **A** ($0.0 \leq \alpha_{ki} \leq 1.0$) to obtain an initial **B** via eq. S6, which is inserted into eq. S5 to obtain an improved **A**, repeated until convergence. A summary of the soft optimization scheme is shown in Fig. S1.

This soft optimization procedure was systematically repeated for each SS classification protocol K times to obtain optimized basis sets with 1-K basis spectra (K being the number of SS elements in the classification protocol). These series of basis sets determine the best fitting accuracy as the function basis set size and SS classification. For each optimization procedure, the convergence criterion was to reach less than $\Delta\mathrm{RMSD}_{\mathrm{fit}}$ = 0.0001 x $10^3$ deg $\mathrm{cm}^2$/dmol change between iterations.

Finally, we note that the hard combination of SS elements is a special case of the more general soft combination approach and therefore, one might expect the latter to yield more accurate calculated spectra for the reference proteins from the same amount of structural information. Because in the soft optimization approach the assignment factors $\alpha_{ki}$ can adopt any real number without further constraints, eq. 3 yields linear combinations of the SS fractions $W_{kj}$. Hence, each basis spectrum $B_i(\lambda)$ can be understood as a 'collective' SS class, such as '0.3 α-helical + 0.7 β-sheet'. Of course, the collective SS classes introduce another layer of complexity to the optimization problem, and therefore increase the chances of overfitting the basis spectra.

#### S5 The analysis of spectral components

The overall accuracy of our method is limited by two factors, first, the information content of the SS composition, and second, the applicability of linear combinations of basis spectra in approximating the experimental CD spectra. The first factor was addressed by our soft optimization approach (Section S4). The second factor determines an upper limit for the fitting accuracy (lowest $RMSD_{fit}$) given a set of reference CD spectra and the number of basis spectra used. To this end, we carried out a principal component analysis (PCA) on CD spectra of the SP175 reference set (see Section 3.1). PCA is a mathematical method to describe a (multidimensional) data set of N members by a basis set of N orthogonal principal component (PC) vectors. How much the data points differ from the average of the set (the variance of the data set) along a PC vector is quantified by its eigenvalue. It is possible to describe a data set with just a few ($F$) PC vectors of the highest eigenvalues (dimensionality reduction)[32], which – by construction – retains the maximum possible variance of the data set, and consequently, provides the reconstruction with the smallest possible deviation. Here, we used PCA to describe the reference CD spectra (a set of $L$ dimensional data points) by basis sets constructed from 1-10

PC vectors of the highest eigenvalues. The basis spectrum coefficients ($C_{ij}$) of the protein $j$ for these basis sets were defined as the projection of the CD spectrum along the particular PC vector $i$ (described below). Note that this analysis is based solely on CD spectra of the reference data set, and does not account for any possible source of inaccuracy related to structure, SS calculations, or scaling errors within the reference set.

During the reconstruction, the PC vectors obtained from PCA were described by the matrix $\mathbf{V}=\{V_{pl}\}$, where the indices $p$ (1….P) and $l$ (1….L) stand for the principal component (in order of their eigenvalue) and wavelength, respectively. In our case, each $\mathbf{v}_{\mathrm{rp}}$ row vector of the matrix $\mathbf{V}$ is one of the discretized PC vectors. The spectra of a reference protein data set were reconstructed using the first P={1-10} principal components

$$S_{jl} = S_l^{ave} + \sum_{p=1}^{P} C_{jp}\, V_{pl}, \qquad \text{(S7)}$$

where $S_{jl}$ is the circular dichroism of the $j^{\mathrm{th}}$ reconstructed protein spectrum at the wavelength $l$, $C_{jp}$ is the projection of that spectrum along the PC vector $p$, $V_{pl}$ and $S_l^{ave}$ are the value of the PC vector and the average CD signal of the data set at wavelength $l$, respectively. The projection of spectrum $j$ along the principal component $p$ can be calculated by taking the scalar product of the normalized spectrum and the PC vector

$$C_{jp} = (\boldsymbol{s}_{rj} - \boldsymbol{s}^{ave})^T\ \boldsymbol{v}_{rp}. \qquad \text{(S8)}$$

The vector $\boldsymbol{s}^{\mathrm{ave}} = \{S_l^{\mathrm{ave}}\}$ is the averaged CD spectrum of the data set.

The projections along the PC vectors are analogous to the basis spectrum coefficients. Therefore, Pearson correlation ($R_{pearson}$) between the SS composition, AA composition, and the projections were calculated for the proteins in the SP175 reference set to estimate the importance of these structural descriptors in calculating the CD spectra. The Pearson correlation between these descriptors were calculated according to

$$R_{\text{pearson}} = \frac{\sum(X_j-\bar{X})\cdot(Y_j-\bar{Y})}{\sqrt{\sum(X_j-\bar{X})^2}\cdot\sqrt{\sum(Y_j-\bar{Y})^2}}, \qquad \text{(S9)}$$

where $X_j$ and $Y_j$ are either the fraction of an AA, the fraction of residues classified as a SS element, or the projection of the CD spectrum along a principal component for the protein $j$, whilst $\bar{X}$ and $\bar{Y}$are the calculated averages for the whole reference set.

#### S6 Describing the error of CD spectrum predictions

Here, we separate the errors of our CD spectrum predictions into an error term that depends on the SS composition of the underlying protein structural model (SS-dependent error), and an SS-independent error term that results from inaccuracies of the prediction method itself. This separation allows us to determine the typical prediction error of SESCA basis sets, and to estimate the error of the underlying structural model based on the accuracy of its predicted CD spectrum.

To this end, we assume that the true SS composition of the solution structure is described by the coefficients $C_{ji}^0$. The CD spectrum predicted from this correct composition deviates slightly from the experimental spectrum (quantified by $RMSD_j^0$), due to inaccuracies in our basis spectra ($B_{il}$), unaccounted contributions to the CD spectrum (e.g. from co-factors and side chains), and potential measurement errors of the spectrum itself. After accounting for all the errors independent of the SS composition at every wavelength ($\Delta S_{jl}$) the experimentally measured CD spectrum is expressed as

$$S_{jl}^{\text{exp}} = \sum_{i=1}^{F} C_{ji}^{0} \cdot B_{il} + \Delta S_{jl}. \qquad \text{(S10)}$$

If we predict the measured CD spectrum from a structural model with the SS composition $C_{ji}$, the RMSD of the predicted spectrum ($S_{jl}^{\text{calc}} = \sum_{i=1}^{F} C_{ji} \cdot B_{il}$) will be

$$RMSD_j = \sqrt{\frac{\sum_{l=1}^{L}\left(S_{jl}^{\text{exp}} - S_{jl}^{\text{calc}}\right)^2}{L}} = \sqrt{\frac{\sum_{l=1}^{L}\left(M_{jl} + \Delta S_{jl}\right)^2}{L}}, \qquad \text{(S11)}$$

where $M_{jl} = \sum_{i=1}^{F} \Delta C_{ji} \cdot B_{il}$ describes the SS-dependent error of the predicted spectrum at wavelength $l$, $\Delta C_{ji} = C_{ji}^{0} - C_{ji}$ is the structural model's error in the coefficient of SS-class $i$, and $B_{il}$ is the CD intensity of for the basis spectrum of SS-class $i$ at wavelength $l$.

Equation S11 can be rewritten using vector notation, considering the total error of the predicted spectrum for protein $j$ at each wavelength $l$ to be described by the vector $\vec{\boldsymbol{R}} = \vec{\boldsymbol{M}} + \vec{\boldsymbol{S}}$. Here, $\vec{\boldsymbol{M}} = \{M_{j1};\ M_{j2}; \ldots\ M_{jL}\}$ is the wavelength vector containing SS-dependent error terms, whereas $\vec{\boldsymbol{S}} = \{\Delta S_{j1};\ \Delta S_{j2}; \ldots\ \Delta S_{jL}\}$ is the wavelength vector of the SS-independent error terms for protein

$j$, respectively. We note that the lengths of these vectors are connected via $|\vec{\boldsymbol{R}}|^2 = |\vec{\boldsymbol{M}}|^2 + |\vec{\boldsymbol{S}}|^2 + 2 \cdot |\vec{\boldsymbol{M}}| \cdot |\vec{\boldsymbol{S}}| \cdot \cos\varphi$, where $\cos\varphi$ is the angle between the vectors $\vec{\boldsymbol{M}}$ and $\vec{\boldsymbol{S}}$. Because $RMSD_j$ is the average root-mean-squared deviation from the measured spectrum, $RMSD_j{}^2 = \frac{1}{L} \cdot |\vec{\boldsymbol{R}}|^2$. Combining these two equations, $RMSD_j$ is written as

$$RMSD_j{}^2 = M_j{}^2 + RMSD_j^{0\,2} + 2 \cdot M_j \cdot RMSD_j^0 \cdot \cos\varphi, \qquad \text{(S12)}$$

where $M_j = \frac{|\vec{\boldsymbol{M}}|}{\sqrt{L}} = \sqrt{\frac{\sum_{l=1}^{L} M_{jl}{}^2}{L}}$ is the average SS-dependent error, and $RMSD_j^0 = \frac{|\vec{\boldsymbol{S}}|}{\sqrt{L}} = \sqrt{\frac{\sum_{l=1}^{L} \Delta S_{jl}{}^2}{L}}$ is the average SS-independent error. We also note that $RMSD_j^0$ is equal to the RMSD of the spectrum predicted from the correct SS composition ($C_{ji}^0$).

Equation S12 allows us to separate the SS-dependent error from the SS-independent error, and to quantify the error in the SS composition of any protein model based on $RMSD_j$ of its predicted spectrum. To this end, we introduce the total error in the model's secondary structure composition as $\Delta SS_j = \sum_{i=1}^{F} |\Delta C_{ji}|/2$, which is equal to the (minimum) fraction of residues in the protein model with incorrect SS classification. We note that any misclassified residue in the model causes a positive (in its current class) and a negative (in its supposed class) error in the coefficients, and thus, the sum of $|\Delta C_{ji}|$ was divided by two to obtain $\Delta SS_j$. The SS-dependent error at wavelength $l$ can be written as $M_{jl} = \left(\sum_{i=1}^{F} \frac{\Delta C_{ji} \cdot B_{il}}{\Delta SS_j}\right) \cdot \Delta SS_j$, where $\Delta SS_j$ gives the magnitude of the structural model's error, and the ratios $\frac{\Delta C_{ji}}{\Delta SS_j}$ describe how $\Delta SS_j$ is distributed between the coefficients of the SS classes. This allows one to express the proportionality between the SS-dependent error and the total model error as

$$M_j = \left( \sqrt{\frac{1}{L} \cdot \sum_{l=1}^{L} \left( \sum_{i=1}^{F} \frac{\Delta C_{ji} \cdot B_{il}}{\Delta SS_j} \right)^2} \right) \cdot \Delta SS_j = m_j \cdot \Delta SS_j \; . \qquad \text{(S13)}$$

Note that, for all structural models with the same $\frac{\Delta C_{ji}}{\Delta SS_j}$ ratios, the SS-dependent error will be a linear function of $\Delta SS_j$. Combining eqs. S12 and S13 results in a second order equation which can be solved for $\Delta SS_j$ to express the error in the structural model

$$\Delta SS_j = \frac{\sqrt{RMSD_j{}^2 + RMSD_j^0{}^2 \cdot (\cos^2\varphi - 1)}}{m_j} - \frac{RMSD_j^0 \cdot \cos\varphi}{m_j}. \qquad \text{(S15)}$$

Equation S15 shows that, for a given $m_j$, $RMSD_j$, and $RMSD_j^0$ value $\Delta SS_j$ will be the largest if the two error vectors are anti-parallel ($\cos\varphi = -1$), in which case $\Delta SS_j = \left(RMSD_j + RMSD_j^0\right)/m_j$, and smallest if the error vectors are parallel ($\cos\varphi = 1$), resulting in $\Delta SS_j = \left(RMSD_j - RMSD_j^0\right)/m_j$. However, if the SS-independent error contains only random noise and no systematic errors, the two error terms should be statistically independent and $\cos\varphi \approx 0$. In this case, eq. S15 is simplified to $\Delta SS_j = \sqrt{RMSD_j{}^2 - RMSD_j^0{}^2}/m_j$, which approaches a linear function $\Delta SS_j = RMSD_j/m_j$ for $RMSD_j{}^2 \gg RMSD_j^0{}^2$.

The obtained equations for the three special cases ($\cos\varphi = \{0, +1, -1\}$ will be used in Section S8 to determine the best estimate for $\Delta SS_j$ assuming that the SS-dependent and SS-independent errors of the predicted spectrum are statistically independent ($\cos\varphi = 0$), but providing the upper and lower bounds for the total model error if the assumption is incorrect.

#### S7 CD spectrum deconvolution

Spectrum deconvolution is a method to decompose a measured CD spectrum of a target protein into a linear combination of basis spectra (see eq. 1 in the main text). This procedure is often used to determine the SS composition ($C_{ji}$) that best describes the solution structure of a target protein. For a given set of basis spectra ($B_{il}$), the SS composition that fits the spectrum best minimizes the average deviation ($\mathrm{RMSD}_j$) between the decomposed ($S_{jl}^{dec} = \sum_{i=1}^{F} C_{ji} \cdot B_{il}$) and the measured spectrum ($S_{jl}^{\exp}$). Therefore, to find the optimal $C_{ji}$ we minimized the function

$$F(C_{ji}) = \sqrt{\frac{1}{L}\sum_{l=1}^{L}\left(S_{jl}^{\exp} - \sum_{i=1}^{F} C_{ji} \cdot B_{il}\right)^2} + \Lambda \cdot \left|1 - \sum_{i=1}^{F} C_{ji}\right| + \Lambda \cdot \sum_{i=1}^{F} \delta(C_{ji}), \qquad \text{(S16)}$$

where the first term is the RMSD of the decomposed spectrum, whilst the second and third terms ensure that $C_{ji}$ add up to 1 and remain positive, respectively. Note that Λ is a constant Lagrange multiplier in this equation, whereas the function $\delta(C_{ji}) = -C_{ji}$ if the coefficient is negative and 0 otherwise.

Our minimizations were carried out using an adaptive simplex minimizer (Nelder-Mead) [4] algorithm from the scientific python package. The minimizations were continued until 5000 iterations or until the convergence criterion $\Delta F(C_{ji}) < 10^{-15}$ was reached. To find the global RMSD minimum of our reference proteins, first 10 minimizations were started from random initial SS compositions, and minimization result with the lowest RMSD was taken. If none of minimizations reached convergence for a protein, a second round of 50 minimizations with

50 000 iterations were launched. If still no minimum was found, then a single, 50 000 step minimization was started from the SS composition of the proteins crystal structure, and the SS composition with the lowest RMSD found was used as the best one.

#### S8 Deriving error models for structure validation

Here, we provide and test error models that estimate the total error ($\Delta SS_j$) in the SS composition ($C_{ji}$) of a protein structural model based on the error ($\mathrm{RMSD_j}$) of its predicted CD spectrum. The equations described in Section S6 allow one to exactly calculate $\Delta SS_j$ from $RMSD_j$, provided that we know the parameters $m_j$, $RMSD_j^0$, and $\cos\varphi$. In summary, $m_j$ is the slope that connects the average SS-dependent error ($M_j$) of the predicted spectrum with the error of the structural model $\Delta SS_j$, $RMSD_j^0$ describes the average SS-independent error of the predicted spectrum, and $\cos\varphi$ describes the non-additivity between the two error terms.

Unfortunately to calculate these parameters, knowing the correct SS composition ($C_{ji}^0$) of protein $j$ under the conditions CD spectrum (which is used for validating the structural model) was measured is essential. Because $C_{ji}^0$ is typically unknown, we determine average parameters that allow the estimation of $\Delta SS_j$ as accurately as possible.

To determine the best $\Delta SS_j$ estimation method we 1) approximate $C_{ji}^0$ and $RMSD_j^0$ for each reference protein of the SP175 set by spectrum deconvolution (see Section S7), 2) compute $RMSD_j$ and $\Delta SS_j$ (compared to $C_{ji}^0$) for each reference structure in the SP175 set using several basis sets, 3) fit the parameters for the $\Delta SS_j$ estimation (described below) to the obtained $RMSD_j$

and $\Delta SS_j$ values, 4) evaluate the error models by comparing the estimated $\Delta SS_j$ to the $\Delta SS_j$ obtained from deconvolution, and 5) determine the error margins for our estimated $\Delta SS_j$ values.

We note that if the measured CD spectrum and the basis set contain no severe systematic errors, $RMSD_j^0$ is the minimum of the RMSD($C_{ji}$) landscape. In the deconvolution algorithm described in Section S7 we approximate $C_{ji}^0$ and $RMSD_j^0$ for the SP175 reference proteins by minimizing $RMSD_j$ as function of $C_{ji}$. We used the differences between the approximated $C_{ji}^0$ and $C_{ji}$ of the reference structures to compute $\Delta SS_j$ for the SP175 proteins, and after predicting the reference spectra from $C_{ji}$, to obtain 71 $RMSD_j$ vs. $\Delta SS_j$ values for several basis sets.

For each of these basis sets the resulting data points in ${RMSD_j}^2$ vs. ${\Delta SS_j}^2$ space were fitted with three equations given in the Fig. S2 (bottom) using orthogonal distance regression [5] (ODR). These equations − derived from eq. S15 (Section S6) using different assumptions − yielded similar fit parameters within the estimated uncertainties ($\sigma_m, \sigma_R$), including a fitted value of $\cos\varphi_f$ = 0±10$^{-5}$ for equation C, which resulted in eqs. B and C giving virtually identical solutions for all tested basis sets. For simplicity, in the table in Fig. S2 we only show fit parameters obtained from eq. B for each basis set, and used for our final error model.

Next, as illustrated in Fig S2 (left) for the DS-dT basis set, we compare the estimated $\Delta SS_j^{\mathrm{est}}$ values (solid colored lines) with the correct $\Delta SS_j$ values (black cross symbols) to determine the best estimate. To quantify the goodness of the fits, we computed $\chi^2 = \sum_{j=1}^{N}\left({\Delta SS_j}^2 - {\Delta SS_j^{\mathrm{est}}}^2\right)^2/(N-P)$, where $N$=71 is the number of reference proteins, and $P$={1,2,3} is the number of fit parameters for eqs. A, B, and C, respectively. The obtained average $\chi^2$ values indicate that eq. A describes the SP175 reference set slightly better than eqs. B and C do.

Additionally, eqs. B and C do not yield real solutions $\Delta SS_j^{est}$ for $RMSD_j < R_f$, in which cases $\Delta SS_j^{est} = 0$ is assumed. This limitation further increases the average deviation between the real and estimated $\Delta SS_j$ as shown by the tabulated $\Delta\Delta SS = \sqrt{\frac{1}{N}\sum_{j=1}^{N}(\Delta SS_j - \Delta SS_j^{\mathrm{est}})^2}$ values in Fig. S2. Therefore, we chose the best estimate to be $\Delta SS_j^{est} = RMSD_j\,/m_f$ for our error models.

Finally, we defined two error margins for $\Delta SS_j^{est}$. The more narrow error margin (purple dashed line) is defined by the average error of the best estimate as $\Delta SS_j^{est} \pm \Delta\Delta SS$, and contains 73% of the data points on average. The broader error margin (green dashed line) is defined based on eq. S15, assuming the fitted average $m_j$ and $RMSD_j^0$ parameters ($m_f$ and $R_f$, respectively), and strong correlations between the error terms ($\cos\varphi = \pm 1$). In this case the upper and lower bounds of $\Delta SS_j^{\mathrm{est}}$ are $\Delta SS_j^{\mathrm{up}} = \left(RMSD_j + (R_f + \sigma_R)\right)/\left(m_f - \sigma_m\right)$ and $\Delta SS_j^{\mathrm{low}} = \left(RMSD_j - (R_f + \sigma_R)\right)/\left(m_f + \sigma_m\right)$, and contain ~94% of reference proteins on average.

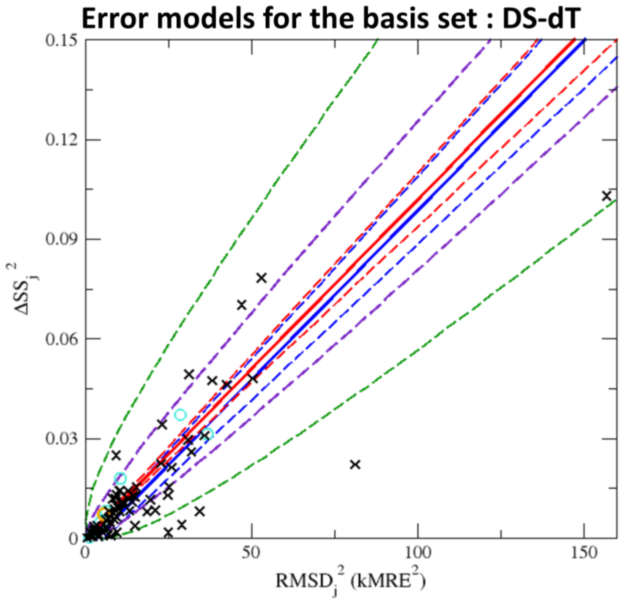


#### Fit Summary

| Basis set | DS-dT | DSSP-1 | HBSS-3 | DS5-4 | Average |
|---|---|---|---|---|---|
| $m_f$ (kMRE) | 31.3 | 18.2 | 17.3 | 15.3 | 21.0 |
| $\sigma_m$ (kMRE) | 1.3 | 2.0 | 1.0 | 1.8 | 1.5 |
| $R_f$ (kMRE) | 1.79 | 1.08 | 0.99 | 2.25 | 1.52 |
| $\sigma_R$ (kMRE) | 0.54 | 0.34 | 1.40 | 0.75 | 0.76 |
| $\chi^2$-A ($10^{-4}$) | 12.67 | 19.35 | 15.99 | 19.65 | 16.91 |
| $\chi^2$-B ($10^{-4}$) | 12.59 | 19.50 | 16.20 | 19.62 | 16.98 |
| $\chi^2$-C ($10^{-4}$) | 12.78 | 19.79 | 16.44 | 19.91 | 17.23 |
| ΔΔSS-A (%) | 2.6 | 5.4 | 6.3 | 7.9 | 5.5 |
| ΔΔSS-B (%) | 2.97 | 5.61 | 6.62 | 9.74 | 6.2 |
| ΔΔSS-C (%) | 2.97 | 5.61 | 6.62 | 9.74 | 6.2 |
| Bounds (%) | 74.6 | 70.4 | 81.7 | 64.8 | 72.9 |
| Bounds (%) | 94.4 | 90.1 | 90.1 | 100 | 93.7 |

| Fit Equations: | Assumptions: |
|---|---|
| Estimate A: $\Delta SS_j^{\,2} = RMSD_j^{\,2} / m_f^{\,2}$ | $\cos\Theta = 0,\ R_f \ll RMSD_j$ |
| Estimate B: $\Delta SS_j^{\,2} = (RMSD_j^{\,2} - R_f^{\,2}) / m_f^{\,2}$ | $\cos\Theta = 0$ |
| Estimate C: $\Delta SS_j^{\,2} = \left(\sqrt{RMSD_j^{\,2} + R_f^{\,2}\cdot(\cos\Theta^2 - 1)} - R_f\cdot\cos\Theta\right)^2 / m_f^{\,2}$ | none |

**Figure S2:** Comparison between three different estimates for the total error in the SS composition ($\Delta SS_j$) of protein structural models. The estimation of $\Delta SS_j$ is based on the deviation ($RMSD_j$) between the measured CD spectrum and spectrum predicted from the structural model. The error estimates (A-C) with three functional forms (at the bottom) were fitted to the SP175 reference proteins shown as black cross symbols on the left side of the figure. The best estimates (shown as solid lines) with estimated error margins (shown as dashed lines, see section S8) are also displayed for the DS-dT basis set. Obtained fit parameters ($m_f, \sigma_m, R_f, \sigma_R$), goodness of fit measures ($\chi^2, \Delta\Delta SS$) and the percentage of data points contained within the selected error margins (Bounds) for four tested basis sets (DS-dT, DSSP-1, HBSS-3, DS5-4) are listed in the table (to the right).

#### S9 Determining the difficulty of CD spectrum predictions

To quantify the difficulty of predicting the CD spectrum of each reference protein j in our data sets (SP175 and TS8) from its structural model, we computed the mean error of CD spectrum predictions ($RMSD_j^{\text{mean}}$) for protein each $j$. As described in Section 3.6, $RMSD_j^{\text{mean}}$ $were$determined by calculating CD spectra using six different prediction methods and averaging the deviations between the measured CD spectrum and the predicted spectra at every wavelength. We also determined the normal range of $RMSD_j^{\text{mean}}$ by calculating the average ($RMSD_{\text{set}}^{\text{mean}}$) and standard deviation of theses values scatter ($\sigma_{\text{set}}$) for our reference sets. We found that these values were similar for the SP175 ($RMSD_{\text{fit}}^{\text{mean}}$ = 3.3 kMRE, $\sigma_{\text{fit}}$= 1.1 kMRE) and TS8 data sets ($RMSD_{\text{cross}}^{\text{mean}}$ = 3.2 kMRE, $\sigma_{\text{cross}}$=1.3 kMRE). As shown in Fig. S3A, we considered proteins as outliers, and therefore, particularly difficult to predict, if their $RMSD_j^{\text{mean}}$ value was larger than $RMSD_{\text{cross}}^{\text{mean}} + \sigma_{\text{cross}}^{\text{mean}}$ = 4.5 kMRE.

Next, we tested if the large $RMSD_j^{\text{mean}}$ values of the outliers – listed in Fig. S3B – can be explained based on two potential sources of error, namely inaccuracy of the used structural models, and the inaccurate normalization of the measured spectrum intensities (identified in Sections 5.1 and 5.2, respectively).

First we accounted for the inaccurate normalization of the reference CD spectra by rescaling the intensities to match the intensity of the predictions (see Section 5.2). Scaling the measured spectra decreased the mean RMSD of the outlier proteins (henceforth the OP13 set) significantly from 6.2 kMRE to 3.6 kMRE), but also decreased $RMSD_{\text{cross}}^{\text{mean}}$ to 2.7 kMRE, and $\sigma_{\text{cross}}$ to 0.7 kMRE, thus lowering the threshold of what is considered an outlier to 3.4 kMRE. Despite the lower RMSD threshold, intensity scaling reduced $RMSD_j^{\text{mean}}$ for five of the thirteen proteins (colored blue in Fig. S3) by so much that they were no longer considered as outliers, including the reference protein with the largest RMSD; Subtilisin Carlsberg (SP175/67, spectra shown in Fig. S3C). Then, the rescaled CD spectra were subjected to deconvolution using the four SESCA basis sets (Section 3.6) to estimate and account for errors in the SS composition ($\Delta SS_j^{\mathrm{mean}}$) of the reference structures. Calculating the CD spectra from the ideal SS composition further reduced both $RMSD_{\mathrm{OP13}}^{\mathrm{mean}}$ to 2.1 kMRE and the threshold RMSD to 2.0 kMRE. In addition, it indicated that for two outliers (in magenta) − Avidin (SP175/8) and Glutamate dehydrogenase (SP175/34) − the above average $RMSD_j^{\mathrm{mean}}$ is due to the combination of inaccurate spectrum normalization and model inaccuracies.

For the remaining six outlier proteins (in red) $RMSD_j^{\mathrm{mean}}$ was reduced but remained above the normal range even after deconvolution, thus intensity scaling and model inaccuracies are not sufficient to explain the poor agreement between the measured and predicted spectra. To further investigate the reasons behind the poor prediction accuracy of the remaining six outliers, we included contributions from the AA side chains, by calculating the CD spectra using four mixed basis sets (DS-dTSC3, DSSP-1SC3, HBSS-3SC1, and DS5-4SC1) Side-chain corrections eliminated two more proteins − Jacalin (SP175/41) shown in Fig S3D and Carbonic anhydrase (SP175/14) − from the list of outliers, reducing $RMSD_j^{\mathrm{mean}}$ of the predicted spectra by ~1.0 kMRE in both cases.

Closer inspection of the reference structures also revealed some common features that allowed us to speculate on possible reasons behind their poor obtained RMSDs. Four out of six outlier structures include $Zn^{2+}$ or $Fe^{2+}$ ions coordinated to histidine, glutamate, and aspartate side chains. These coordination centers likely have their own contribution to the CD spectrum, which is not accounted for in our predictions, and increases $RMSD_j^{\mathrm{mean}}$ for these proteins.

Additionally, five out of six outlier proteins proteins – except for hemerythrin (TS8/1) – have an unusually high fraction of turn structures, which is represented in a very coarse-grained way in our smaller optimized basis sets. Indeed, when the spectra were deconvolved using the DS5-4SC1 basis set which has three basis spectra dedicated to these structures $RMSD_j$ for four of the six – except for hemerythrin and insulin (SP175/40) – proteins were in the normal range.

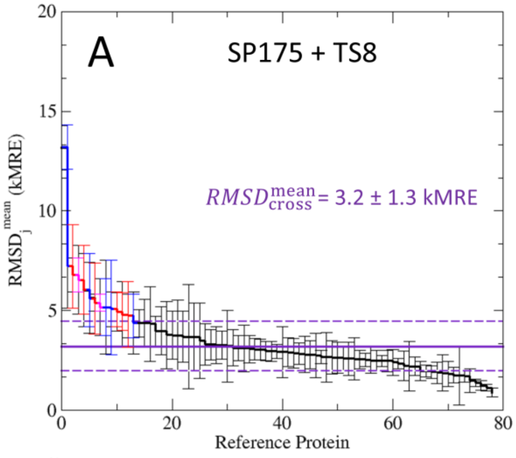


| B | $RMSD_j^{mean}$ (kMRE) | | | $SF_j^{mean}$ | $\Delta SS_j^{mean}$ |
|---|---|---|---|---|---|
| | no corr | scaled | deconv. | | (%) |
| SP175 / 67 | 13.17 | 1.44 | 0.915 | 0.4 | 7.7 |
| SP175 / 14 | 7.23 | 4.89 | 2.726 | 0.13 | 25.3 |
| SP175 / 34 | 6.78 | 4.03 | 1.69 | 0.71 | 27 |
| TS8 / 1 | 6.54 | 3.85 | 2.52 | 1.28 | 17.8 |
| SP175 / 7 | 6.03 | 3.62 | 1.19 | 0.54 | 8.6 |
| SP175 / 40 | 5.62 | 4.79 | 3.09 | 1.33 | 12.5 |
| SP175 / 8 | 5.38 | 3.58 | 1.89 | 0.46 | 18 |
| SP175 / 3 | 5.17 | 2.58 | 1.54 | 0.64 | 16.2 |
| SP175 / 56 | 5.16 | 2.59 | 1.45 | 1.47 | 7.4 |
| SP175 / 16 | 5.07 | 4.52 | 2.77 | 1.31 | 14.7 |
| SP175 / 41 | 4.95 | 3.81 | 3.01 | 0.4 | 17.9 |
| SP175 / 49 | 4.79 | 4.27 | 2.68 | 1.27 | 18.7 |
| SP175 / 32 | 4.73 | 3.42 | 1.30 | 0.89 | 20.8 |
| outlier above | 4.46 | 3.42 | 2.03 | | |

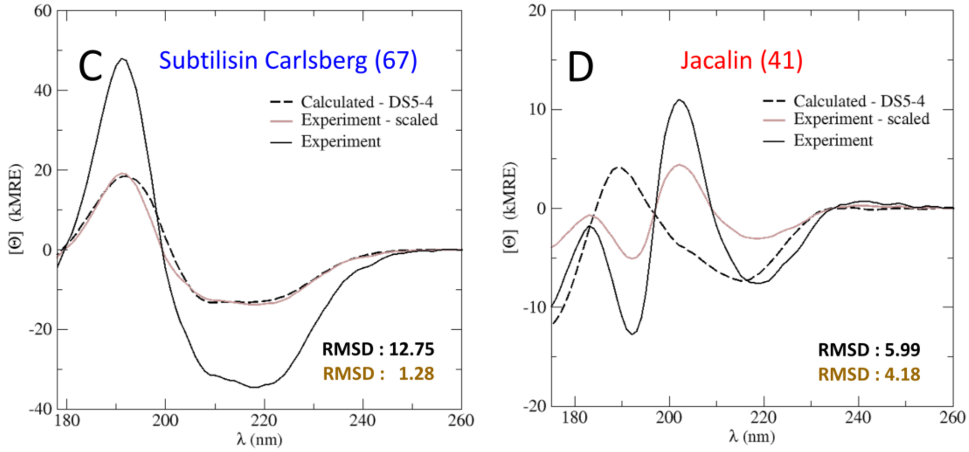


**Figure S3:** Outliers of the SP175 and TS8 reference data sets. A) Reference proteins sorted by the mean deviation ($RMSD_j^{mean}$) between their measured reference CD spectrum and spectra calculated from their reference structure. . Proteins were selected as outliers if their $RMSD_j^{mean}$

are by more than one standard deivation (dashed purple lines) larger than the mean prediction accuracy ($RMSD_{\mathrm{cross}}^{\mathrm{mean}}$, purple solid line) of the TS8 set. To determine the problem with the outliers their measured spectrum was scaled to match the predicted spectra, then deconvolved to find the SS composition that fits the spectrum best. B) List of the mean RMSDs for the thirteen outliers in the two data sets, including their set and ID number, $RMSD_j^{\mathrm{mean}}$ before corrections (no corr), after scaling (scaled), and subsequent deconvolution (deconv), along with the used scaling factor ($SF_j^{\mathrm{mean}}$), and the calculated SS error of the structural model ($\Delta SS_j^{\mathrm{mean}}$). Outlier proteins in the figure are colored blue if their large RMSD is mostly due to incorrect intensity scaling, magenta if it is due to scaling combined with model inaccuracies, and red otherwise. C) Example protein 1: significant RMSD improvement by scaling the experimental CD spectrum and D) Example protein 2: where scaling could not improve the RMSD significantly. For panels C and D the experimental CD spectrum is shown as a solid black line, the rescaled experimental spectrum is shown as a solid brown line, and the spectrum calculated by the SESCA basis set DS5-4 is shown as a black dashed line. The name and ID number of the protein is shown on the top of the panel, while the RMSDs for the predicted spectra in kMRE units are shown on the bottom in black and brown respectively.

#### S10 Amino acid side-chain contributions in the far UV range

We derived the average contribution of side-chain groups to the CD signal of proteins as described in Section S3 from a new mixed reference set (MP79), which included 59 globular proteins of the SP175 reference set and a set of 20 tri-peptides designated as GXG20 set (where ‘X’ stands for one of the twenty natural AAs). As shown in Fig. S4A, the CD spectra of the GXG20 peptide set differ substantially from one another, despite the fact that the peptides were too short to form the hydrogen bonds required for stable α-helices and β-sheets, and therefore mostly adopted a random coil structure. We therefore assumed that the spectra of these peptides are largely defined by their single side-chain group. The GXG20 spectra already indicate that aromatic side groups – and particularly phenyl-alanine and tyrosine – have strong positive contributions to the CD spectra, which differs from the signals of other side chains. The CD spectrum of the GAG peptide, on the other hand, shows the largest negative peak at ~195 nm, similar to CD signal that is associated with a random coil protein, whereas the CD signal of the GGG peptide – in the absence of a chirality center – is very weak.

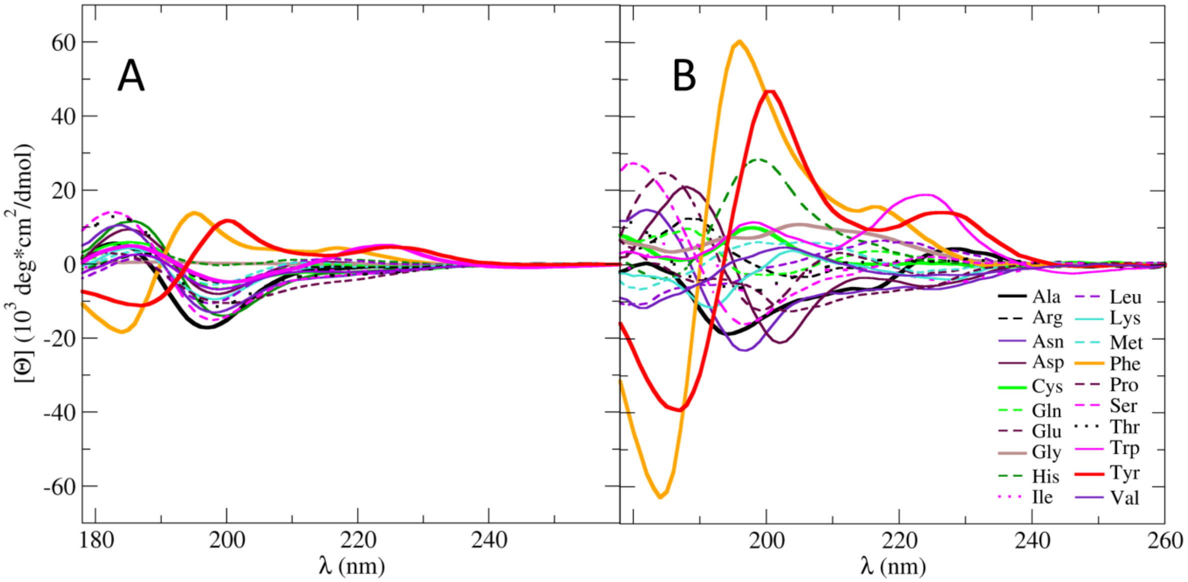


**Figure S4:** Circular dichroism contribution of AA side chains. A) Experimentally measured CD spectra measured at neutra p*H* for the 20 Ac-GXG-$NH_2$ peptides (GXG20). B) Calculated side-chain contributions for each AA side chain, derived from the SS composition and CD spectra of 59 globular proteins and the GXG20 peptides. The (CD/basis) spectra are color coded according to the AA side-chain groups they represent.

The pure basis spectra we derived from MP79 reference set (Fig S4B) are significantly larger than the CD spectra of the independent AAs (due to the normalization for the number of residues), and confirm the large contributions of the phenyl-alanine and tyrosine side chains. In addition, the basis spectra show moderate contributions from the AA side groups of asparagine, aspartate, glutamate, histidine, serine, and tryptophan, while the side chains of other AAs such as valine, isoleucine, and threonine had weaker CD signals.

We note that the GXG20 CD spectra differed considerably from the spectra of isolated amino acids measured by Nisihno *et al* [6]. at neutral, acidic and basic *pH*. In Fig. S32 we compare the two sets of CD spectra at neutral *pH* (7.0), where – despite the differences – the strongest signals were observed for phenyl-alanine, tyrosine, tryptophan.

#### S11 Analysis of the PCA basis sets

We determined the best achievable accuracy for a given number of basis spectra from basis sets that were constructed from the eigenvectors of a PCA of the SP175 reference CD spectra as described in Section S5. The first ten obtained PCA basis spectra are illustrated in Fig. S5A. In line with previous results [7–9], the first two PCA basis spectra are similar to the CD spectrum of purely α-helical and β-sheet proteins, and represent already about 94% of the variance within the spectra of the reference data set. As the sorted eigenvalues (Fig. S5B) suggest, only a few basis spectra should be required to achieve good to very high accuracy. Indeed, almost 99% of the variance of the SP175 CD spectra are represented by only the first five basis spectra, and the first ten basis spectra essentially describe the full data set. This expectation is confirmed in Fig. S5C, showing the reconstruction of the α-amylase precursor spectrum (SP175/3) using one to ten PCA basis spectra. For this spectrum already the first three basis spectra allow a good reconstruction with an average RMSD of 2.105 kMRE units ($10^3$ deg*cm$^2$/dmol), and using more than six or seven basis spectra essentially recovers the reference spectrum. For comparison, the average spectrum is shown in brown, corresponding to using no basis spectra at all, and serving as a lower limit of how well the spectra can be 'predicted' without any information. The table in Fig. S5D quantifies the changes in fitting accuracy for three sample spectra, taken from representative proteins of the three main structure classes (α-helical, β-sheet, and mixed α/β) and also provides the average RMSD for all 71 spectrum reconstructions (RMSD_ref).

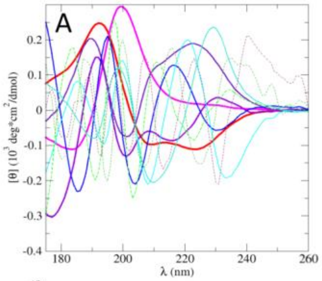


B

| PC # | Eigenvalue | Var % | Cumulative |
|---|---|---|---|
| 1 | 4417 | 81.0 | 81.0 |
| 2 | 677.6 | 12.4 | 93.4 |
| 3 | 138.8 | 2.55 | 96.0 |
| 4 | 103.3 | 1.89 | 97.9 |
| 5 | 47.7 | 0.87 | 98.8 |
| 6 | 42.8 | 0.79 | 99.6 |
| 7 | 11.4 | 0.21 | 99.8 |
| 8 | 8.4 | 0.15 | 99.9 |
| 9 | 1.6 | 0.029 | 99.9 |
| 10 | 0.8 | 0.015 | 100.0 |

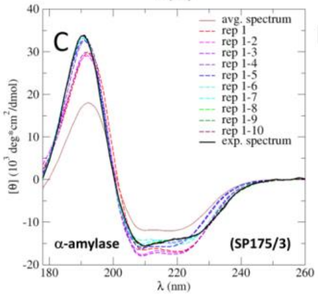


D

| Basis set # | Aamy $R_3$ | Cah1 $R_{14}$ | Cona $R_{20}$ | SP175 av. $RMSD_{ref}$ |
|---|---|---|---|---|
| 0 | 5.455 | 12.534 | 6.131 | 6.359 |
| 1 | 2.459 | 3.170 | 4.236 | 2.820 |
| 2 | 2.180 | 2.259 | 1.083 | 1.680 |
| 3 | 2.105 | 2.245 | 1.044 | 1.335 |
| 4 | 1.359 | 1.179 | 1.023 | 0.955 |
| 5 | 1.121 | 0.705 | 1.018 | 0.720 |
| 6 | 0.682 | 0.221 | 0.458 | 0.481 |
| 7 | 0.466 | 0.212 | 0.350 | 0.350 |
| 8 | 0.207 | 0.207 | 0.243 | 0.237 |
| 9 | 0.201 | 0.191 | 0.194 | 0.203 |
| 10 | 0.157 | 0.156 | 0.176 | 0.182 |

**Figure S5:** Principal component analysis of the SP175 protein CD spectra. A) graphical representation of the first 10 principal component vectors sorted by their contribution to the spectral variance. B) Eigenvalue, contribution to variance, and cumulative contribution to the spectral variance for the same PC vectors. C) Reconstruction of the CD spectrum of α-amylase (Aamy) by its projection on the first 0-10 PC vectors. The original spectrum is shown in black, the average spectrum of SP 175 data set is shown in brown. The reconstructed spectra are shown as colored dashed lines. D) RMSD between the reconstruction of three selected proteins – α-amylase, carbonic anhydrase I (Cah1), and Concanavalin A (Cona) – and their original CD spectrum as function of PC vectors used. The column SP175 av. shows average RMSD for all 71 proteins in the data set.

To determine how much the SS elements and side chains contribute to the variance of CD spectra in the SP175 reference set, we also analyzed the correlations between the principal components describing the shape of the CD spectra (see Section S5) and the occurrence of AAs and SS elements in the reference proteins. To this aim, we calculated the Pearson correlation coefficients between the projections of the first ten PC vectors, the AA composition of the proteins, as well as the SS compositions determined by the BestSel, DISICL, DSSP and HBSS algorithms.

Table S11 shows those structural properties which correlate most strongly with the principal components (PCs) of the CD spectra. As can be seen, the first three principal components involve mainly SS elements: PC 1 − which accounts for over 80% of the spectral variance of the reference set − was very strongly correlated ($R_{pearson}$ ~0.9) to the presence of α-helices in the protein structure, whilst PC 2 and 3 are moderately correlated to β-strand and turn structures. However, PCs 4, 6, 9, and 10 correlate more strongly with the presence of AAs than SS elements. Since these principal components describe ~3% of the spectral variance, one would expect a somewhat smaller but still notable contribution from side-chain groups. In addition, the most commonly considered correction to CD spectra are associated with the aromatic side chains of tryptophan, phenyl-alanine, and tyrosine because these AAs have the strongest CD signals in isolation. Our analysis also suggests that AA side chains with weaker CD activity, particularly arginine, histidine, cysteine and serine, may also contribute significantly to the CD spectra.

#### S12 Model validation for the P53/CBP complex

The quality of the structure models was determined through cross-validation by predicting the NMR chemical shifts from both structural models (NMR and MD) and then comparing the results to the experimental chemical shifts from the original NMR measurements (obtained from biological magnetic resonance databank, entry no. 17073). We computed the backbone chemical shifts (including those for the N, $C_\alpha$, $C_\beta$, C, $H_N$, and $H_\alpha$ atoms) using the chemical shift predictor Sparta+ [10]. Figure S6A shows the comparison between the experimental and calculated $C_\alpha$ secondary chemical shifts. Secondary chemical shift values are corrected for the average random coil chemical shift of the AA, and therefore indicative of the local protein (secondary) structure. A sequence of large positive secondary $C_\alpha$ shifts indicates a high propensity for α-helix in that region, whilst a sequence of large negative values shows a preference towards β-strands or extended structures. The overall agreement between the measured and predicted chemical shifts was quantified the through average RMSD of their secondary chemical shift profiles.

The comparison in Fig. S6A also revealed that the RMSD of the MD ensemble $C_\alpha$ chemical shifts (1.06 ppm) was lower than that of the NMR bundle (1.39 ppm). The same trends were observed for the average RMSD of all backbone chemical shifts as well. The $C_\alpha$ chemical shifts also indicate that our models agree well with the experimental chemical shifts on the position of the helical regions, but overestimate the helix propensities, especially for the C-terminal helix of CBP-NCBD, and the helical regions in P53-AD2. These regions are also the ones where the average SS composition is considerably less helical in the MD ensemble than in the NMR bundle. Additionally, the residues of the short β-sheets observed only in the MD model possess some of the largest negative $C_\alpha$ secondary chemical shifts of the experimental profile, suggesting that presence of these β-sheets also contribute to the lower average RMSD of the MD model.

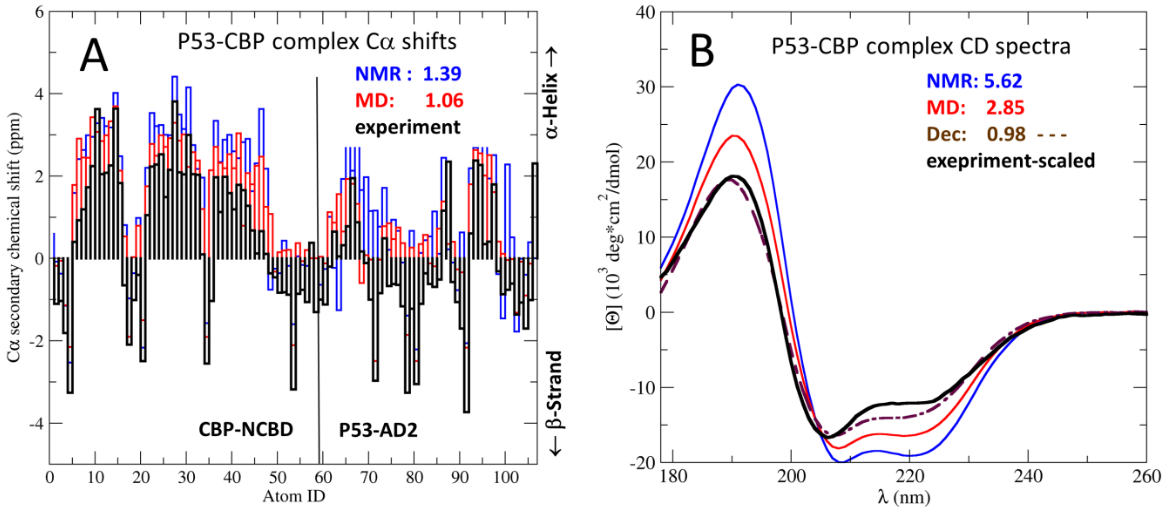


**Figure S6:** Model validation by comparison to experimental observables. The measured observables (experiment) were compared to the same observables calculated from two structural models (NMR and MD). Panels of the figure show the comparison of A) the Cα secondary chemical shifts and B) the CD spectra, respectively. The measured observables are shown as black solid lines, calculated observables are shown in different colors according to the underlying model. The dashed spectrum in B) shows the prediction from the ideal SS composition determined by spectrum deconvolution (Dec). The average RMSD from the experimental observable (in ppm and mean residue ellipticity units, respectively) for each model is shown in the corresponding color. The CD spectra in B) were calculated using the DS-dTSC3 basis set, and include side-chain corrections.

We also analyzed the differences in the SS composition of the two models. A summary over SS composition of each structural model is shown below their cartoon representation in Fig. 8A. As the figure shows, about 47% of the residues in the NMR are in an $\alpha$-helical conformation. Although no $\beta$-sheets appeared in this model, a low percentage AAs adopted a local conformation typical for an extended $\beta$-strand at the termini of the two protein domains.

The P53/CBP complex was very dynamic during the MD simulations. The two domains remained strongly bound during the simulation, but the conformational fluctuations resulted in a 38% average $\alpha$-helix content. In addition, while the total $\beta$-strand content decreased slightly in the MD model compared to the NMR bundle, 2.8% of the residues in the MD model was in a regular $\beta$-strand conformation, and established the hydrogen bonds to form two short $\beta$-sheets which appeared with ~15% probability in the ensemble. These short $\beta$ sheets connected the N-terminus of CBP-NCBD with residues 25-27 of P53-AD2, and the two termini P53-AD2. Predicting CD spectra from these SS compositions (Fig. S6B) suggested similar trends observed for the $C_\alpha$ secondary chemical shifts, whereas deconvolution of the measured CD spectrum indicated an ideal SS composition that is 8-10% less helical than that of the MD model.

### Additional Supplementary Tables:

**Table S1:** The SP175 dataset. Identification number (ID), abbreviation (short code), protein databank (pdb) code, protein circular dichroism databank (pcddb) code, and name for each of the proteins in the dataset is provided, as well as number residues (size) and resolution (resol.) of the model structure. For further details see ref. [11]. Entries marked by plus signs in the ID sections (and highled in red) were part of the SP175 dataset, but not part of the TR64 dataset, whilst entries marked by stars ( and highlighted in blue) were part SP175 but not part of the GP59 dataset

| ID number | short code abbrev. | structure pdb code | CD spectrum pcddb code | Protein Name | Size (res) | resol. (A) |
|---|---|---|---|---|---|---|
| 1 | Aldo | 1ado | CD0000001 | Aldolase | 1452 | 1.90 |
| 2 | alkp | 1ed9 | CD0000002 | Alkalnie phosphatase | 898 | 1.75 |
| 3* | aamy | 1vjs | CD0000003 | Alpha amylase prec | 469 | 1.70 |
| 4 | abun | 1hc9 | CD0000004 | Alpha-bungarotoxin | 173 | 1.80 |
| 5 | actp | 5cha | CD0000005 | Alpha Chymotrypsin | 474 | 1.67 |
| 6 | actn | 2cga | CD0000006 | Alpha chymotrypsinogen | 490 | 1.80 |
| 7* | ptib | 5pti | CD0000007 | Pancreatic trypsin inhibitor (bov.) | 58 | 1.00 |
| 8* | avdn | 1rav | CD0000008 | Avidin (recombinant) | 248 | 2.20 |
| 9 | bamy | 1fa2 | CD0000009 | Beta amylase (sweet potato) | 498 | 2.30 |
| 10 | bglc | 1bgl | CD0000010 | Beta galactosidase | 8168 | 2.50 |
| 11 | blac | 1b8e | CD0000011 | Beta lactoglobulin (bovine) | 152 | 1.95 |
| 12 | cphy | 1ha7 | CD0000012 | C-phycocyanin | 3996 | 2.20 |
| 13 | calm | 1lin | CD0000013 | Calmodulin | 146 | 2.00 |
| 14* | cah1 | 1hcb | CD0000014 | Carbonic anhydrase I | 258 | 1.60 |
| 15 | cah2 | 1v9e | CD0000015 | Carbonic anhydrase II (bovine) | 518 | 1.95 |
| 16* | carp | 5cpa | CD0000016 | Carboxypeptidase | 307 | 1.54 |
| 17 | ecat | 1dgf | CD0000017 | Erythrocyte catalase (human) | 1998 | 1.50 |
| 18 | cerp | 1kcw | CD0000018 | Ceruloplasmin (human) | 1017 | 3.00 |
| 19 | cits | 2cts | CD0000019 | Citrate synthase | 437 | 2.00 |
| 20+ | cona | 1nls | CD0000020 | Concanavalin A | 237 | 0.94 |
| 21+ | cytc | 1hrc | CD0000021 | Cytochrome C (horrse heart) | 104 | 1.90 |
| 22 | bbcr | 2bb2 | CD0000022 | Beta b2 crystallin (bovine) | 176 | 2.10 |
| 23 | gbcr | 4gcr | CD0000023 | Gamma b crystallin (bovine) | 174 | 1.47 |
| 24 | gdcr | 1elp | CD0000024 | Gamma d crystalline | 346 | 1.95 |
| 25 | gecr | 1m8u | CD0000025 | Gamma e crystallin (bovine) | 173 | 1.65 |
| 26 | gscr | 1ha4 | CD0000026 | Gamma S crystallin C-term (human) | 174 | 2.40 |
| 27 | gdc2 | 1hk0 | CD0000027 | Gamma d crystallin (human) | 173 | 1.25 |
| 28 | dqd1 | 1qfe | CD0000028 | Dehydroquinate dehydratase I (S. Thypi) | 504 | 2.10 |
| 29 | dqd2 | 2dhq | CD0000029 | Dehydroquinate dehydratase I (M. Tuber) | 136 | 2.00 |
| 30 | dna1 | 3dni | CD0000030 | Dnase I | 258 | 2.00 |
| 31 | pela | 3est | CD0000031 | Pancreatic elastase | 229 | 1.65 |
| 32* | fesf | 2fdn | CD0000032 | 2[4Fe-4S] ferredoxin | 55 | 0.94 |
| 33 | glco | 1cf3 | CD0000033 | Glucose oxidase | 581 | 1.90 |
| 34* | gldh | 3mw9 | CD0000034 | Glutamate dehydrogenase (bovine) | 3009 | 2.40 |

**Table S1:** The SP175 dataset (cont.)

| ID number | short code abbrev. | structure pdb code | CD spectrum pcddb code | Protein Name | Size (res) | resol. (A) |
|---|---|---|---|---|---|---|
| 35 | glps | 1gpb | CD0000035 | Glycogen phosphorylase | 823 | 1.90 |
| 36 | hadh | 1bn6 | CD0000036 | Haloalkaene dehalogenase | 291 | 1.50 |
| 37+ | hglb | 1hda | CD0000037 | Hemoglobin (human) | 572 | 2.20 |
| 38 | hsa1 | 1n5u | CD0000038 | Serum Albumin (human) | 583 | 1.90 |
| 39 | igg2 | 1igt | CD0000039 | Immunglobulin G2a (mouse) | 1300 | 2.80 |
| 40* | hins | 1trz | CD0000040 | Insulin A&C (human) | 102 | 1.60 |
| 41* | jaca | 1ku8 | CD0000041 | Jacalin | 596 | 1.75 |
| 42 | lacf | 1blf | CD0000042 | Lactoferrin (bovine) | 685 | 2.80 |
| 43 | lect | 1les | CD0000043 | Lectin (lentil) | 458 | 1.90 |
| 44 | lept | 1ax8 | CD0000044 | Leptin E100 (human) | 130 | 2.40 |
| 45+ | lysm | 193l | CD0000045 | Lysozime (hen, egg-white) | 129 | 1.33 |
| 46 | moll | 1mol | CD0000046 | Mollein | 188 | 1.70 |
| 47 | mmyo | 1ymb | CD0000047 | Metmyoglobin | 153 | 1.90 |
| 48+ | Mglb | 1a6m | CD0000048 | Myoglobin (oxy) | 151 | 1.00 |
| 49* | Nmra | 1k6j | CD0000049 | Nmra trans. Regulator | 634 | 1.80 |
| 50 | Oalb | 1ova | CD0000050 | Ovalbumin | 1446 | 1.95 |
| 51 | Otrn | 1dot | CD0000051 | Ovotransferrin (duck) | 686 | 2.35 |
| 52+ | Papn | 1ppn | CD0000052 | Papain (papaya) | 212 | 1.60 |
| 53 | lec2 | 1ofs | CD0000053 | Lectin (pea) | 460 | 1.80 |
| 54 | Plyc | 1air | CD0000054 | Pectate lyase C | 352 | 2.20 |
| 55 | Peps | 2psg | CD0000055 | Pepsinogen | 326 | 1.80 |
| 56* | pox1 | 7atj | CD0000056 | Peroxydase C1A (horse-raddish) | 305 | 1.47 |
| 57 | Pgmu | 3pmg | CD0000057 | Phosphoglucomuatse (rabbit) | 1120 | 2.40 |
| 58 | Pgkn | 3pgk | CD0000058 | Phosphoglycerate kinase | 415 | 2.50 |
| 59 | pla2 | 1une | CD0000059 | Phospholipase A2 (bovine) | 123 | 1.50 |
| 60 | Pemt | 1hnn | CD0000060 | Phenylethanolamine N-methyltransferase | 528 | 2.40 |
| 61 | Pkin | 1a49 | CD0000061 | Pyruvate kinase | 4152 | 2.10 |
| 62 | Rhod | 1rhs | CD0000062 | rhodanase (sulphur-subst, bovine) | 292 | 1.36 |
| 63+ | Rnas | 3rn3 | CD0000063 | Ribonuclase A (bovine) | 124 | 1.45 |
| 64 | Rubr | 1r0i | CD0000064 | Rubredoxin (Cadmium subs) | 53 | 1.50 |
| 65 | tpi2 | 1ba7 | CD0000065 | Trypsin inhibitor (soy bean) | 334 | 2.50 |
| 66 | Stvn | 1stp | CD0000066 | Streptavidin | 121 | 2.60 |
| 67* | subc | 1scd | CD0000067 | Subtilisin Carlsberg (crosslinked) | 247 | 2.30 |
| 68 | cttp | 1cgj | CD0000068 | Chmyotrypsinogen A + Trypsin inhibitor | 301 | 2.30 |
| 69 | thau | 1thw | CD0000069 | Thaumatin | 207 | 1.75 |
| 70 | tpis | 7tim | CD0000070 | Triosephosphate isomerase | 494 | 1.90 |
| 71 | ubqn | 1ubi | CD0000071 | ubiquitin (human) | 76 | 1.80 |

**Table S2**: The TS8 globular protein cross -validation protein set Identification number (ID), abbreviation (short code), RSCB protein databank (pdb) code, circular dichroism filename, and name for each of the proteins in the dataset is provided, as well as number residues (size) and resolution (resol.) of the model structure. The CD spectra of this dataset were extracted from the globular protein training set provided with the CCA deconvolution program of Hollósi et al. See ref.[2] for further details.

| ID number | short code abbreviation | structure pdb code | CD spectrum short code | Protein Name | Size (res) (res) | resol. (A) |
|---|---|---|---|---|---|---|
| 1 | hmrt | 4xpx | CD_HMRT.dat | Hemerythrin | 130 | 1.03 |
| 2 | azu | 5azu | CD_AZU.dat | Azurin | 512 | 1.90 |
| 3 | pral | 2pab | CD_PRAL.dat | Prealbumin | 228 | 1.80 |
| 4 | ldh | 6ldh | CD_LDH.dat | Lactate dehydrogenase | 329 | 2.00 |
| 5 | thml | 8tln | CD_THML.dat | Thermolysin | 318 | 1.60 |
| 6 | gdp | 3gdp | CD_GDP.dat | Hydroxynitrile lyase | 1042 | 1.57 |
| 7 | subn | 1sbt | CD_SUBN.dat | Subtilisin novo | 275 | 2.50 |
| 8 | tnf | 2tnf | CD_TNF.dat | Tumor necrosis factor alpha | 444 | 1.40 |

**Table S3:** The EV9 evaluation protein dataset. Identification number (ID), abbreviation (short code), RSCB protein databank (pdb) code, circular dichroism filename, and name for each of the proteins in the dataset is provided, as well as number residues (size) and resolution (resol.) of the model structure. The CD spectra of this dataset were extracted from the globular protein training set provided with the CCA deconvolution program of Hollósi et al. See ref. [2] for further details.

| ID number | short code abbreviation | structure pdb code | CD spectrum short code | Protein name | Size (res) (res) | resol. (A) |
|---|---|---|---|---|---|---|
| 1 | mglb | 1vxf | CD_MGLB.out | Myoglobin | 153 | 1.70 |
| 2 | hglb | 2mhb | CD_HGLB.out | Hemoglobin | 287 | 2.00 |
| 3 | cytc | 5cyt | CD_CYTC.out | Cytochrome C | 103 | 1.50 |
| 4 | lysm | 4lzt | CD_LYSM.out | Lysozyme | 129 | 0.95 |
| 5 | papn | 9pap | CD_PAPN.out | Papain | 211 | 1.65 |
| 6 | rnas | 1rbx | CD_RNAS.out | Ribonuclease A | 124 | 1.69 |
| 7 | bnjn | 1rei | CD_BNJN.out | Benes-Jones peptide | 214 | 2.00 |
| 8 | cona | 1nls | CD_CONA.out | Concanvallin A | 237 | 0.94 |
| 9 | suds | 1sxn | CD_SUDS.out | Superoxide dismutase | 302 | 1.90 |

**Table S4:** Secondary structure elements recognized by the structure classification algorithm DSSP (see ref [12]). DSSP recognizes SS elements based on backbone hydrogen bond patterns. Note that the 4-Helix class represents the regular $\alpha$-helix, while 3-Helix and 5-Helix is associated with the distorted $3_{10}$- and $\pi$-helices, respectively.

| **DSSP secondary structure classes** | | |
|---|---|---|
| **Struct. Element** | **K** | **Abbr.** |
| 4-Helix | 1 | 4H |
| Beta-Strand | 2 | BS |
| 3-Helix | 3 | 3H |
| 5-Helix | 4 | 5H |
| Beta-Bridge | 5 | BB |
| Bend | 6 | BE |
| Turn | 7 | TU |
| Unclassified | 8 | UC |

**Table S5:** Secondary structure elements recognized by the structure classification algorithm DISICL (see ref [13]). DISICL recognizes SS elements based on backbone dihedral angles in short segments of the protein, to provide a better resolution on loop- and turn structures.

| **Detailed DISICL classes** | | | **Simplified DISICL classes** | | |
|---|---|---|---|---|---|
| **Struct. Element** | **K** | **Abbr.** | **Struct. Element** | **abbr.** | **i** |
| α-Helix | 1 | ALH | Helical | HEL | 1 |
| π-Helix | 2 | PIH | Helical | HEL | 1 |
| Helix-Cap | 3 | HC | Helical | HEL | 1 |
| extended β-Strand | 4 | EBS | β-Strand | BS | 2 |
| normal β-Strand | 5 | NBS | β-Strand | BS | 2 |
| β-Cap | 6 | BC | β-Strand | BS | 2 |
| Turn type 2 | 7 | TII | β-Turn | BT | 3 |
| Turn type 8 | 8 | TVIII | β-Turn | BT | 3 |
| Turn type 1 | 9 | TI | 3-Helical Turn | 3HT | 4 |
| $3_{10}$ Helix | 10 | 3H | 3-Helical Turn | 3HT | 4 |
| Turn-Cap | 11 | TC | 3-Helical Turn | 3HT | 4 |
| Poly-proline Helical | 12 | PP | Irregular b | IRB | 5 |
| Bulge | 13 | Bu | Irregular b | IRB | 5 |
| Left-handed Helix | 14 | LHH | Left-Handed Turns | LHT | 6 |
| Left-handed Turn II | 15 | LTII | Left-Handed Turns | LHT | 6 |
| Haripin 2:2 | 16 | HP | Other Tight Turns | OTT | 7 |
| γ-Turn | 17 | GXT | Other Tight Turns | OTT | 7 |
| Schellmann-Turn | 18 | SCH | Other Tight Turns | OTT | 7 |
| Unclassified | 19 | UC | Unclassified | UC | 8 |

**Table S6:** Secondary structure elements recognized by the structure classification algorithm HbSS. This algorithm recognizes SS elements based on backbone hydrogen bond patterns, similarly to DSSP. In addition, it can identify parallel and anti-parallel β-strands (see Fig. S2), and further distinguish strand handedness (based on β-strand twist angles) to be more comparable to BestSel algorithm (see ref [3]).

| **Extended HbSS classes** | | | **HbSS secondary structure classes** | | |
|---|---|---|---|---|---|
| **Struct. Element** | **K** | **Abbr.** | **Struct. Element** | **K** | **Abbr.** |
| α-Helix | 1 | 4H | α-Helix | 1 | 4H |
| 3/10Helix | 2 | 3H | 3/10Helix | 2 | 3H |
| π-Helix | 3 | 5H | π-Helix | 3 | 5H |
| Left-twist. parallel β | 4 | LHP | Parallel β-strand | 4 | BSP |
| Non-twist. parallel β | 5 | NBP | Parallel β-strand | 4 | BSP |
| Right-twist. parallel β | 6 | RHP | Parallel β-strand | 4 | BSP |
| Left-twist. anti-par. β | 7 | LHA | Anti-par. β-strand | 5 | BSA |
| Non-twist. anti-par. β | 8 | NBA | Anti-par. β-strand | 5 | BSA |
| Right-twist. anti-par. β | 9 | RHA | Anti-par. β-strand | 5 | BSA |
| H-bonded Turn | 10 | TU | H-bonded Turn | 6 | TU |
| Unclassified | 11 | UNC | Unclassified | 7 | UNC |

**Table S7:** Secondary structure elements recognized by the CD deconvolution algorithm BestSel (see ref [3]). This deconvolution basis set was derived from the SP175 set, to provide a better resolution on β-strand SS elements. Note that the 'Irregular Helix' class is associated with the distorted ends of a regular α-helix, and the classical $3_{10}$-helix was assigned to the 'Other' class

| **BestSel deconvolution basis spectra** | | |
|---|---|---|
| **Struct. Element** | **k** | **Abbr.** |
| α-Helix | 1 | Helix1 |
| Irregular Helix | 2 | Helix2 |
| Left-twist anti-par. β | 3 | LHA |
| Relaxed anti-par. β | 4 | NBA |
| Right-twist anti-par. β | 5 | RHA |
| Parallel β-strand | 6 | NBP |
| Turns | 7 | Turn |
| Other | 8 | Other |

**Table S8:** Calculated accuracy of top-ranking optimized basis sets based on protein SS composition. The table shows the name of the basis set, the number of basis spectra (size), the average root mean square deviation (RMSD) between the experimental and calculated CD spectra for four protein reference sets (SP175, TR64, EV9, TS8). Basis sets in the table were derived using SP175 set and cross-validated on the TS8 set. The type section describes the underlying SS classification protocol used to derive the basis set (DS_sim, DS_det, Dssp, HBSS, or HBSS_ext, see Section 3.4 for details).

| Basis spectrum set | | | $RMSD_{set}$ (kMRE) | | | |
|---|---|---|---|---|---|---|
| **Name** | **Type** | **size** | **SP175** | **TR64** | **EV9** | **TS8** |
| DS3-1 | DS_sim | 5 | 3.603 | 3.718 | 2.998 | 3.124 |
| DS3-3 | DS_sim | 4 | 3.625 | 3.714 | 3.262 | 3.241 |
| DS5-4 | DS_det | 6 | 3.462 | 3.56 | 2.582 | 3.189 |
| DS6-1* | DS_det | 6 | 3.269 | 3.344 | 3.402 | 3.042 |
| DS-dT | DS_det | 3 | 3.696 | 3.793 | 3.131 | 3.091 |
| DSSP-1 | Dssp | 4 | 3.784 | 3.893 | 3.175 | 3.034 |
| DSSP-T* | Dssp | 3 | 3.819 | 3.924 | 3.274 | 3.012 |
| HBSS-2 | HBSS_ext | 6 | 3.705 | 3.752 | 3.559 | 3.536 |
| HBSS-3* | HBSS_ext | 5 | 3.754 | 3.794 | 3.594 | 3.288 |
| HBSS-4 | HBSS | 4 | 3.913 | 4.008 | 3.432 | 3.483 |
| DS_simF | DS_sim[I] | 8 | 3.465 | 3.576 | 3.19 | 3.326 |
| DS_detF | DS_det[I] | 19 | 2.777 | 2.812 | 3.896 | 3.909 |
| DSSP-F | Dssp[I] | 8 | 3.551 | 3.605 | 3.402 | 3.033 |
| HBSS-E | HBSS_ext[I] | 11 | 3.379 | 3.456 | 3.416 | 4.314 |
| HBSS-F | HBSS[I] | 7 | 3.7 | 3.744 | 3.606 | 3.508 |
| Sreer-1 | DS_det[D] | 6 | 7.631 | 7.593 | 6.91 | 5.163 |
| Perczel-1 | DS_det[D] | 5 | 4.717 | 4.880 | 3.562 | 3.748 |
| Perczel-2 | Dssp[D] | 5 | 4.819 | 4.947 | 3.690 | 3.426 |

*: most predictive basis set found for a given classification algorithm.

**D**: basis spectra were determined by deconvolution methods previously, and not computed in this study.

**I**: non-optimized initial basis sets.

**Table S9:** Calculated accuracy for other CD spectrum prediction methods, and mixed basis sets. The table shows the name of the basis set or method, the number of number of backbone and side-chain basis spectra (size), the average root mean square deviation (RMSD) between the experimental and calculated CD spectra for four reference sets (MP79, GXG20, SP175, and TS8). ). Basis sets in the table were derived using MP79 set and cross-validated on the TS8 set. The type section describes the underlying structure classification protocol for basis sets (DS_det, DSSP, or HbSS_ext) complemented by SC for side-chain corrections. For other methods: Best denotes a basis set is based on SS predicted from the CD spectra by the BestSel algorithm, corresponds to the RMSD using perfect structural models. MatM stands for direct spectrum calculation with the matrix method, whereas SVM abbreviates an empirical spectrum calculation based on support vector machine techniques.

| Basis spectrum set | | | $RMSD_{set}$ (kMRE) | | | |
|---|---|---|---|---|---|---|
| **Name** | **Type** | **size** | **MP79** | **GXG20** | **SP175** | **TS8** |
| DichroCalc | MatM[C] | ----- | ----- | ----- | 6.059 | 6.124 |
| PDB2CD | SVM[C,S] | ----- | ----- | ----- | 2.395 | 4.725 |
| BestSel_der | Best[S] | 8+0 | 2.859 | 4.136 | 2.931 | 1.828 |
| DS-dTSC3 | DS_det+SC | 3+4 | 2.895 | 2.282 | 3.728 | 2.879 |
| DS5-4SC1 | DS_det+SC | 6+6 | 2.755 | 2.21 | 3.574 | 2.704 |
| DS6-1SC1 | DS_det+SC | 6+6 | 2.664 | 2.462 | 3.397 | 2.911 |
| DSSP-1SC3 | DSSP+SC | 4+6 | 3.084 | 2.682 | 3.849 | 2.825 |
| DSSP-TSC1 | DSSP+SC | 3+6 | 3.139 | 3.162 | 3.763 | 2.844 |
| HBSS-3SC1 | HbSS_ext+SC | 5+4 | 3.014 | 2.31 | 3.833 | 3.019 |
| HBSS-3SCF | HbSS_ext+SC | 5+20 | 2.686 | 1.181 | 3.789 | 3.18 |

[S]: method trained on the SP175 reference set.

[C]: competing method.

**Table S10:** The TS14 protein dataset. Identification number (ID), abbreviation (short code), RSCB protein databank (pdb) code, protein circular dichroism databank filename, and name for each of the proteins in the dataset is provided, as well as number residues (size) and resolution (resol.) of the model structure. The CD spectra of this dataset were used to repeat the cross-validation performed by Mavridis et al. See ref. [14] for further details.

| ID number | short code abbrev. | structure pdb code | CD spectrum short code | Protein Name | Size (res) | resol. (A) |
|---|---|---|---|---|---|---|
| 1 | a1at | 1qlp | CD0003891000 | Alpha-1-antitrypsin | 394 | 2.0 |
| 2 | atr3 | 1sr5 | CD0003889000 | Antithrombin-III | 689 | 3.1 |
| 3 | bcr2 | 1ytq | CD0003670000 | Beta-crystallin B2 | 181 | 1.7 |
| 4 | bcr3 | 1bd7 | CD0003672000 | Beta-crystallin B2 | 347 | 2.78 |
| 5 | Bcr4 | 1oki | CD0003668000 | Beta-crystallin B1 | 366 | 1.4 |
| 6 | bcry | 1a7h | CD0003669000 | Gamma-crystallin S | 172 | 2.56 |
| 7 | bmgl | 2yxf | CD0003894000 | Beta-2-microglobulin | 99 | 1.13 |
| 8 | cxcn | 2ccm | CD0004676000 | Calexcitin | 386 | 1.8 |
| 9 | ectn | 1ecz | CD0003896000 | Ecotin | 284 | 2.68 |
| 10 | hutp | 1q5u | CD0003897000 | human dUTPase | 390 | 2.0 |
| 11 | ipdh | 2y3z | CD0003898000 | 3-isopropylmalate dehydrogenase | 351 | 1.8 |
| 12 | magl | 4acq | CD0003893000 | Alpha-2-macroglobulin | 5132 | 4.3 |
| 13 | npta | 2zxe | CD0001180000 | Na/K transporting ATPase | 1296 | 2.4 |
| 14 | sctx | 4kyp | CD0004244000 | Beta-scorpion toxin | 290 | 1.7 |

**Table S11:** Correlation analysis of the spectral components.

| PC1 | Corr. | Prop | Desc. | PC6 | Corr. | Prop | Desc. |
|---|---|---|---|---|---|---|---|
| 1 | 0.921 | Hel1 (Best) | α-helix | 1 | 0.201 | SER (AA) | Amino A. |
| 2 | 0.906 | Hel1 (SEL) | α-helix | 2 | 0.163 | CYS (AA) | Amino A. |
| 3 | 0.9 | ALH (DISICL) | α-helix | 3 | 0.138 | RHA (HbSS) | Strand |
| 4 | 0.898 | Hel (DISICL) | α-helix | 4 | 0.157 | Hel2 (Best) | Helix |
| 5 | 0.892 | 4H (DSSP) | α-helix | 5 | 0.126 | Hel1 (SEL) | α-helix |
| 6 | 0.891 | 4H (HbSS) | α-helix | 6 | 0.116 | ALH (DISICL) | α-helix |
| **PC2** | **Corr.** | **Prop** | **Desc.** | **PC7** | **Corr.** | **Prop** | **Desc.** |
| 1 | 0.532 | EBS (DISICL) | β-strand | 1 | 0.285 | RHP (HbSS) | β-strand |
| 2 | 0.513 | Anti1 (BEST) | β-strand | 2 | 0.274 | BSP (HbSS) | β-strand |
| 3 | 0.444 | NBA (HbSS) | β-strand | 3 | 0.25 | Para (Best) | β-strand |
| 4 | 0.418 | Anti2 (Best) | β-strand | 4 | 0.23 | Turn (Sel) | Turn |
| 5 | 0.395 | BS (HbSS) | β-strand | 5 | 0.205 | Bend (DSSP) | Turn |
| 6 | 0.352 | HIS (AA) | Amino A. | 6 | 0.169 | GXT (DISICL) | Turn |
| **PC3** | **Corr.** | **Prop** | **Desc.** | **PC8** | **Corr.** | **Prop** | **Desc.** |
| 1 | 0.31 | BS (HbSS) | β-strand | 1 | 0.386 | 3H (DSSP) | Helix |
| 2 | 0.299 | SCH (DISICL) | Turn | 2 | 0.344 | 3H (HbSS) | Helix |
| 3 | 0.254 | NBS (DISICL) | β-strand | 3 | 0.3 | 5H (HbSS) | Helix |
| 4 | 0.23 | Bend (DSSP) | Turn | 4 | 0.273 | HC (DISICL) | Turn |
| 5 | 0.216 | NBA (HbSS) | β-strand | 5 | 0.253 | MET (AA) | Amino A. |
| 6 | 0.205 | THR (AA) | Amino A. | 6 | 0.139 | Other (Best) | Turn |
| **PC4** | **Corr.** | **Prop** | **Desc.** | **PC9** | **Corr.** | **Prop** | **Desc.** |
| 1 | 0.471 | ARG (AA) | Amino A. | 1 | 0.223 | ASP (AA) | Amino A. |
| 2 | 0.397 | LHH (DISICL) | Turn | 2 | 0.202 | 3H(HbSS) | Helix. |
| 3 | 0.306 | Anti2 (Best) | β-strand | 3 | 0.192 | GLU (AA) | Amino A |
| 4 | 0.293 | NBA (HbSS) | β-strand | 4 | 0.152 | ILE (AA) | Amino A. |
| 5 | 0.299 | SCH (DISICL) | Turn | 5 | 0.152 | 3H (DSSP) | Helix |
| 6 | 0.272 | LHT (DISICL) | Turn | 6 | 0.126 | PIH (DISICL) | Helix |
| **PC5** | **Corr.** | **Prop** | **Desc.** | **PC10** | **Corr.** | **Prop** | **Desc.** |
| 1 | 0.394 | 3HT( DISICL) | Helix | 1 | 0.214 | PHE (AA) | Amino A. |
| 2 | 0.376 | 3H (DISICL) | Helix | 2 | 0.15 | TRP (AA) | Amino A. |
| 3 | 0.33 | 3H (DSSP) | Helix | 3 | 0.14 | SER (AA) | Amino A. |
| 4 | 0.321 | 3H (HbSS) | Helix | 4 | 0.133 | RHA (HbSS) | β-strand |
| 5 | 0.296 | Cys (AA) | Amino A. | 5 | 0.116 | Bend (DSSP) | Turn |
| 6 | 0.294 | Hel2 (SEL) | Helix | 6 | 0.102 | LHT (DISICL) | Turn |

The six best correlated structural properties are listed for each of the first ten principal components of the SP175 CD spectra. The table displays the abbreviated code of the structural property (Prop), the Pearson correlation score (Corr.) between the projections of the PC vector, and the coefficients of the structural property (the fraction of SS element or AA in a protein), as well as the type and a short description of the structural property. The type (in parenthesis) defines the source algorithm for SS elements (DSSP, HbSS, DISICL or BestSel algorithms) and (AA) stands for amino acids. The short description shows if a SS element is either associated with α-helix, irregular helix (Helix), β-strand or turn structures.

#### Additional Supplementary Figures:

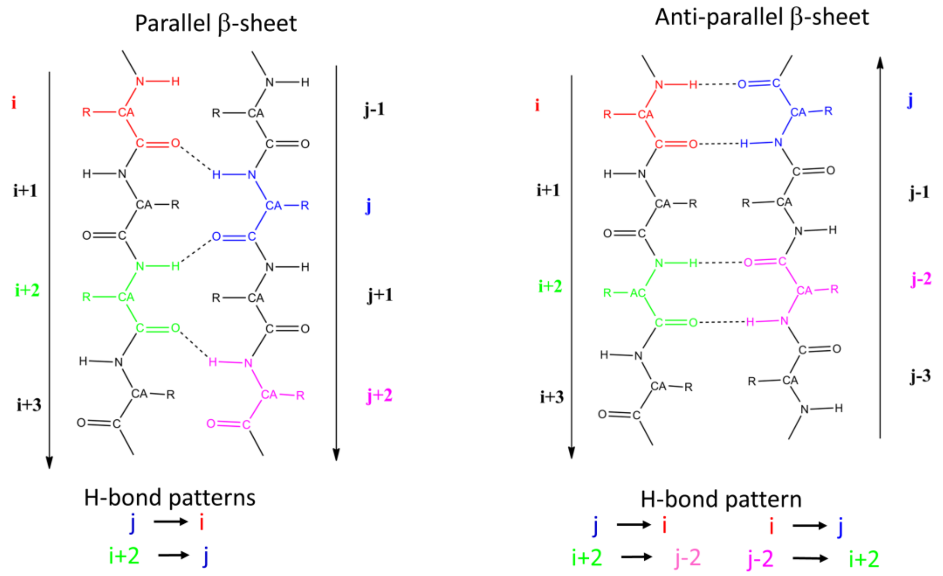


**Figure S7:** Definition of parallel and anti-parallel hydrogen bonding for the algorithm HbSS. The indicated AAs relative to the hydrogen bonded pair i and j are identified as a parallel or anti-parallel β-sheet, if hydrogen bonds indicated at the bottom are present. The arrows denote a hydrogen bond between N-H donor group on the left side and the C=O acceptor group on right side.

| Map_DSSP-T | i | 1 | 2 | 3 |
|---|---|---|---|---|
| k | $\alpha_{ki}$ | Helix | Sheet | Coil |
| 1 | 3H | 1 | 0 | 0 |
| 2 | 4H | 1 | 0 | 0 |
| 3 | 5H | 1 | 0 | 0 |
| 4 | BS | 0 | 1 | 0 |
| 5 | BB | 0 | 0 | 1 |
| 6 | TU | 0 | 0 | 1 |
| 7 | BE | 0 | 0 | 1 |
| 8 | UC | 0 | 0 | 1 |

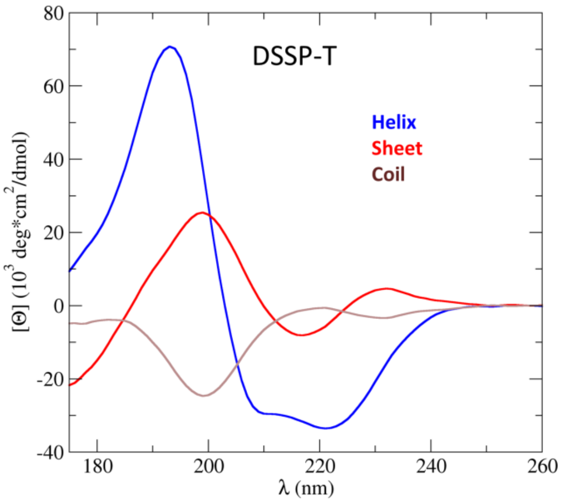


**Figure S8:** Assignment (left) and basis spectra (right) for the basis set DSSP-T. Secondary structure elements for the assignment are abbreviated according to Table S4.

| Map_DSSP-1 | i | 1 | 2 | 3 | 4 |
|---|---|---|---|---|---|
| k | $\alpha_{ki}$ | Helix | Beta | Bridge | Other |
| 1 | 3H | 1 | 0 | 0 | 0 |
| 2 | 4H | 1 | 0 | 0 | 0 |
| 3 | 5H | 0 | 1 | 0 | 0 |
| 4 | BS | 0 | 1 | 0 | 0 |
| 5 | BB | 0 | 0 | 1 | 0 |
| 6 | TU | 0 | 0 | 0 | 1 |
| 7 | BE | 0 | 0 | 0 | 1 |
| 8 | UC | 0 | 0 | 0 | 1 |

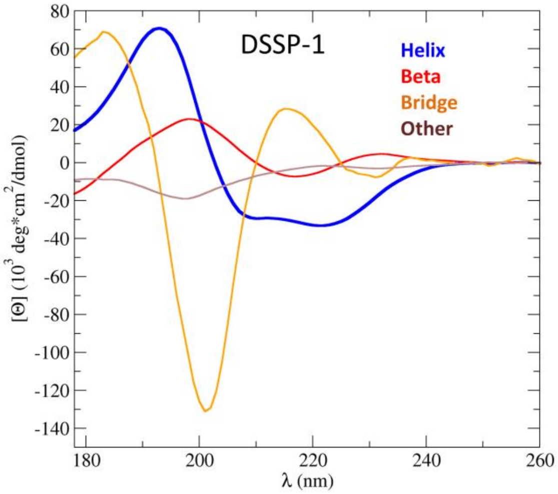


**Figure S9:** Assignment (left) and basis spectra (right) for the basis set DSSP-1. Secondary structure elements for the assignment are abbreviated according to Table S4.

| Map_DS3-3 | i | 1 | 2 | 3 | 4 |
|---|---|---|---|---|---|
| K | $\alpha_{ki}$ | Helix | Strand | Turns | Other |
| 1 | 3HT | 0 | 0 | 1 | 0 |
| 2 | HEL | 1 | 0 | 0 | 0 |
| 3 | BS | 0 | 1 | 0 | 0 |
| 4 | IRB | 0 | 0 | 1 | 0 |
| 5 | BT | 0 | 0 | 1 | 0 |
| 6 | OTT | 0 | 0 | 0 | 1 |
| 7 | LHT | 1 | 0 | 0 | 0 |
| 8 | UC | 0 | 0 | 0 | 1 |

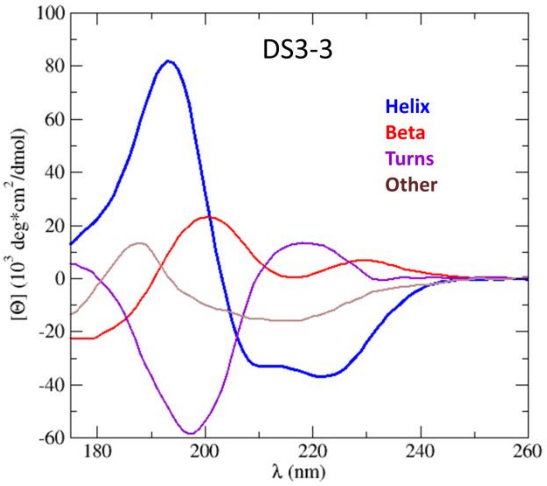


**Figure S10:** Assignment (left) and basis spectra (right) for the basis set DS3-3. Secondary structure elements for the assignment are abbreviated according to Table S5.

| Map_DS-dT | i | 1 | 2 | 3 |
|---|---|---|---|---|
| k | $\alpha_{ki}$ | Helix | Beta | Coil |
| 1 | 3H | 1 | 0 | 0 |
| 2 | ALH | 1 | 0 | 0 |
| 3 | PIH | 1 | 0 | 0 |
| 4 | EBS | 0 | 1 | 0 |
| 5 | NBS | 0 | 1 | 0 |
| 6 | PP | 0 | 0 | 1 |
| 7 | TI | 0 | 0 | 1 |
| 8 | TII | 0 | 0 | 1 |
| 9 | TVIII | 0 | 0 | 1 |
| 10 | GXT | 0 | 0 | 1 |
| 11 | SCH | 0 | 0 | 1 |
| 12 | HP | 0 | 0 | 1 |
| 13 | TC | 0 | 0 | 1 |
| 14 | HC | 0 | 0 | 1 |
| 15 | BC | 1 | 0 | 1 |
| 16 | BU | 0 | 0 | 1 |
| 17 | LTII | 0 | 0 | 1 |
| 18 | LHH | 0 | 0 | 1 |
| 19 | UC | 0 | 0 | 1 |

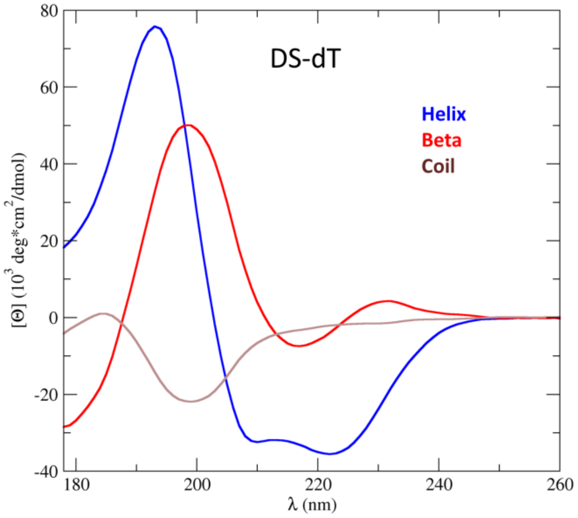


**Figure S11:** Assignment (left) and basis spectra (right) for the basis set DS-dT. Secondary structure elements for the assignment are abbreviated according to Table S5.

| Map_DS5-4 | i | 1 | 2 | 3 | 4 | 5 | 6 |
|---|---|---|---|---|---|---|---|
| k | $\alpha_{ki}$ | Helix-1 | Beta-1 | Helix-2 | Turn-1 | Turn-2 | Other |
| 1 | 3H | 0 | 0 | 1 | 0 | 0 | 0 |
| 2 | ALH | 1 | 0 | 0 | 0 | 0 | 0 |
| 3 | PIH | 0 | 0 | 0 | 0 | 1 | 0 |
| 4 | EBS | 0 | 1 | 0 | 0 | 0 | 0 |
| 5 | NBS | 0 | 1 | 0 | 0 | 0 | 0 |
| 6 | PP | 0 | 0 | 0 | 1 | 0 | 0 |
| 7 | TI | 0 | 0 | 0 | 1 | 0 | 0 |
| 8 | TII | 0 | 0 | 1 | 0 | 0 | 0 |
| 9 | TVIII | 1 | 0 | 0 | 0 | 0 | 0 |
| 10 | GXT | 0 | 0 | 0 | 1 | 0 | 0 |
| 11 | SCH | 0 | 0 | 0 | 0 | 1 | 0 |
| 12 | HP | 0 | 0 | 0 | 0 | 1 | 0 |
| 13 | TC | 0 | 0 | 0 | 0 | 0 | 1 |
| 14 | HC | 0 | 0 | 0 | 0 | 1 | 0 |
| 15 | BC | 0 | 0 | 0 | 0 | 1 | 0 |
| 16 | BU | 0 | 0 | 0 | 1 | 0 | 0 |
| 17 | LTII | 0 | 0 | 0 | 0 | 1 | 0 |
| 18 | LHH | 0 | 0 | 0 | 0 | 1 | 0 |
| 19 | UC | 0 | 0 | 0 | 0 | 0 | 1 |

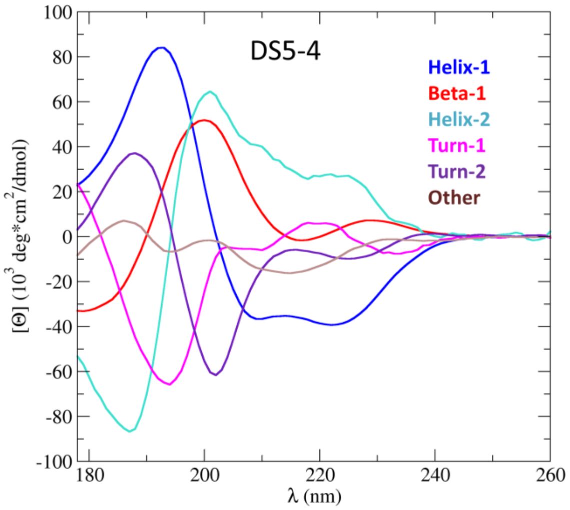


**Figure S12:** Assignment (left) and basis spectra (right) for the basis set DS5-4. Secondary structure elements for the assignment are abbreviated according to Table S5.

| Map_DS6-1 | i | 1 | 2 | 3 | 4 | 5 | 6 |
|---|---|---|---|---|---|---|---|
| k | $\alpha_{ki}$ | HEL1 | HEL2 | BST1 | BST2 | TRN1 | COIL |
| 1 | 3H | 0 | 0 | 1 | 0 | 0 | 0 |
| 2 | ALH | 1 | 0 | 0 | 0 | 0 | 0 |
| 3 | PIH | 0 | 1 | 0 | 0 | 0 | 0 |
| 4 | EBS | 0 | 0 | 0 | 1 | 0 | 0 |
| 5 | NBS | 0 | 0 | 1 | 0 | 0 | 0 |
| 6 | PP | 0 | 1 | 0 | 0 | 0 | 0 |
| 7 | TI | 0 | 0 | 0 | 0 | 1 | 0 |
| 8 | TII | 0 | 0 | 0 | 0 | 0 | 1 |
| 9 | TVIII | 0 | 0 | 0 | 0 | 0 | 1 |
| 10 | GXT | 0 | 0 | 1 | 0 | 0 | 0 |
| 11 | SCH | 0 | 1 | 0 | 0 | 0 | 0 |
| 12 | HP | 0 | 0 | 0 | 1 | 0 | 0 |
| 13 | TC | 1 | 0 | 0 | 0 | 0 | 0 |
| 14 | HC | 1 | 0 | 0 | 0 | 0 | 0 |
| 15 | BC | 0 | 0 | 0 | 0 | 1 | 0 |
| 16 | BU | 0 | 0 | 0 | 0 | 1 | 0 |
| 17 | LTII | 0 | 0 | 0 | 0 | 1 | 0 |
| 18 | LHH | 1 | 0 | 0 | 0 | 0 | 0 |
| 19 | UC | 0 | 0 | 0 | 0 | 0 | 1 |

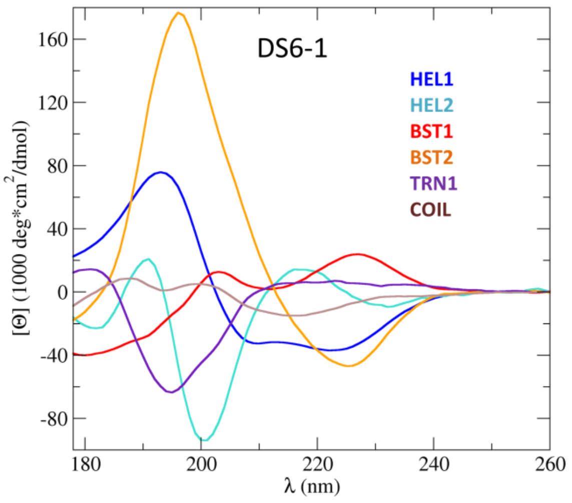


**Figure S13:** Assignment (left) and basis spectra (right) for the basis set DS6-1. Secondary structure elements for the assignment are abbreviated according to Table S5.

| Map_HBSS-4 | i | 1 | 2 | 3 | 4 |
|---|---|---|---|---|---|
| k | $\alpha_{ki}$ | Helix | BetaP | BetaA | Other |
| 1 | 4H | 1 | 0 | 0 | 0 |
| 2 | 3H | 1 | 0 | 0 | 0 |
| 3 | 5H | 1 | 0 | 0 | 0 |
| 4 | BSP | 0 | 1 | 0 | 0 |
| 8 | BSA | 0 | 0 | 1 | 0 |
| 10 | TU | 0 | 0 | 0 | 1 |
| 11 | UNC | 0 | 0 | 0 | 1 |

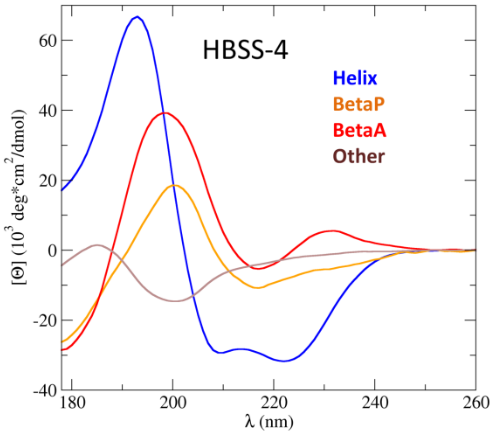


**Figure S14:** Assignment (left) and basis spectra (right) for the basis set HBSS-4. Secondary structure elements for the assignment are abbreviated according to Table S6.

| Map_HBSS-3 | i | 1 | 2 | 3 | 4 | 5 |
|---|---|---|---|---|---|---|
| k | $\alpha_{ki}$ | Hel_reg | Hel_irr | Beta_all | Turn | Other |
| 1 | 4H | 1 | 0 | 0 | 0 | 0 |
| 2 | 3H | 0 | 1 | 0 | 0 | 0 |
| 3 | 5H | 0 | 1 | 0 | 0 | 0 |
| 4 | LHA | 0 | 0 | 1 | 0 | 0 |
| 5 | NBA | 0 | 0 | 1 | 0 | 0 |
| 6 | RHA | 0 | 0 | 1 | 0 | 0 |
| 7 | LHP | 0 | 0 | 1 | 0 | 0 |
| 8 | NBP | 0 | 0 | 1 | 0 | 0 |
| 9 | RHP | 0 | 0 | 1 | 0 | 0 |
| 10 | TU | 0 | 0 | 0 | 1 | 0 |
| 11 | UNC | 0 | 0 | 0 | 0 | 1 |

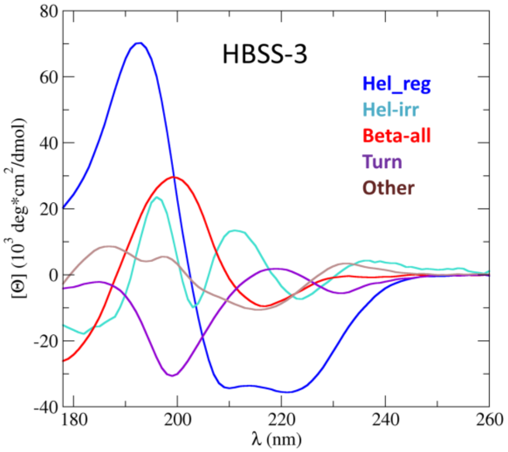


**Figure S15:** Assignment (left) and basis spectra (right) for the basis set HBSS-3. Secondary structure elements for the assignment are abbreviated according to Table S6.

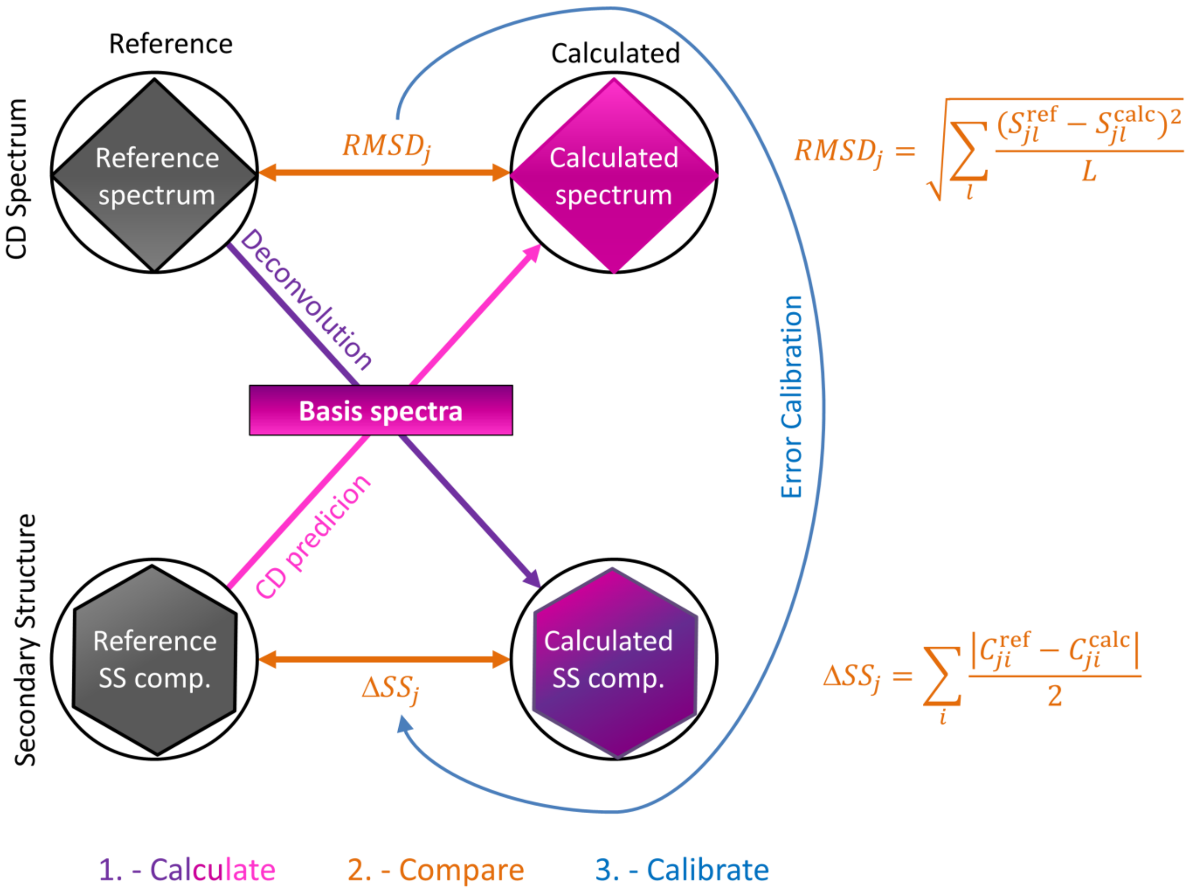


**Figure S16:** Calibration of the total model error. To estimate the error in the SS composition of structural models ($\Delta SS_j$) based on the error of their predicted CD spectrum ($RMSD_j$), first CD spectra is predicted from the SS composition of reference structures, and ideal SS compositions are predicted (via deconvolution) from the available reference CD spectra for each protein $j$. Then, the reference spectra and SS compositions are compared to the predicted ones to obtain $RMSD_j$ vs. $\Delta SS_j$ pairs for each protein, which are fitted with an error model during an error calibration step (Section S8). The resulting error model can be used to estimate $\Delta SS_j$ from CD predictions using the same basis spectra.

| BB_DS-dTSC3 | i | 1 | 2 | 3 |
|---|---|---|---|---|
| k | $\alpha_{ki}$ | Alpha | Beta | Coil |
| 1 | 3H | 1 | 0 | 0 |
| 2 | ALH | 1 | 0 | 0 |
| 3 | PIH | 1 | 0 | 0 |
| 4 | EBS | 0 | 1 | 0 |
| 5 | NBS | 0 | 1 | 0 |
| 6 | PP | 0 | 0 | 1 |
| 7 | TI | 0 | 0 | 1 |
| 8 | TII | 0 | 0 | 1 |
| 9 | TVIII | 0 | 0 | 1 |
| 10 | GXT | 0 | 0 | 1 |
| 11 | SCH | 0 | 0 | 1 |
| 12 | HP | 0 | 0 | 1 |
| 13 | TC | 0 | 0 | 1 |
| 14 | HC | 0 | 0 | 1 |
| 15 | BC | 0 | 0 | 1 |
| 16 | BU | 0 | 0 | 1 |
| 17 | LTII | 0 | 0 | 1 |
| 18 | LHH | 0 | 0 | 1 |
| 19 | UC | 0 | 0 | 1 |

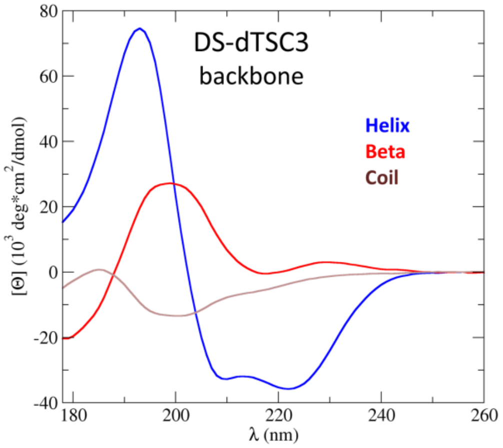


| SC_DS-dTSC3 | i | 1 | 2 | 3 | 4 |
|---|---|---|---|---|---|
| k | $\alpha_k$ | GRP1 | GRP2 | GRP3 | GRP4 |
| 1 | ALA | 0 | 0 | 0 | 1 |
| 2 | ASP | 0 | 1 | 0 | 0 |
| 3 | ASN | 0 | 1 | 0 | 0 |
| 4 | ARG | 0 | 1 | 0 | 0 |
| 5 | CYS | 0 | 0 | 0 | 1 |
| 6 | GLY | 0 | 0 | 1 | 0 |
| 7 | GLU | 0 | 1 | 0 | 0 |
| 8 | GLN | 0 | 1 | 0 | 0 |
| 9 | HIS | 1 | 0 | 0 | 0 |
| 10 | ILE | 0 | 0 | 0 | 1 |
| 11 | LEU | 0 | 0 | 0 | 1 |
| 12 | LYS | 0 | 0 | 0 | 1 |
| 13 | MET | 0 | 0 | 0 | 1 |
| 14 | PHE | 1 | 0 | 0 | 0 |
| 15 | PRO | 0 | 0 | 0 | 1 |
| 16 | SER | 0 | 0 | 0 | 1 |
| 17 | THR | 0 | 0 | 0 | 1 |
| 18 | TRP | 1 | 0 | 0 | 0 |
| 19 | TYR | 1 | 0 | 0 | 0 |
| 20 | VAL | 0 | 0 | 0 | 1 |

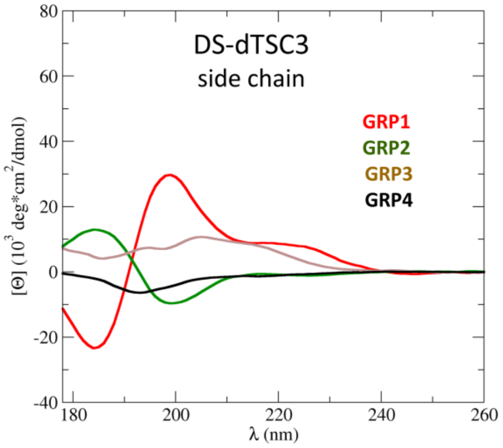


**Figure S17:** Assignment (left) and basis spectra (right) for the mixed basis set DS-dTSC3. The backbone and side-chain related basis spectra are shown on the top and bottom, respectively. Secondary structure elements for the assignment are abbreviated according to Table S5.

| BB_DS6-1SC1 | i | 1 | 2 | 3 | 4 | 5 | 6 |
|---|---|---|---|---|---|---|---|
| k | $\alpha_{ki}$ | Helix-1 | Helix-2 | Beta-1 | Beta-2 | Turn | Coil |
| 1 | 3H | 0 | 0 | 1 | 0 | 0 | 0 |
| 2 | ALH | 1 | 0 | 0 | 0 | 0 | 0 |
| 3 | PIH | 0 | 1 | 0 | 0 | 0 | 0 |
| 4 | EBS | 0 | 0 | 0 | 1 | 0 | 0 |
| 5 | NBS | 0 | 0 | 1 | 0 | 0 | 0 |
| 6 | PP | 0 | 1 | 0 | 0 | 0 | 0 |
| 7 | TI | 0 | 0 | 0 | 0 | 1 | 0 |
| 8 | TII | 0 | 0 | 0 | 0 | 0 | 1 |
| 9 | TVIII | 0 | 0 | 0 | 0 | 0 | 1 |
| 10 | GXT | 0 | 0 | 1 | 0 | 0 | 0 |
| 11 | SCH | 0 | 1 | 0 | 0 | 0 | 0 |
| 12 | HP | 0 | 0 | 0 | 1 | 0 | 0 |
| 13 | TC | 1 | 0 | 0 | 0 | 0 | 0 |
| 14 | HC | 1 | 0 | 0 | 0 | 0 | 0 |
| 15 | BC | 0 | 0 | 0 | 0 | 1 | 0 |
| 16 | BU | 0 | 0 | 0 | 0 | 1 | 0 |
| 17 | LTII | 0 | 0 | 0 | 0 | 1 | 0 |
| 18 | LHH | 1 | 0 | 0 | 0 | 0 | 0 |
| 19 | UC | 0 | 0 | 0 | 0 | 0 | 1 |

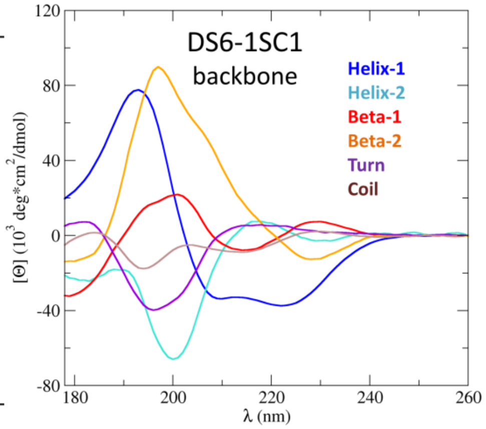


| SC_DS6-1SC1 | I | 1 | 2 | 3 | 4 | 5 | 6 |
|---|---|---|---|---|---|---|---|
| k | $\alpha_{ki}$ | GRP1 | GRP2 | GRP3 | GRP4 | GRP5 | GRP6 |
| 1 | ALA | 0 | 0 | 0 | 0 | 1 | 0 |
| 2 | ASP | 1 | 0 | 0 | 0 | 0 | 0 |
| 3 | ASN | 0 | 0 | 1 | 0 | 0 | 0 |
| 4 | ARG | 0 | 0 | 0 | 0 | 0 | 1 |
| 5 | CYS | 0 | 1 | 0 | 0 | 0 | 0 |
| 6 | GLY | 0 | 0 | 0 | 0 | 0 | 1 |
| 7 | GLU | 0 | 0 | 1 | 0 | 0 | 0 |
| 8 | GLN | 0 | 0 | 0 | 1 | 0 | 0 |
| 9 | HIS | 0 | 0 | 0 | 1 | 0 | 0 |
| 10 | ILE | 0 | 0 | 1 | 0 | 0 | 0 |
| 11 | LEU | 0 | 0 | 0 | 0 | 1 | 0 |
| 12 | LYS | 0 | 0 | 0 | 1 | 0 | 0 |
| 13 | MET | 0 | 0 | 0 | 1 | 0 | 0 |
| 14 | PHE | 1 | 0 | 0 | 0 | 0 | 0 |
| 15 | PRO | 0 | 0 | 0 | 0 | 1 | 0 |
| 16 | SER | 0 | 0 | 1 | 0 | 0 | 0 |
| 17 | THR | 0 | 1 | 0 | 0 | 0 | 0 |
| 18 | TRP | 0 | 1 | 0 | 0 | 0 | 0 |
| 19 | TYR | 0 | 0 | 0 | 0 | 0 | 1 |
| 20 | VAL | 1 | 0 | 0 | 0 | 0 | 0 |

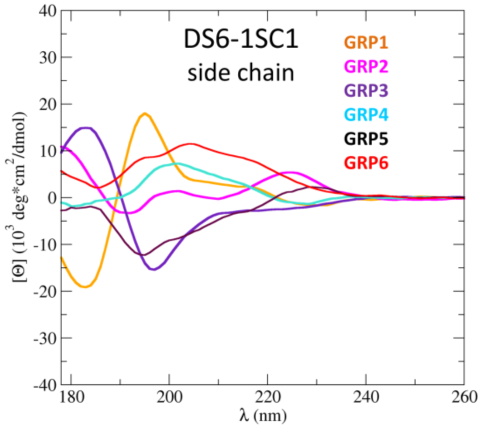


**Figure S18:** Assignment (left) and basis spectra (right) for the mixed basis set DS6-1SC1. The backbone and side-chain related basis spectra are shown on the top and bottom, respectively. Secondary structure elements for the assignment are abbreviated according to Table S5.

| BB_DS5-4SC1 | i | 1 | 2 | 3 | 4 | 5 | 6 |
|---|---|---|---|---|---|---|---|
| k | $\alpha_{ki}$ | Helix-1 | Beta-1 | Helix-2 | Turn-1 | Turn-2 | Other |
| 1 | 3H | 0 | 0 | 1 | 0 | 0 | 0 |
| 2 | ALH | 1 | 0 | 0 | 0 | 0 | 0 |
| 3 | PIH | 0 | 0 | 0 | 0 | 1 | 0 |
| 4 | EBS | 0 | 1 | 0 | 0 | 0 | 0 |
| 5 | NBS | 0 | 1 | 0 | 0 | 0 | 0 |
| 6 | PP | 0 | 0 | 0 | 1 | 0 | 0 |
| 7 | TI | 0 | 0 | 0 | 1 | 0 | 0 |
| 8 | TII | 0 | 0 | 1 | 0 | 0 | 0 |
| 9 | TVIII | 1 | 0 | 0 | 0 | 0 | 0 |
| 10 | GXT | 0 | 0 | 0 | 1 | 0 | 0 |
| 11 | SCH | 0 | 0 | 0 | 0 | 1 | 0 |
| 12 | HP | 0 | 0 | 0 | 0 | 1 | 0 |
| 13 | TC | 0 | 0 | 0 | 0 | 0 | 1 |
| 14 | HC | 0 | 0 | 0 | 0 | 1 | 0 |
| 15 | BC | 0 | 0 | 0 | 0 | 1 | 0 |
| 16 | BU | 0 | 0 | 0 | 1 | 0 | 0 |
| 17 | LTII | 0 | 0 | 0 | 0 | 1 | 0 |
| 18 | LHH | 0 | 0 | 0 | 0 | 1 | 0 |
| 19 | UC | 0 | 0 | 0 | 0 | 0 | 1 |

| SC_DS5-4SC1 | i | 1 | 2 | 3 | 4 | 5 | 6 |
|---|---|---|---|---|---|---|---|
| k | $\alpha_{ki}$ | GRP1 | GRP2 | GRP3 | GRP4 | GRP5 | GRP6 |
| 1 | ALA | 0 | 0 | 0 | 0 | 1 | 0 |
| 2 | ASP | 0 | 0 | 0 | 1 | 0 | 0 |
| 3 | ASN | 0 | 0 | 0 | 0 | 1 | 0 |
| 4 | ARG | 1 | 0 | 0 | 0 | 0 | 0 |
| 5 | CYS | 0 | 0 | 0 | 1 | 0 | 0 |
| 6 | GLY | 0 | 0 | 0 | 0 | 0 | 1 |
| 7 | GLU | 0 | 0 | 1 | 0 | 0 | 0 |
| 8 | GLN | 0 | 0 | 0 | 1 | 0 | 0 |
| 9 | HIS | 0 | 1 | 0 | 0 | 0 | 0 |
| 10 | ILE | 1 | 0 | 0 | 0 | 0 | 0 |
| 11 | LEU | 0 | 0 | 1 | 0 | 0 | 0 |
| 12 | LYS | 0 | 0 | 0 | 0 | 0 | 1 |
| 13 | MET | 0 | 0 | 0 | 0 | 0 | 1 |
| 14 | PHE | 0 | 1 | 0 | 0 | 0 | 0 |
| 15 | PRO | 0 | 0 | 0 | 0 | 1 | 0 |
| 16 | SER | 1 | 0 | 0 | 0 | 0 | 0 |
| 17 | THR | 0 | 0 | 0 | 0 | 1 | 0 |
| 18 | TRP | 1 | 0 | 0 | 0 | 0 | 0 |
| 19 | TYR | 0 | 1 | 0 | 0 | 0 | 0 |
| 20 | VAL | 0 | 0 | 0 | 0 | 0 | 1 |

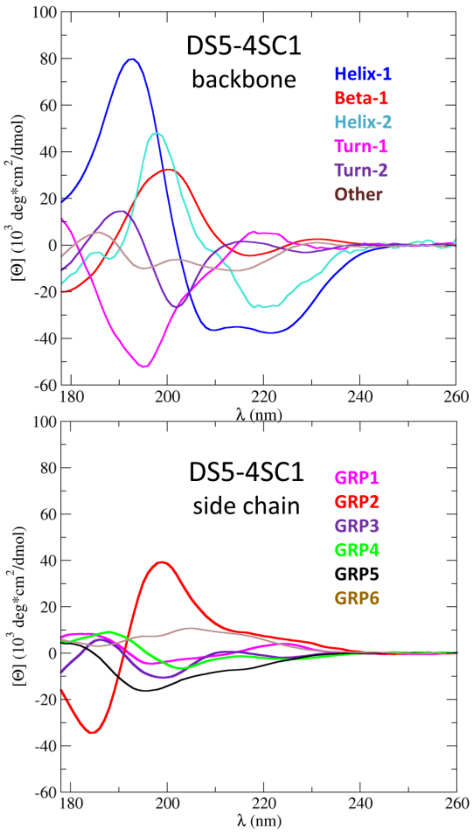


**Figure S19:** Assignment (left) and basis spectra (right) for the mixed basis set DS5-4SC1. The backbone and side-chain related basis spectra are shown on the top and bottom, respectively. Secondary structure elements for the assignment are abbreviated according to Table S5.

| BB_DSSP-1SC3 | i | 1 | 2 | 3 | 4 |
|---|---|---|---|---|---|
| k | $\alpha_{ki}$ | Helix | Sheet | Bridge | Other |
| 1 | 3H | 1 | 0 | 0 | 0 |
| 2 | 4H | 1 | 0 | 0 | 0 |
| 3 | 5H | 0 | 1 | 0 | 0 |
| 4 | BS | 0 | 1 | 0 | 0 |
| 5 | BB | 0 | 0 | 1 | 0 |
| 6 | TU | 0 | 0 | 0 | 1 |
| 7 | BE | 0 | 0 | 0 | 1 |
| 8 | UC | 0 | 0 | 0 | 1 |

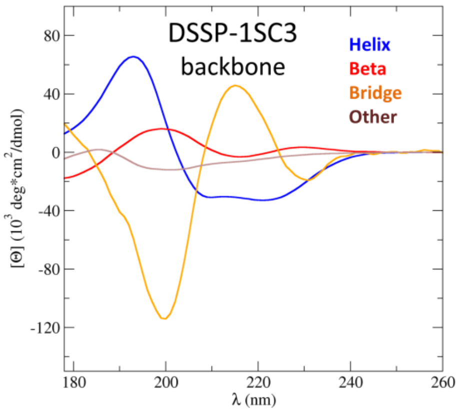


| SC_DSSP-1SC3 | i | 1 | 2 | 3 | 4 | 5 | 6 |
|---|---|---|---|---|---|---|---|
| k | $\alpha_{ki}$ | GRP1 | GRP2 | GRP3 | GRP4 | GRP5 | GRP6 |
| 1 | ALA | 0 | 0 | 0 | 0 | 1 | 0 |
| 2 | ASP | 0 | 0 | 0 | 0 | 0 | 1 |
| 3 | ASN | 0 | 0 | 0 | 1 | 0 | 0 |
| 4 | ARG | 0 | 0 | 0 | 1 | 0 | 0 |
| 5 | CYS | 1 | 0 | 0 | 0 | 0 | 0 |
| 6 | GLY | 0 | 0 | 0 | 0 | 1 | 0 |
| 7 | GLU | 1 | 0 | 0 | 0 | 0 | 0 |
| 8 | GLN | 1 | 0 | 0 | 0 | 0 | 0 |
| 9 | HIS | 0 | 0 | 0 | 0 | 1 | 0 |
| 10 | ILE | 0 | 1 | 0 | 0 | 0 | 0 |
| 11 | LEU | 0 | 0 | 0 | 0 | 0 | 1 |
| 12 | LYS | 0 | 0 | 0 | 0 | 0 | 1 |
| 13 | MET | 0 | 0 | 0 | 0 | 0 | 1 |
| 14 | PHE | 0 | 0 | 0 | 0 | 1 | 0 |
| 15 | PRO | 0 | 0 | 1 | 0 | 0 | 0 |
| 16 | SER | 0 | 0 | 0 | 0 | 1 | 0 |
| 17 | THR | 0 | 0 | 1 | 0 | 0 | 0 |
| 18 | TRP | 0 | 0 | 0 | 1 | 0 | 0 |
| 19 | TYR | 0 | 0 | 1 | 0 | 0 | 0 |
| 20 | VAL | 0 | 0 | 0 | 1 | 0 | 0 |

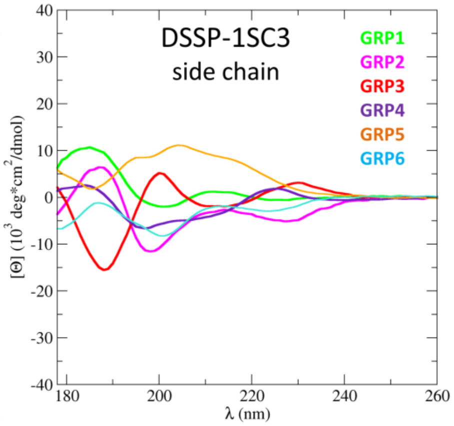


**Figure S20:** Assignment (left) and basis spectra (right) for the mixed basis set DSSP-1SC3. The backbone and side-chain related basis spectra are shown on the top and bottom, respectively. Secondary structure elements for the assignment are abbreviated according to Table S4.

| BB_HBSS-3SC1 | i | 1 | 2 | 3 | 4 | 5 |
|---|---|---|---|---|---|---|
| k | $\alpha_{ki}$ | Hel_reg | Hel_irr | Beta_all | Turn | Other |
| 1 | 4H | 1 | 0 | 0 | 0 | 0 |
| 2 | 3H | 0 | 1 | 0 | 0 | 0 |
| 3 | 5H | 0 | 1 | 0 | 0 | 0 |
| 4 | LHA | 0 | 0 | 1 | 0 | 0 |
| 5 | NBA | 0 | 0 | 1 | 0 | 0 |
| 6 | RHA | 0 | 0 | 1 | 0 | 0 |
| 7 | LHP | 0 | 0 | 1 | 0 | 0 |
| 8 | NBP | 0 | 0 | 1 | 0 | 0 |
| 9 | RHP | 0 | 0 | 1 | 0 | 0 |
| 10 | TU | 0 | 0 | 0 | 1 | 0 |
| 11 | UNC | 0 | 0 | 0 | 0 | 1 |

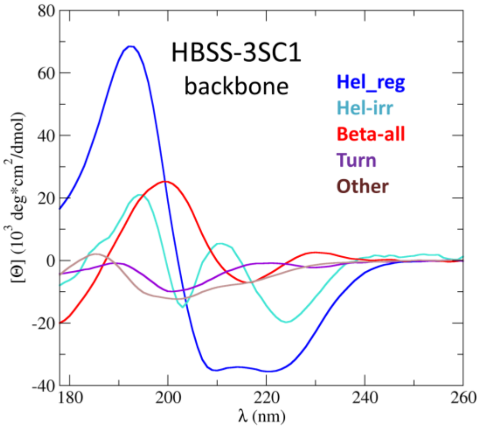


| SC_HBSS-3SC1 | i | 1 | 2 | 3 | 4 |
|---|---|---|---|---|---|
| k | $\alpha_{ki}$ | GRP1 | GRP2 | GRP3 | GRP4 |
| 1 | ALA | 0 | 0 | 0 | 1 |
| 2 | ASP | 0 | 1 | 0 | 0 |
| 3 | ASN | 0 | 1 | 0 | 0 |
| 4 | ARG | 0 | 1 | 0 | 0 |
| 5 | CYS | 0 | 0 | 0 | 1 |
| 6 | GLY | 0 | 0 | 1 | 0 |
| 7 | GLU | 0 | 1 | 0 | 0 |
| 8 | GLN | 0 | 1 | 0 | 0 |
| 9 | HIS | 1 | 0 | 0 | 0 |
| 10 | ILE | 0 | 0 | 0 | 1 |
| 11 | LEU | 0 | 0 | 0 | 1 |
| 12 | LYS | 0 | 0 | 0 | 1 |
| 13 | MET | 1 | 0 | 0 | 0 |
| 14 | PHE | 1 | 0 | 0 | 0 |
| 15 | PRO | 0 | 0 | 0 | 1 |
| 16 | SER | 0 | 1 | 0 | 0 |
| 17 | THR | 0 | 0 | 0 | 1 |
| 18 | TRP | 1 | 0 | 0 | 0 |
| 19 | TYR | 1 | 0 | 0 | 0 |
| 20 | VAL | 0 | 0 | 0 | 1 |

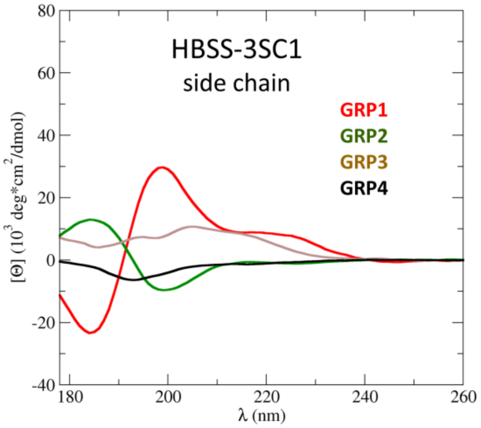


**Figure S21:** Assignment (left) and basis spectra (right) for the mixed basis set HBSS-3SC1. The backbone and side-chain related basis spectra are shown on the top and bottom, respectively. Secondary structure elements for the assignment are abbreviated according to Table S6.

| BB_HBSS-3SCF | i | 1 | 2 | 3 | 4 | 5 |
|---|---|---|---|---|---|---|
| k | $\alpha_{ki}$ | Hel-reg | Hel-irr | Beta-all | Turn | Other |
| 1 | 4H | 1 | 0 | 0 | 0 | 0 |
| 2 | 3H | 0 | 1 | 0 | 0 | 0 |
| 3 | 5H | 0 | 1 | 0 | 0 | 0 |
| 4 | LHA | 0 | 0 | 1 | 0 | 0 |
| 5 | NBA | 0 | 0 | 1 | 0 | 0 |
| 6 | RHA | 0 | 0 | 1 | 0 | 0 |
| 7 | LHP | 0 | 0 | 1 | 0 | 0 |
| 8 | NBP | 0 | 0 | 1 | 0 | 0 |
| 9 | RHP | 0 | 0 | 1 | 0 | 0 |
| 10 | TU | 0 | 0 | 0 | 1 | 0 |
| 11 | UNC | 0 | 0 | 0 | 0 | 1 |

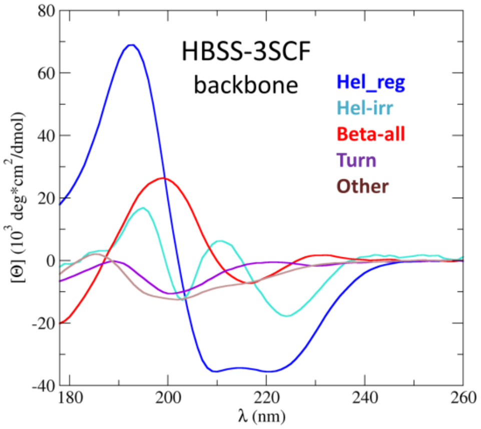


SC_HBSS-3SCF (Basis spectra abbreviated by 1-letter AA codes)

| | I | 1 | 2 | 3 | 4 | 5 | 6 | 7 | 8 | 9 | 10 | 11 | 12 | 13 | 14 | 15 | 16 | 17 | 18 | 19 | 20 |
|---|---|---|---|---|---|---|---|---|---|---|---|---|---|---|---|---|---|---|---|---|---|
| k | $\alpha_{ki}$ | A | D | N | R | C | G | E | Q | H | I | L | K | M | F | P | S | T | W | Y | V |
| 1 | ALA | 1 | 0 | 0 | 0 | 0 | 0 | 0 | 0 | 0 | 0 | 0 | 0 | 0 | 0 | 0 | 0 | 0 | 0 | 0 | 0 |
| 2 | ASP | 0 | 1 | 0 | 0 | 0 | 0 | 0 | 0 | 0 | 0 | 0 | 0 | 0 | 0 | 0 | 0 | 0 | 0 | 0 | 0 |
| 3 | ASN | 0 | 0 | 1 | 0 | 0 | 0 | 0 | 0 | 0 | 0 | 0 | 0 | 0 | 0 | 0 | 0 | 0 | 0 | 0 | 0 |
| 4 | ARG | 0 | 0 | 0 | 1 | 0 | 0 | 0 | 0 | 0 | 0 | 0 | 0 | 0 | 0 | 0 | 0 | 0 | 0 | 0 | 0 |
| 5 | CYS | 0 | 0 | 0 | 0 | 1 | 0 | 0 | 0 | 0 | 0 | 0 | 0 | 0 | 0 | 0 | 0 | 0 | 0 | 0 | 0 |
| 6 | GLY | 0 | 0 | 0 | 0 | 0 | 1 | 0 | 0 | 0 | 0 | 0 | 0 | 0 | 0 | 0 | 0 | 0 | 0 | 0 | 0 |
| 7 | GLU | 0 | 0 | 0 | 0 | 0 | 0 | 1 | 0 | 0 | 0 | 0 | 0 | 0 | 0 | 0 | 0 | 0 | 0 | 0 | 0 |
| 8 | GLN | 0 | 0 | 0 | 0 | 0 | 0 | 0 | 1 | 0 | 0 | 0 | 0 | 0 | 0 | 0 | 0 | 0 | 0 | 0 | 0 |
| 9 | HIS | 0 | 0 | 0 | 0 | 0 | 0 | 0 | 0 | 1 | 0 | 0 | 0 | 0 | 0 | 0 | 0 | 0 | 0 | 0 | 0 |
| 10 | ILE | 0 | 0 | 0 | 0 | 0 | 0 | 0 | 0 | 0 | 1 | 0 | 0 | 0 | 0 | 0 | 0 | 0 | 0 | 0 | 0 |
| 11 | LEU | 0 | 0 | 0 | 0 | 0 | 0 | 0 | 0 | 0 | 0 | 1 | 0 | 0 | 0 | 0 | 0 | 0 | 0 | 0 | 0 |
| 12 | LYS | 0 | 0 | 0 | 0 | 0 | 0 | 0 | 0 | 0 | 0 | 0 | 1 | 0 | 0 | 0 | 0 | 0 | 0 | 0 | 0 |
| 13 | MET | 0 | 0 | 0 | 0 | 0 | 0 | 0 | 0 | 0 | 0 | 0 | 0 | 1 | 0 | 0 | 0 | 0 | 0 | 0 | 0 |
| 14 | PHE | 0 | 0 | 0 | 0 | 0 | 0 | 0 | 0 | 0 | 0 | 0 | 0 | 0 | 1 | 0 | 0 | 0 | 0 | 0 | 0 |
| 15 | PRO | 0 | 0 | 0 | 0 | 0 | 0 | 0 | 0 | 0 | 0 | 0 | 0 | 0 | 0 | 1 | 0 | 0 | 0 | 0 | 0 |
| 16 | SER | 0 | 0 | 0 | 0 | 0 | 0 | 0 | 0 | 0 | 0 | 0 | 0 | 0 | 0 | 0 | 1 | 0 | 0 | 0 | 0 |
| 17 | THR | 0 | 0 | 0 | 0 | 0 | 0 | 0 | 0 | 0 | 0 | 0 | 0 | 0 | 0 | 0 | 0 | 1 | 0 | 0 | 0 |
| 18 | TRP | 0 | 0 | 0 | 0 | 0 | 0 | 0 | 0 | 0 | 0 | 0 | 0 | 0 | 0 | 0 | 0 | 0 | 1 | 0 | 0 |
| 19 | TYR | 0 | 0 | 0 | 0 | 0 | 0 | 0 | 0 | 0 | 0 | 0 | 0 | 0 | 0 | 0 | 0 | 0 | 0 | 1 | 0 |
| 20 | VAL | 0 | 0 | 0 | 0 | 0 | 0 | 0 | 0 | 0 | 0 | 0 | 0 | 0 | 0 | 0 | 0 | 0 | 0 | 0 | 1 |

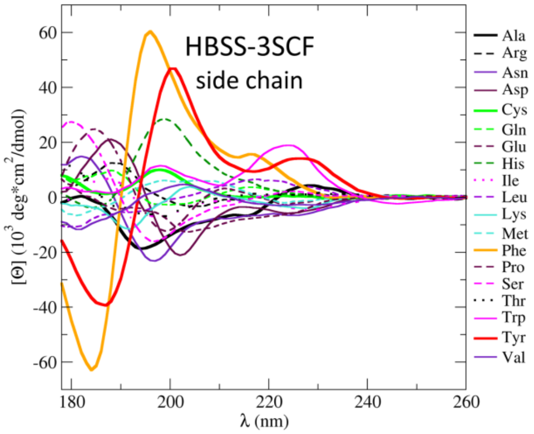


**Figure S22:** Assignment (left) and basis spectra (right) for the mixed basis set HBSS-3SCF. The backbone and side-chain related basis spectra are shown on the top and bottom, respectively. Secondary structure elements for the assignment are abbreviated according to Table S6.

| Set_Sreer-1 | i | 1 | 2 | 3 | 4 | 5 | 6 |
|---|---|---|---|---|---|---|---|
| k | $\alpha_k$ | Hel-reg | Hel-irr | Beta-reg | Bet-irr | Turn | Unord. |
| 1 | 3H | 0 | 1 | 0 | 0 | 0 | 0 |
| 2 | ALH | 1 | 0 | 0 | 0 | 0 | 0 |
| 3 | PIH | 1 | 0 | 0 | 0 | 0 | 0 |
| 4 | EBS | 0 | 0 | 1 | 0 | 0 | 0 |
| 5 | NBS | 0 | 0 | 1 | 0 | 0 | 0 |
| 6 | PP | 0 | 0 | 0 | 0 | 0 | 1 |
| 7 | TI | 0 | 1 | 0 | 0 | 0 | 0 |
| 8 | TII | 0 | 0 | 0 | 0 | 1 | 0 |
| 9 | TVIII | 0 | 0 | 0 | 0 | 1 | 0 |
| 10 | GXT | 0 | 0 | 0 | 0 | 1 | 0 |
| 11 | SCH | 0 | 1 | 0 | 0 | 0 | 0 |
| 12 | HP | 0 | 0 | 0 | 1 | 0 | 0 |
| 13 | TC | 0 | 0 | 0 | 0 | 1 | 0 |
| 14 | HC | 0 | 1 | 0 | 0 | 0 | 0 |
| 15 | BC | 0 | 0 | 0 | 1 | 0 | 0 |
| 16 | BU | 0 | 0 | 0 | 1 | 0 | 0 |
| 17 | LTII | 0 | 0 | 0 | 0 | 1 | 0 |
| 18 | LHH | 0 | 0 | 0 | 0 | 1 | 0 |
| 19 | UC | 0 | 0 | 0 | 0 | 0 | 1 |

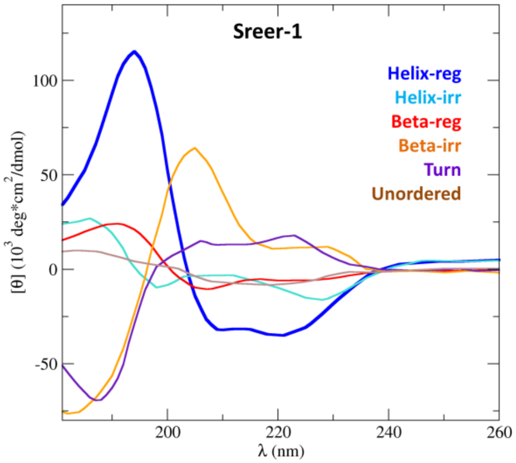


**Figure S23:** Assignment (left) and basis spectra (right) for the adapted basis set Sreer-1. Secondary structure elements for the assignment are abbreviated according to Table S5.

| Map_DSSP-1 | i | 1 | 2 | 3 | 4 | 5 |
|---|---|---|---|---|---|---|
| k | $\alpha_{ki}$ | Helix | Beta | Turn-1 | Turn-2 | Aper. |
| 1 | 3H | 0 | 0 | 1 | 0 | 0 |
| 2 | 4H | 1 | 0 | 0 | 0 | 0 |
| 3 | 5H | 0 | 0 | 0 | 0 | 1 |
| 4 | BS | 0 | 1 | 0 | 0 | 0 |
| 5 | BB | 0 | 0 | 0 | 0 | 1 |
| 6 | TU | 0 | 0 | 0 | 1 | 0 |
| 7 | BE | 0 | 0 | 0 | 0 | 1 |
| 8 | UC | 0 | 0 | 0 | 0 | 1 |

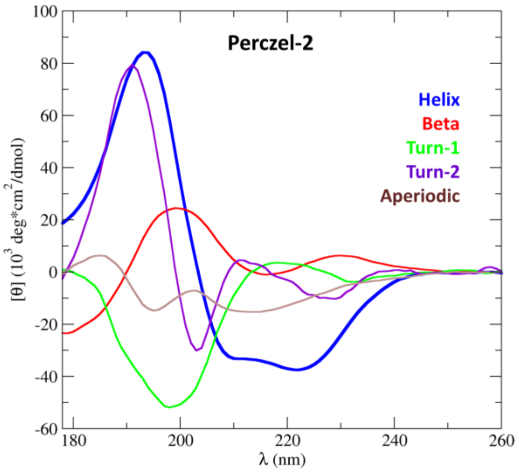


**Figure S24:** Assignment (left) and basis spectra (right) for the adapted basis set Preczel-2. Secondary structure elements for the assignment are abbreviated according to Table S4.

| Set_BestSel-2 | i | 1 | 2 | 3 | 4 | 5 | 6 | 7 | 8 |
|---|---|---|---|---|---|---|---|---|---|
| k | $\alpha_{ki}$ | Hel-reg | Hel-irr | Anti1 | Anti2 | Anti3 | Para | Turn | Other |
| 1 | 4H | 1 | 0 | 0 | 0 | 0 | 0 | 0 | 0 |
| 2 | 3H | 0 | 1 | 0 | 0 | 0 | 0 | 0 | 0 |
| 3 | 5H | 0 | 1 | 0 | 0 | 0 | 0 | 0 | 0 |
| 4 | LHA | 0 | 0 | 1 | 0 | 0 | 0 | 0 | 0 |
| 5 | NBA | 0 | 0 | 0 | 1 | 0 | 0 | 0 | 0 |
| 6 | RHA | 0 | 0 | 0 | 0 | 1 | 0 | 0 | 0 |
| 7 | LHP | 0 | 0 | 0 | 0 | 0 | 1 | 0 | 0 |
| 8 | NBP | 0 | 0 | 0 | 0 | 0 | 1 | 0 | 0 |
| 9 | RHP | 0 | 0 | 0 | 0 | 0 | 1 | 0 | 0 |
| 10 | TU | 0 | 0 | 0 | 0 | 0 | 0 | 1 | 0 |
| 11 | UNC | 0 | 0 | 0 | 0 | 0 | 0 | 0 | 1 |

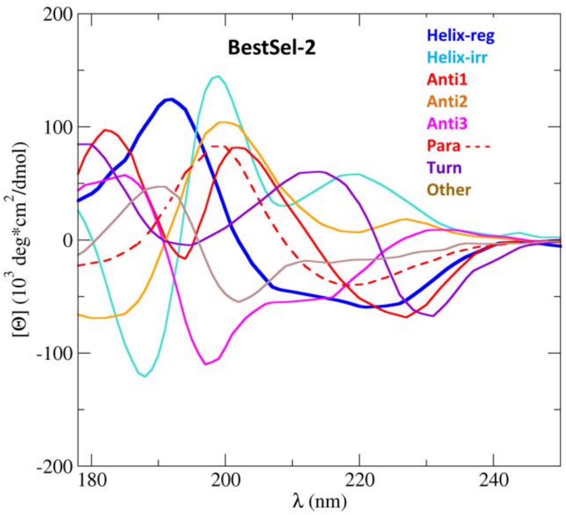


**Figure S25:** Assignment (left) and basis spectra (right) for the adapted basis set BestSel-2. Derived previously by Micsonai *et al.* [3] Secondary structure elements for the assignment are abbreviated according to Table S6.

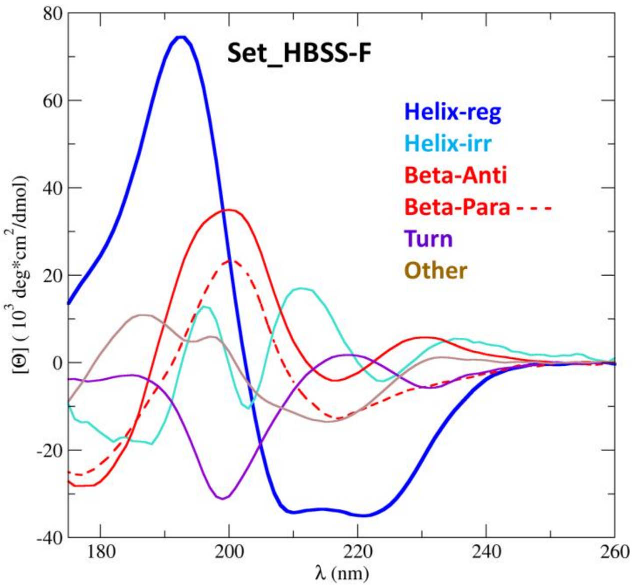


**Figure S26:** Derived initial basis spectra for the structure classification HbSS, basic classification library.

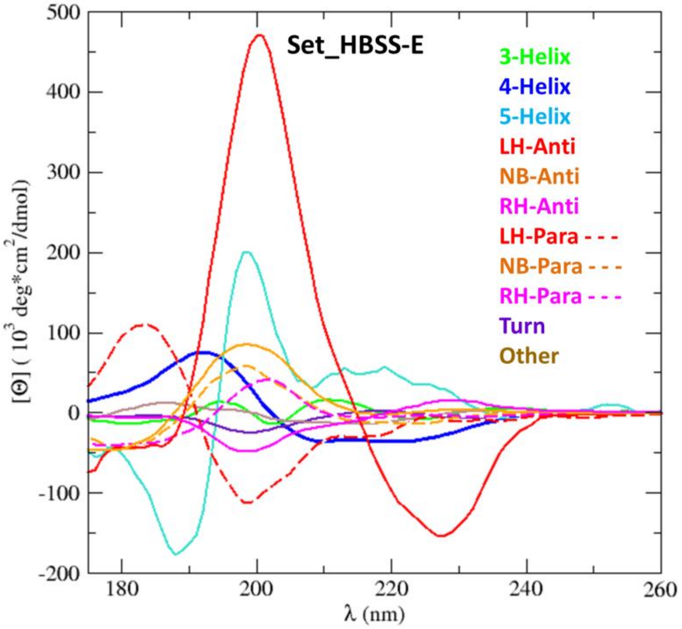


**Figure S27:** Derived initial basis spectra for the structure classification algorithm HbSS, extended library (HbSS_ext).

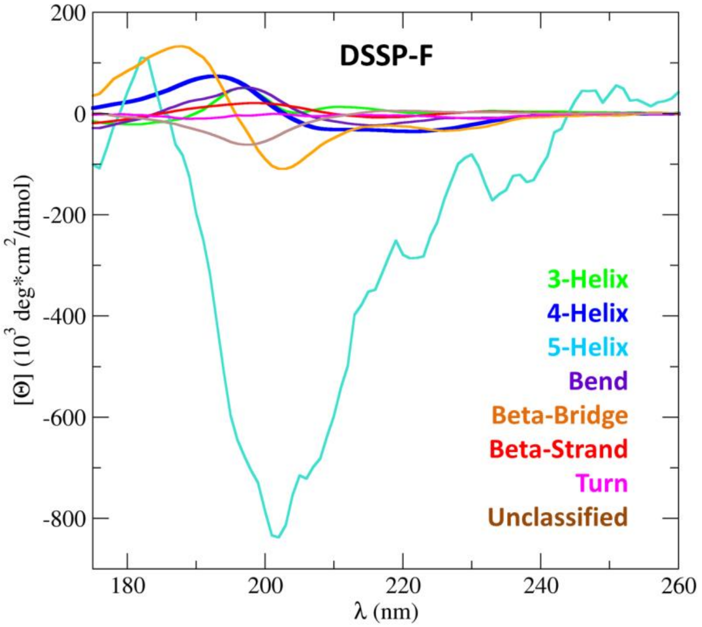


**Figure S28:** Derived initial basis spectra for the structure classification DSSP.

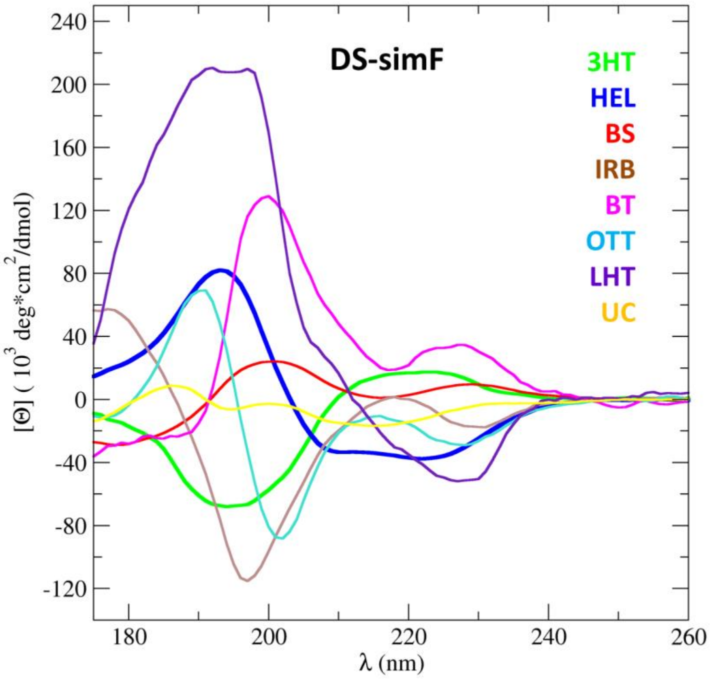


**Figure S29:** Derived initial basis spectra for the structure classification DISICL, simplified classification library (DS_sim).

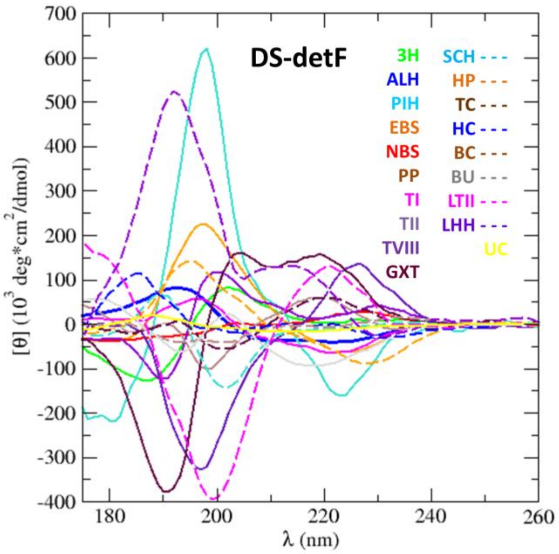


**Figure S30:** Derived initial basis spectra for the structure classification DISICL, detailed classification library (DS_det).

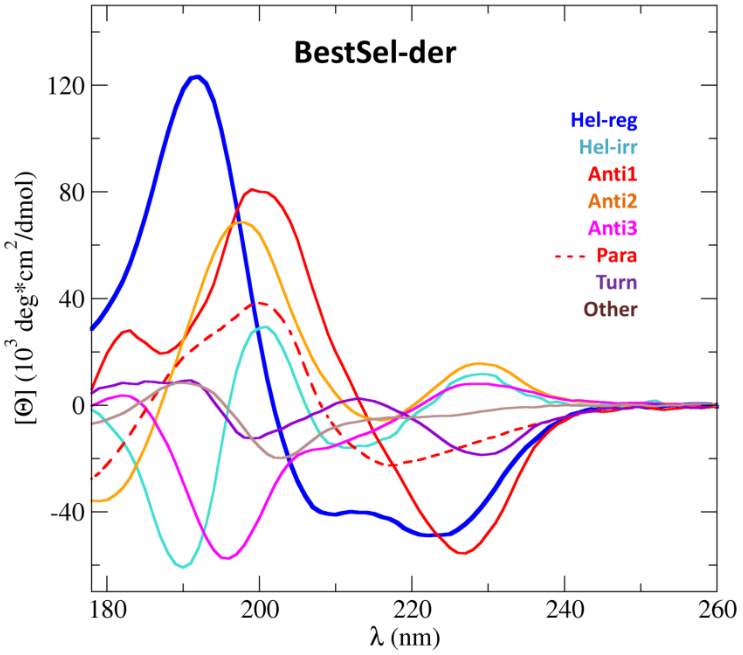


**Figure S31:** derived basis spectra for the BestSel deconvolution algorithm.

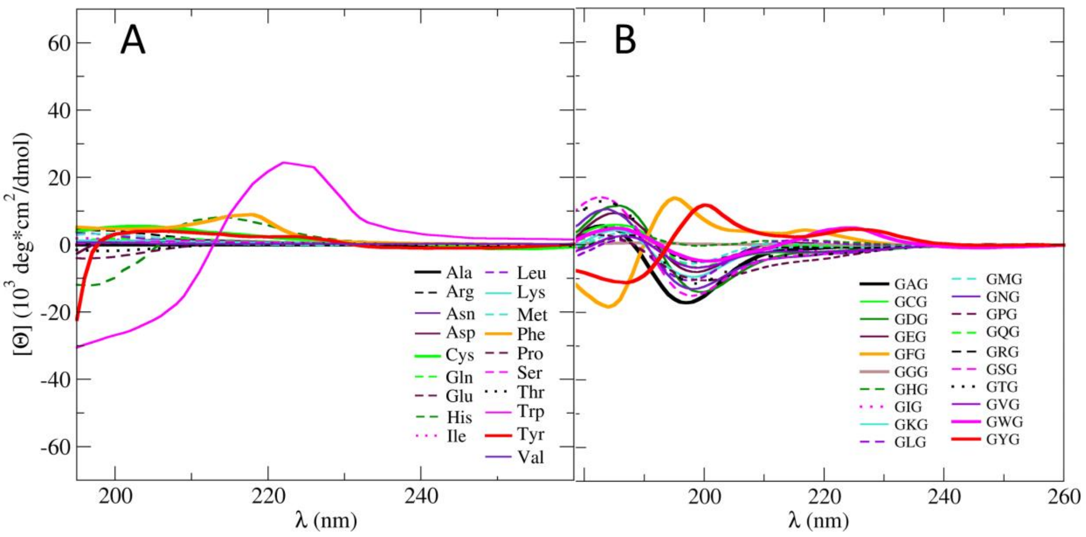


**Figure S32:** Comparison between the CD spectra of A) isolated natural amino acids and B) of Ac-GXG-NH2 peptides (X being any amino acid)..
